# Structure and function of IWS1 in transcription elongation

**DOI:** 10.1101/2025.08.28.672863

**Authors:** Della Syau, Felix Steinruecke, Sophie Roth, Ernst Schmid, Karen Adelman, Johannes Walter, Lucas Farnung

**Affiliations:** Department of Cell Biology, Blavatnik Institute, Harvard Medical School, Boston, MA, USA; Department of Biological Chemistry & Molecular Pharmacology, Blavatnik Institute, Harvard Medical School, Boston, MA, USA; The Eli and Edythe L. Broad Institute, Cambridge MA 02142, USA; Ludwig Center at Harvard, Boston, MA 02115, USA; Howard Hughes Medical Institute, Boston, MA 02115, USA

**Author notes:** Correspondence should be addressed to L.F.

## Abstract

Transcription elongation by RNA polymerase II is a tightly regulated process that requires coordinated interactions between transcription elongation factors. IWS1 (Interacts with SPT6) has been implicated as a core elongation factor, but its molecular role remains unclear. We show that the intrinsically disordered C-terminal region of IWS1 contains short linear motifs (SLiMs) that multivalently engage the elongation machinery. Using cryo-electron microscopy, we map SLiMs in IWS1 that interact with Pol II subunits RPB1, RPB2, and RPB5, as well as elongation factors DSIF, SPT6, and ELOF1. Functional assays demonstrate that distinct IWS1 SLiMs specify IWS1 recruitment and IWS1-dependent transcription stimulation. IWS1 recruitment to the transcription elongation complex depends on association via the RPB1 jaw and binding of downstream DNA. Transcription elongation stimulation requires interactions with the RPB2 lobe and ELOF1. We identify other transcription elongation factors including ELOA and RECQL5 that bind the RPB1 jaw and demonstrate that IWS1 protects the activated transcription elongation complex from RECQL5 inhibition. We also reveal the binding of the histone reader and IWS1 interactor LEDGF to a transcribed downstream nucleosome. Our findings establish IWS1 as a modular scaffold that helps organize the transcription elongation complex, illustrating how disordered regions regulate transcription elongation.

## Introduction

Transcription elongation by RNA polymerase II (Pol II) requires the coordinated action of elongation factors that collectively maintain the enzyme in a processive, chromatin-compatible state. Rather than acting individually, elongation factors form extensive interaction networks on Pol II. Despite structural and functional advances in elucidating the architecture of elongation complexes, the principles by which individual elongation factors assemble onto Pol II to form the productive transcription elongation machinery remain incompletely understood.

IWS1 (Interacts with SPT6) is a highly conserved Pol II– associated elongation factor required for normal gene expression from yeast to mammals (1-7). Genetic ablation of IWS1 in mice causes early embryonic lethality (8,9), and depletion of IWS1 in human cells leads to Pol II accumulation at promoters with loss of transcription across gene bodies (2), underscoring its essential role in productive elongation. Beyond transcription itself, IWS1 has been implicated in cotranscriptional RNA processing (10) and chromatin regulation (11-17).

Structurally, IWS1 contains a central helical bundle formed by a HEAT repeat and a TFIIS N-terminal domain (TND) that directly bind to the transcription elongation complex (6,18). The central HEAT/TND domain is flanked by long disordered N- and C-terminal regions (Figure 1A). The N-terminal region binds histones and contains well-characterized TND-interacting motifs (TIMs) that bind transcription elongation factors such as TFIIS, ELOA, and LEDGF (2,13,19). Current structural and functional knowledge of IWS1, however, is heavily skewed toward its N-terminal intrinsically disordered region (IDR) and the structured HEAT/TND core (6,18,20-22). In contrast, the C-terminal tail is highly conserved and known to interact with DNA (13), yet no structural elements or binding motifs have been assigned. The persistence of strong evolutionary conservation in the apparently disordered C-terminal tail (Figure 1A) suggests it may encode important short linear motifs (SLiMs) for protein interactions. IDRs often serve as interaction hubs through such SLiMs, and the unexplained conservation of the IWS1 C-terminus raises the possibility that critical, yet undiscovered, interactions reside in the C-terminal tail. Addressing this knowledge gap is essential for a complete understanding of how IWS1 functions as a core transcription elongation factor.

**Figure 1.**
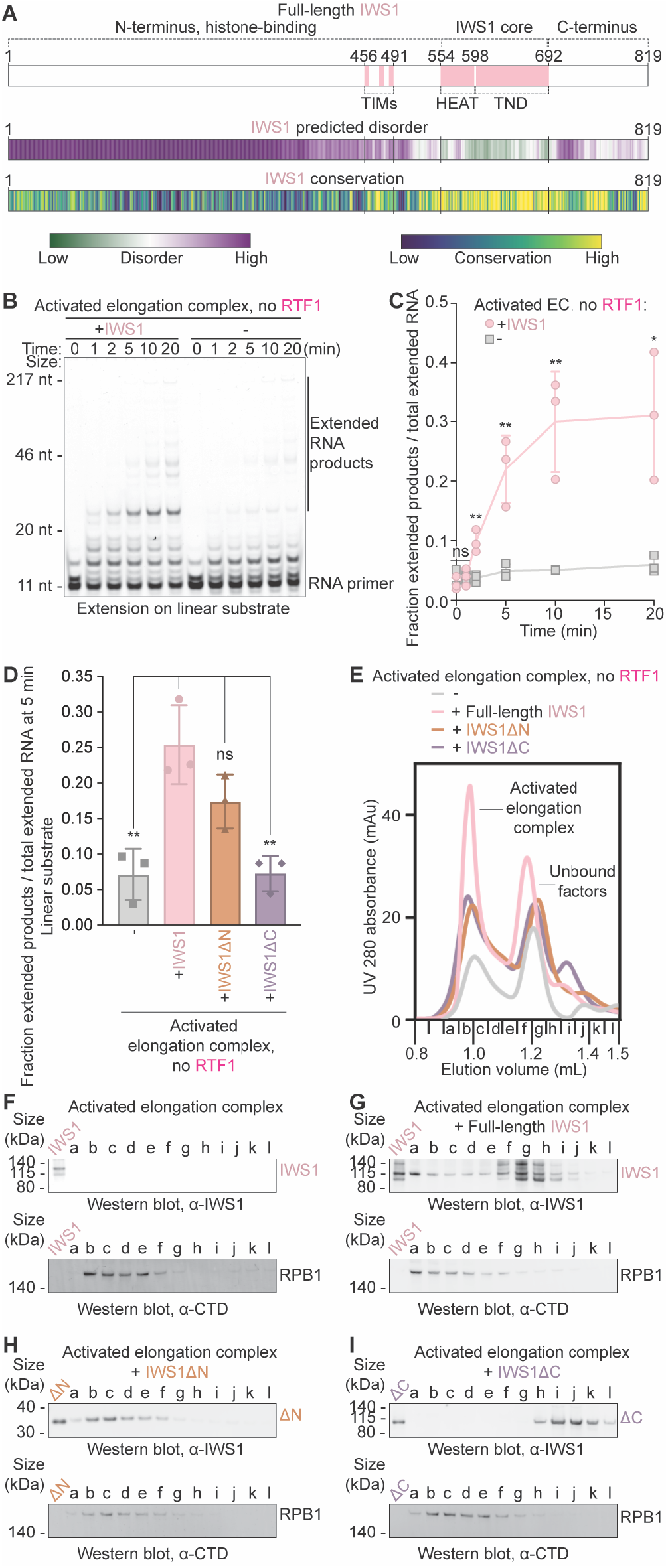
IWS1 stimulates transcription elongation and requires its C-terminus for elongation complex binding. **(A)** Domain architecture of IWS1 with known elements labeled. Predicted disorder (IUpred score) and conservation (ConSurf score) for each residue indicated by color scheme. **(B)** Time-course RNA extension assay without RTF1 and with or without IWS1 on linear DNA. Samples were separated by denaturing gel electrophoresis. RNA extension was monitored using FAM dye on RNA. Initial RNA and extended products are marked. Results are representative of at least three independent experiments. **(C)** Quantification of (B) from n=3 independent experiments. * p<0.05, ** p<0.01, ns = not significant relative to samples of the same timepoint with no IWS1 using unpaired t-test. **(D)** Quantification of n=3 independent RNA extension assays without RTF1 and without IWS1, with full-length IWS1, with IWS1ΔN (551--819), or with IWS1ΔC (1-699) on linear DNA, quenched after 5 minutes after addition of CTP, GTP, and UTP (10 µM). Representative gel image shown in Figure 4L, * p<0.05, ** p<0.01, ns = not significant relative to +IWS1 using unpaired t-test. **(E)** UV 280 nm size exclusion chromatogram of elongation complexes assembled without IWS1, with full-length IWS1, with IWS1ΔN (551-819), or with IWS1ΔC (1-699). RTF1 and TFIIS were excluded in complex formation. Peaks corresponding to activated elongation complex or unbound factors indicated. **(F-I)** Western blots of size exclusion fractions in (E) for activated elongation complex assembled (F) without IWS1, (G) with full-length IWS1, (H) with IWS1ΔN (551-819), and (I) with IWS1ΔC (1-699). Activated EC = Pol II, DSIF, SPT6, PAF1c, TFIIS, ELOF1. Individual data points are shown for C and D. Columns represent mean values across n = 3 independent experiments and error bars represent standard deviations in C and D.

Using functional assays and single-particle cryo-electron microscopy (cryo-EM), we show that IWS1 directly stimulates transcription elongation, and we identify that the far C-terminus of IWS1 is constitutively required for binding the elongation complex. We find that IWS1 is an integral component of activated transcription elongation complexes and we identify previously uncharacterized SLiMs within the IWS1 C-terminus which interact with Pol II subunits RPB1, RPB2, and elongation factors DSIF, SPT6, and ELOF1. Structure-function analyses show that mutational disruption of individual IWS1 SLiMs impairs transcription elongation *in vitro*. This work establishes IWS1’s C-terminal tail as a previously unrecognized scaffold that organizes the Pol II elongation complex, illustrating a general principle whereby conserved but often disordered regions of elongation factors orchestrate the assembly and processivity of the transcription elongation apparatus. Together, our findings represent the most complete structure of a transcription elongation complex to date and highlight the broader significance of intrinsically disordered protein segments in the regulation of gene expression.

## Results

### IWS1 stimulates transcription elongation

To assess the role of IWS1 in transcription elongation, we first used a fully reconstituted transcription elongation system to perform RNA extension assays with porcine RNA polymerase II (Pol II) and purified human transcription elongation factors (DSIF, SPT6, PAF1 complex including RTF1 [PAF1c], TFIIS, ELOF1, P-TEFb, and IWS1) on a nucleosomal DNA substrate (Figure S1 and S2A, Methods). We observed that addition of IWS1 increased the fraction of full-length transcripts in a concentration-dependent manner (Figures S2B-C). To precisely evaluate the contribution of IWS1, we next omitted the strong transcription elongation stimulator RTF1 from our assay. While RTF1 strongly stimulates transcription both *in vitro* and in cells (23,24), it is not required for the formation of the activated elongation complex (23,25-31). As expected, we observed substantially less transcription in the absence of RTF1 (Figures 1B-C). Strikingly, addition of full-length IWS1 in reactions lacking RTF1, however, led to a strong increase in extended RNA products. Thus, IWS1 directly stimulates transcription elongation (Figures 1B-C).

To further test the effects of IWS1 on transcription elongation complexes on diverse DNA substrates, we performed time-course assays on linear, mono-nucleosomal, and dinucleosomal DNA substrates. Indeed, IWS1 increased fulllength transcripts 2-3.5–fold on all substrates even in the presence of RTF1 (Figure S2). Additionally, IWS1 facilitated polymerase passage through nucleosome entry barriers (Figure S2K). Together, these findings establish IWS1 as a transcription elongation factor that directly promotes Pol II elongation on naked DNA and chromatinized templates.

We next addressed whether the IWS1 N- and C-terminal regions are required for transcription stimulation. Deletion of the intrinsically disordered N-terminus (IWS1 ΔN, Figure 1A) only showed a minor reduction in IWS1’s stimulatory effect on elongation for linear and mono-nucleosomal DNA substrates (Figure 1D, S2D-I). In contrast, deletion of the IWS1 C-terminus (IWS1 ΔC, Figure 1A), which contains a patchwork of conserved and non-conserved, disordered regions, fully abrogated the stimulatory effect on elongation for linear, mono-nucleosomal, and dinucleosomal DNA substrates (Figure 1D, S2D-L). Unexpectedly, these results suggested that the C-terminal tail of IWS1 is critical for promoting transcription elongation, whereas the histone-binding N-terminal domain contributes only minimally.

To determine whether the loss of stimulation in the presence of IWS1 ΔC was due to impaired association with the elongation machinery, we performed size exclusion chromatography experiments with reconstituted elongation complexes (Methods). Due to the similar molecular weight of IWS1 ΔC and RTF1 (Figure S1A), we again omitted RTF1 from our reactions to unambiguously determine the presence of IWS1 and its truncation mutants in the complex. We ran SDS-PAGE and Western blots of the chromatography fractions to assess incorporation of IWS1 into the activated elongation complex. We first confirmed that the activated elongation complex assembled with full-length IWS1 in the absence of RTF1, shown by the detection of both IWS1 and RPB1 C-terminal domain (CTD) signal in the fractions corresponding to the earliest chromatogram peak (Figures 1E-G, S3A-B). Whereas IWS1 ΔN co-eluted with the elongation complex similarly to full-length IWS1 (Figures 1H, S3C), IWS1 ΔC failed to co-elute with the complex (Figures 1I, S3D), suggesting a complete loss of association in the absence of the IWS1 C-terminus. Accordingly, IWS1 HEAT/TND (a truncation lacking the IWS1 N- and C-termini) also did not associate with the elongation complex (Figure S3E-F). We conclude that the C-terminal region, but not the N-terminus, is required for IWS1 binding to the transcription elongation complex and that the structured HEAT/TND alone is not sufficient to support IWS1 recruitment to the transcription elongation complex.

### Cryo-EM structure of the activated transcription elongation complex with ELOF1 and IWS1

Because the intrinsically disordered C-terminal tail of IWS1 lacked structural domains but showed high levels of amino acid sequence conservation (Figure 1A), we asked whether it harbored short linear motifs (SLiMs) that interact with the elongation machinery. We prepared a cryo-EM sample containing Pol II, the kinase P-TEFb, ELOF1, IWS1, and human elongation factors DSIF, PAF1c, and SPT6 on a mono-nucleosomal substrate (Figure S4, Methods). Of note, we included LEDGF in the preparation because of its known interaction with IWS1 and nucleosomes (19,32-34). Pol II was positioned in front of the nucleosome and transcribed up to 4 base pairs (bp) in front of the nucleosome upon addition of NTPs (Figure S1C, S4). We performed size exclusion chromatography to select for the formed complex containing IWS1, mildly cross-linked relevant fractions using glutaraldehyde, and prepared the sample for single-particle cryo-EM characterization (Figure S4B, Methods).

Analysis and classification of the cryo-EM data resulted in a reconstruction of the Pol II-DSIF-SPT6-PAF1c-TFIISIWS1-ELOF1-LEDGF transcription elongation complex at a nominal resolution of 2.4 Å (FSC 0.143) from 762,523 particles (Figure 2B, Table 1, Video 1). The complex is bound to a partially unwrapped downstream nucleosome resolved at a lower resolution (see below). Further classification and masked refinements resulted in local reconstructions of different parts of the elongation complex at resolutions between 2.4 Å and 7.1 Å (Figures S5-S6, Methods). A pseudo-atomic model was built into the overall reconstruction with the aid of the locally refined cryo-EM maps and an extensive AlphaFold-Multimer prediction screen (Figure S5-S7, Methods, Table 2). The structure shows good stereochemistry (Figures S5-S6, Table 1).

**Figure 2.**
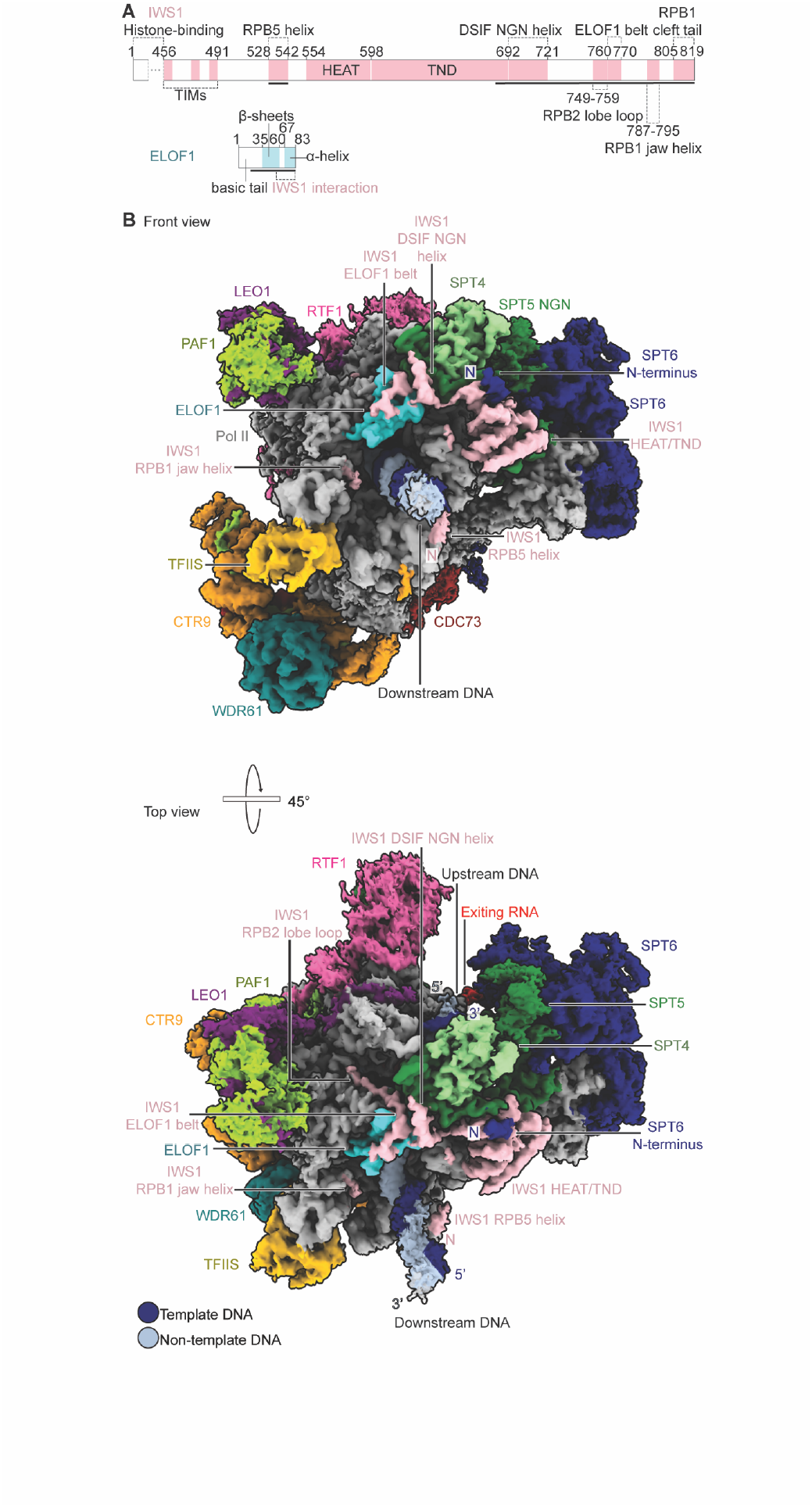
Structure of the RNA polymerase II-DSIF-SPT6-PAF1c-TFIIS-IWS1-ELOF1 transcription elongation complex. **(A)** Domain architecture of IWS1 and ELOF1. Modelled regions are indicated as solid black lines. **(B)** Two views of the Pol II-DSIF-SPT6-PAF1c-TFIIS-IWS1-ELOF1 transcription elongation complex on composite map T generated from Frankenmap. Color code used throughout.

### Architecture of the activated transcription elongation complex with IWS1 and ELOF1

The resulting structure of the Pol II-DSIF-SPT6-PAF1c-TFIIS-IWS1-ELOF1-LEDGF-nucleosome complex positions the polymerase 9 base pairs (bp) in front of the nucleosome (bp -9) with five nucleotides of backtracked RNA (Figure S8B-C). The complex had an architecture similar to mammalian elongation complex structures lacking ELOF1 and IWS1 (22,23,35,36). In addition to revealing four SLiM-like interactions confined to 10–20-residue long segments in the IWS1 C-terminal region with the elongation machinery (see below), it revealed the following new features. First, we observed a ∼40 Å shift of the PAF1-LEO1 dimerization domain towards the RPB2 lobe (Figure S8M). This shift likely reflects the conformational flexibility of the PAF1-LEO1 module previously observed across different structures (23), with the presence of IWS1 and ELOF1 potentially favoring a conformation proximal to the RPB2 lobe (Figure S8M). Second, we visualized a C-terminal helix of LEO1 (residues 528-561) next to the upstream DNA and SPT4 (Figure S8N). Third, we observed the CDC73 Ras-like domain bound to the RPB1 foot next to the SPT6 tSH2 domain (Figure S8O) (21). Fourth, our structure positions the zinc finger of TFIIS domain III within the Pol II funnel, likely favored by the backtracked configuration that we observed in the Pol II active site and funnel (Figure S8B,C,P). Finally, we visualized ELOF1. As observed in transcription-coupled nucleotide excision repair (TC-NER) complexes (37,38), ELOF1 binds the lobe of RPB2 (Figure S8Q). In our complex, ELOF1 is positioned next to the SPT5 NGN domain and contacts the IWS1 C-terminus (Figure S8Q, see below). We visualized additional residues of the ELOF1 N-terminus (residues 17-19) that stretch along the Pol II clamp towards the transcription bubble (Figure S8R). Similar to structural observations of Elf1 in yeast (21,39,40), ELOF1 is an integral component of the activated elongation complex in mammals. Together, this structure provides the most complete overview of the mammalian transcription elongation complex yet.

### IWS1 structure and interactions with the activated elongation complex

In our structure, IWS1 interacted extensively with the transcription elongation machinery and stabilized transcription elongation factors including ELOF1 and SPT5. First, we observed a density positioned against RPB5 next to the downstream DNA (Figures 3C, S8S). Due to the absence of this helix in structures without IWS1 (23,36) and the high interaction score of IWS1 and RPB5 in an AlphaFold-Multimer prediction screen (Methods, Figure S7A), we docked the predicted IWS1 helix (residues 528-542) onto this density. Next, we fitted and refined the IWS1 HEAT/TND (residues 554-692). The HEAT/TND is positioned at the RPB1 clamp head above the SPT5 KOW2-3 domains as visualized in other activated transcription elongation complexes containing IWS1 or the yeast homolog Spn1 (Figures 3D, S8T) (21,22,41).

**Figure 3.**
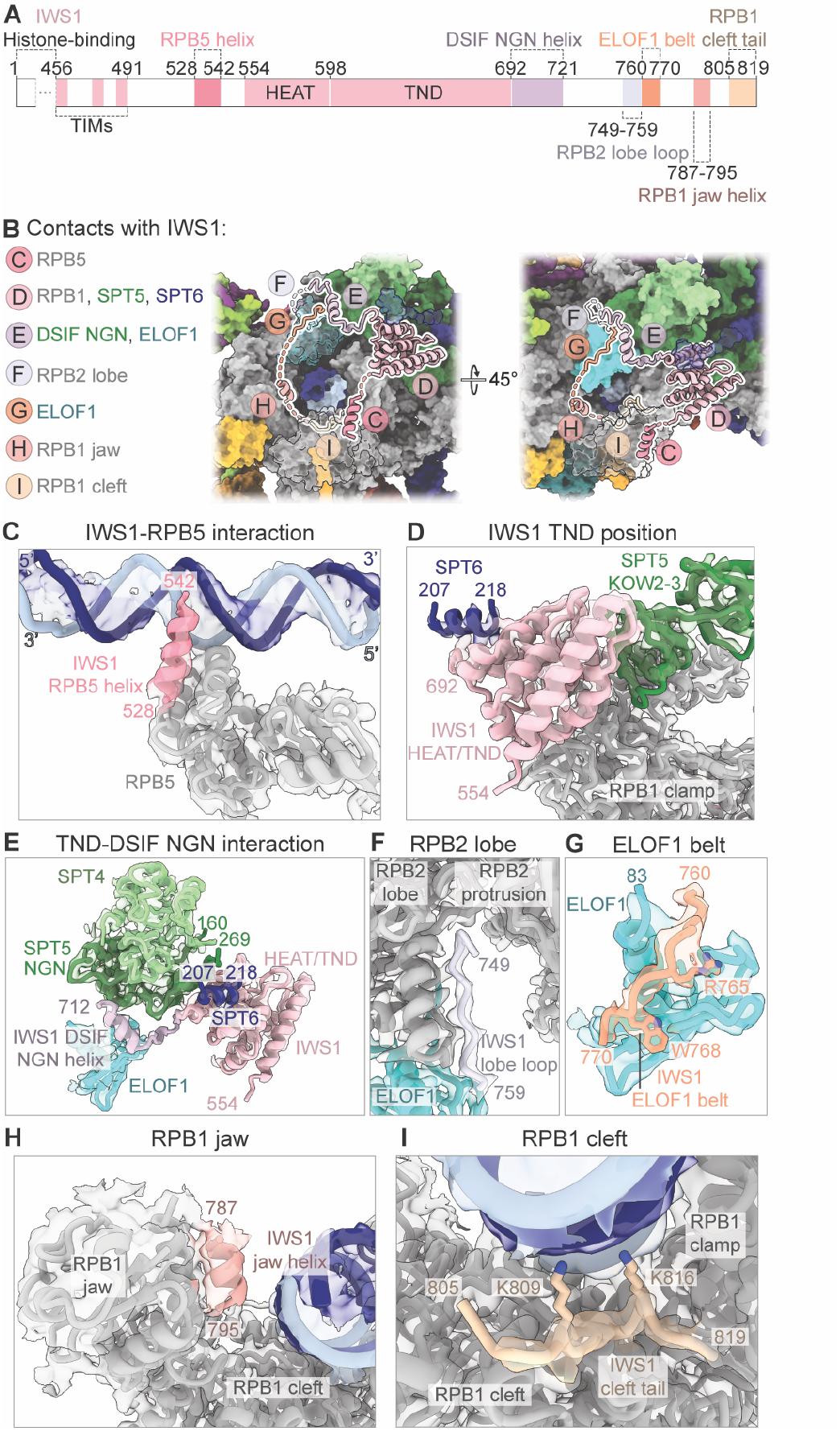
IWS1 interaction interfaces with RNA polymerase II and transcription elongation factors. **(A)** Domain architecture diagram of IWS1 with boundaries of newly modeled SLiMs indicated. Color code used throughout. **(B)** Schematic overview of IWS1 contacts with transcription elongation complex. **(C)** IWS1 interaction with RPB5 overlaid on map I. **(D)** Position of IWS1 HEAT/TND and interactions with SPT5 and SPT6 overlaid on map B. **(E)** IWS1 interaction with DSIF NGN and ELOF1 overlaid on map B. **(F)** IWS1 interaction with RPB2 lobe overlaid on map B. **(G)** IWS1 interaction with ELOF1 overlaid on map B. Residues of IWS1 ELOF1 belt with clear side-chain densities shown as sticks. **(H)** IWS1 interaction with RPB1 jaw overlaid on map C. **(I)** IWS1 interaction with RPB1 cleft overlaid on map C. Positively charged residues shown as sticks.

Similar to structural studies of Spn1 (18,20,21), we observed a helix from the SPT6 N-terminus (residues 207-230) that bound to the IWS1 TND (Figures 3D, S8T, S9B). Notably, casein kinase 2 was assumed to be important for mediating the SPT6-IWS1 interaction (42), but our complex formation that only used P-TEFb for phosphorylation indicates that casein kinase 2 is not constitutively required. Additionally, in proximity to the known SPT6-IWS1 interaction, we found an SPT5 helix (SPT5 residues 160-170) that contacts the IWS1 TND and the SPT6 N-terminus (Figures 3E, S8U). In the past (22), we likely did not observe these interactions between SPT6, SPT5, and IWS1 in the mammalian elongation complex due to lack of sufficient resolution or absence of ELOF1.

Surprisingly, we observed multiple IWS1 SLiMs along the IWS1 C-terminus (residues 693-819) that interact with (i) the SPT5 NGN domain, (ii) the RPB2 lobe, (iii) ELOF1, and (iv) the RPB1 jaw. Residues of IWS1 (residues 693-721) immediately after the HEAT/TND and part of the IWS1 C-terminus (IWS1 “DSIF NGN helix”) wrap around the DSIF NGN domain, mediating contacts between the DSIF NGN, IWS1, and ELOF1 (Figures 3E, S8U). IWS1 subsequently inserts into the region between the RPB2 protrusion and the RPB2 lobe above the non-template strand (IWS1 “RPB2 lobe loop,” residues 749-759, Figures 3F, S8V).

From the RPB2 lobe, IWS1 residues 760-770 wrap around the solvent-exposed surface of ELOF1 (Figures 3G, S8W). We named this segment the “ELOF1 belt” of IWS1. Our AlphaFold-Multimer predictions recapitulated these findings (Figure S9A). The extensive interaction surface of more than 2800 Å^2^ between IWS1, ELOF1, and DSIF may stabilize the binding of DSIF and ELOF1. Indeed, SPT4 and the SPT5 NGN domain are more stably anchored to the elongation complex compared to previous structures (23,43) (Figure S9C). This could also stabilize the observed interactions between the SPT5 and SPT6 N-terminal helices and the IWS1 HEAT/TND.

Finally, we fitted the most C-terminal helix of IWS1 (residues 787-795, “RPB1 jaw helix”) into a density between the RPB1 jaw and the downstream DNA (Figures 3H, S8X). AlphaFold-Multimer also predicted that the very C-terminus of IWS1 could meander into a crevice formed between the downstream DNA, the RPB1 clamp, the RPB1 cleft, and RPB5 (Figure S7D). We were able to assign backbone and some sidechain densities of IWS1 residues 805-819 to this interaction and named it the IWS1 “RPB1 cleft tail” (Figures 3I, S8Y). These contacts between the extreme C-terminus of IWS1, RPB1, and RPB5 position positively charged residues of IWS1 (residues R787, K790, R793, K794, K809, K816) near the downstream DNA backbone (Figure S9D). Consequently, we observed that truncating the IWS1 C-terminus led to an overall reduction in IWS1 DNA binding using a fluorescence polarization assay (Methods, Figure S9E).

Importantly, the IWS1 C-terminal SLiM interactions are conserved (Figures S9F-I, S10A-D). AlphaFold-Multimer predictions of *S. cerevisiae* Rpb1-Spn1, Rpb2-Elf1-Spn1, Rpb5-Spn1, Spt4-Spt5-Spn1, and Spt6-Spn1 captured similar interactions involving the Spn1 C-terminus (Figure S9F-I). Notably, the DSIF NGN helix and RPB1 cleft tail are not conserved in our predicted Spn1 interactions (Figures S9F-I, S10).

In summary, we observe that IWS1, ELOF1, SPT6, and DSIF form an interconnected network, favored by highly specific interactions of the IWS1 C-terminal SLiMs that contribute to enhanced stability of the elongation complex, rationalizing why IWS1 is a core transcription elongation factor.

### The extreme C-terminus of IWS1 mediates association with the elongation complex

To better understand which of the newly identified C-terminal interactions are required for IWS1 recruitment to the elongation complex, we generated a series of C-terminal IWS1 truncations (Figure 4A) that removed the entire C-terminus downstream of the DSIF NGN helix (IWS1 1-721), the ELOF1 belt and extreme C-terminus (IWS1 1-759), or only the extreme C-terminus containing the RPB1 jaw helix and the RPB1 cleft tail (IWS1 1-786). Using size exclusion chromatography, we then reconstituted elongation complexes with these truncations and ran SDS-PAGE and Western blots of the eluted fractions. We could not detect any IWS1 signal co-eluting with the RPB1 CTD for the larger C-terminal truncations (IWS1 1-721, IWS1 1-759), showing that they did not associate with the elongation complex (Figures 4B-F, S11B-C). We could not detect any signal for the most minimal truncation (IWS1 1-786) on SDS-PAGE (Figure S11A), and it exhibited a very faint IWS1 signal in the Western blot (Figure 4D). Hence, loss of the RPB1 jaw helix and interaction with the downstream DNA in the Pol II cleft severely compromised IWS1 recruitment and association with the transcription elongation complex. As expected, RNA extension assays showed that none of the C-terminal truncations stimulated transcription elongation as IWS1 recruitment is impacted in these mutants (Figure 4L-M).

**Figure 4.**
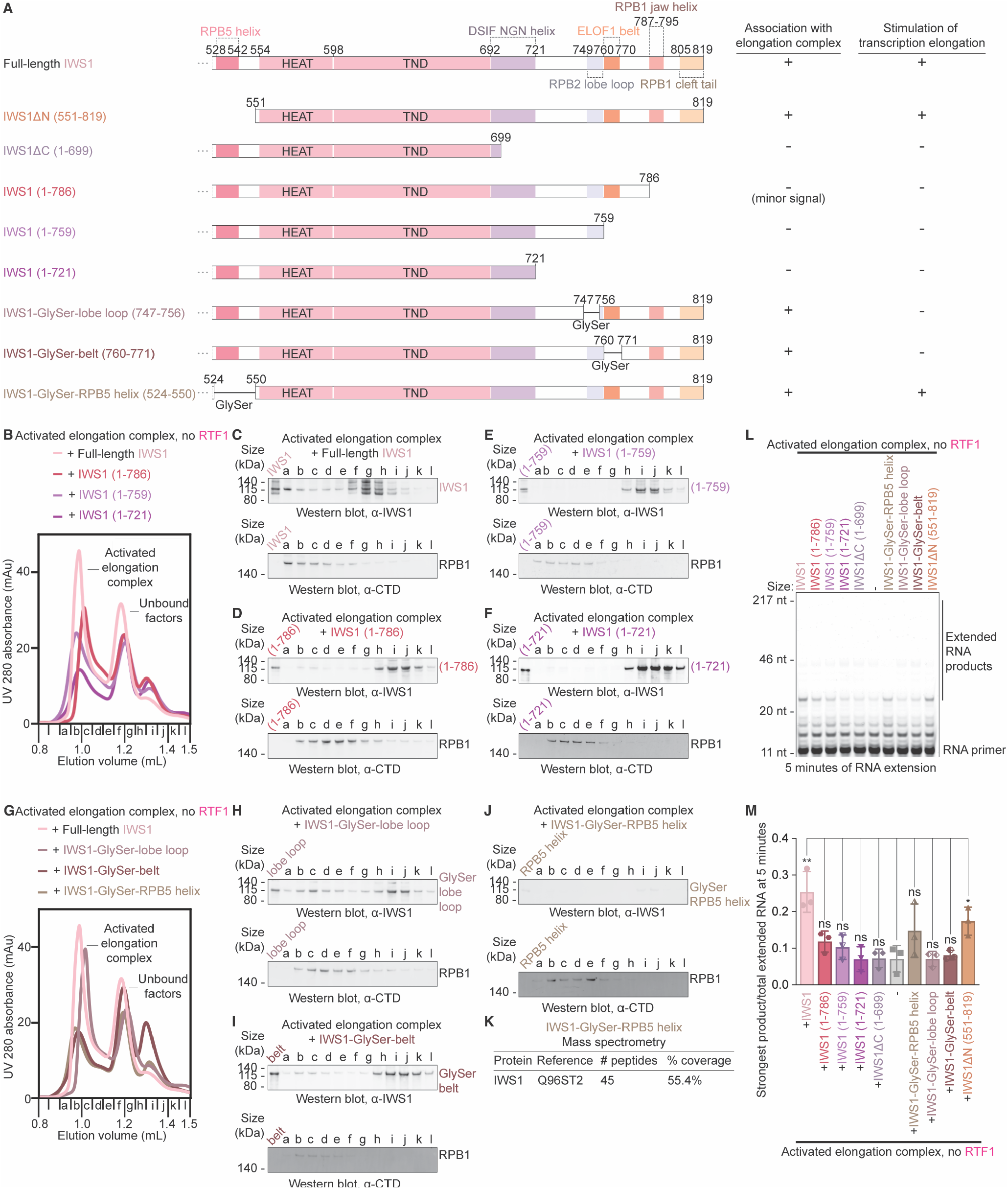
Roles of IWS1 C-terminal SLiMs in association with elongation complex and stimulation of transcription elongation. **(A)** Domain architecture of IWS1 and its mutants. GlySer = GGGGS repeats replacing the indicated region of IWS1. For IWS1-GlySer-lobe loop, residues 747-756 replaced with GGGSGGGGS. For IWS1-GlySer-belt, residues 760-771 replaced with GGGGSGGGGS. For IWS1-GlySer-RPB5 helix, residues 524-550 replaced with GGGGSGGGGSGGGGSGGGGSGGGGS. Table summarizes properties of elongation complex association and transcription elongation stimulation for each construct. **(B)** UV 280 nm size exclusion chromatograms of elongation complexes assembled with full-length IWS1, IWS1 (1-786), IWS1 (1-759), or IWS1 (1-721). RTF1 and TFIIS were excluded in complex formation. Peaks corresponding to activated elongation complex or unbound factors indicated. Chromatogram of elongation complex with full-length IWS1 was previously shown in Figure 1E and included here for comparison. **(C)** Western blots of size exclusion fractions from full-length IWS1 assembly previously shown in Figure 1G shown again as a reference. **(D-F)** Western blots of size exclusion fractions in (B) for activated elongation complex assembled with (D) IWS1 (1-786), (E) IWS1 (1-759), and (F) IWS1 (1-721). **(G)** UV 280 nm size exclusion chromatograms of elongation complexes assembled with full-length IWS1, IWS1-GlySer-lobe loop, IWS1-GlySer-belt, or IWS1-GlySer-RPB5 helix. RTF1 and TFIIS were excluded in complex formation. Peaks corresponding to activated elongation complex or unbound factors indicated. Chromatogram of elongation complex with full-length IWS1 was previously shown in Figure 1E and included here for comparison. **(H-J)** Western blots of size exclusion fractions in (G) for activated elongation complex assembled with (H) IWS1-GlySer-lobe loop, (I) IWS1-GlySer-belt, and (J) IWS1-GlySer-RPB5 helix. **(K)** Mass spectrometry peptide coverage of purified IWS1-GlySer-RPB5 helix. **(L)** RNA extension assay without RTF1 with IWS1 mutants on linear DNA. Reactions were quenched 5 min after addition of CTP, GTP, and UTP (10 µM). Results are representative of at least three independent experiments. **(M)** Quantification of (L) from n = 3 independent experiments. Individual data points shown and error bars represent standard deviations. Columns represent mean values across n = 3 experiments. * p<0.05, ** p<0.01, ns = not significant relative to -IWS1 using unpaired t-test. Activated elongation complex = Pol II, DSIF, SPT6, PAF1c, TFIIS, ELOF1.

### Specific regions in the C-terminus of IWS1 stimulate transcription elongation

To further investigate how the other IWS1 SLiMs influence IWS1-elongation complex binding or transcription activity, we generated separate glycine-serine mutations of the RPB2 lobe loop (IWS1 residues 747-756) and the ELOF1 belt (IWS1 residues 760-771), where the mutant was designed to encode GlyGlyGlyGlySer repeats of equivalent length to the segment of IWS1 being replaced. We also generated a glycine-serine mutation of the IWS1 RPB5 interacting helix (IWS1 residues 524-550) (Figure 4A). The IWS1-GlySer-lobe loop and IWS1-GlySer-belt mutants stably associated with the elongation complex, consistent with our findings that the IWS1 RPB1 jaw helix and cleft tail interactions determine elongation complex association (Figures 4G-I, S11D-E). We found that the IWS1-GlySer-RPB5 helix mutant could not be detected by the IWS1 antibody in Western blotting (Figure 4J) despite showing incorporation into the activated elongation complex in SDS-PAGE (Figure S11F). Mass spectrometry verified that IWS1 is present (Figures 4K, S11G). Despite stable elongation complex association, RNA extension levels of IWS1-GlySer-lobe loop and IWS1-GlySer-belt were significantly compromised (Figure 4L-M). IWS1-GlySer-RPB5 helix, however, showed a mild, but not statistically significant, increase in transcription stimulation compared to no IWS1. These mutational studies of IWS1 show that the disruption of individual SLiM interactions within the IWS1 C-terminus are detrimental for efficient transcription elongation, even if IWS1 is stably bound to the elongation complex. Interactions of IWS1 with ELOF1 and the RPB2 lobe are required for direct stimulation of transcription elongation.

In addition to disrupting IWS1 SLiM interactions by mutating the SLiMs on IWS1, we identified specific residues on ELOF1 that interact with IWS1 and mutated these residues to alanine (ELOF1_beltmut_) (Figure 5A-B). Similar to our results with the IWS1-GlySer-belt mutant, reconstituted elongation complexes containing ELOF1_beltmut_ showed binding of wildtype IWS1 (Figure 5C-D) but complexes containing ELOF1_beltmut_ and wildtype IWS1 did not stimulate RNA extension relative to reactions without IWS1 but containing wildtype ELOF1 (Figure 5E-F). Together, our size exclusion chromatography and RNA extension assays demonstrate that recruitment of IWS1 to the transcription elongation complex and transcription stimulation by IWS1 involve interactions of distinct IWS1 regions that have non-overlapping functions.

**Figure 5.**
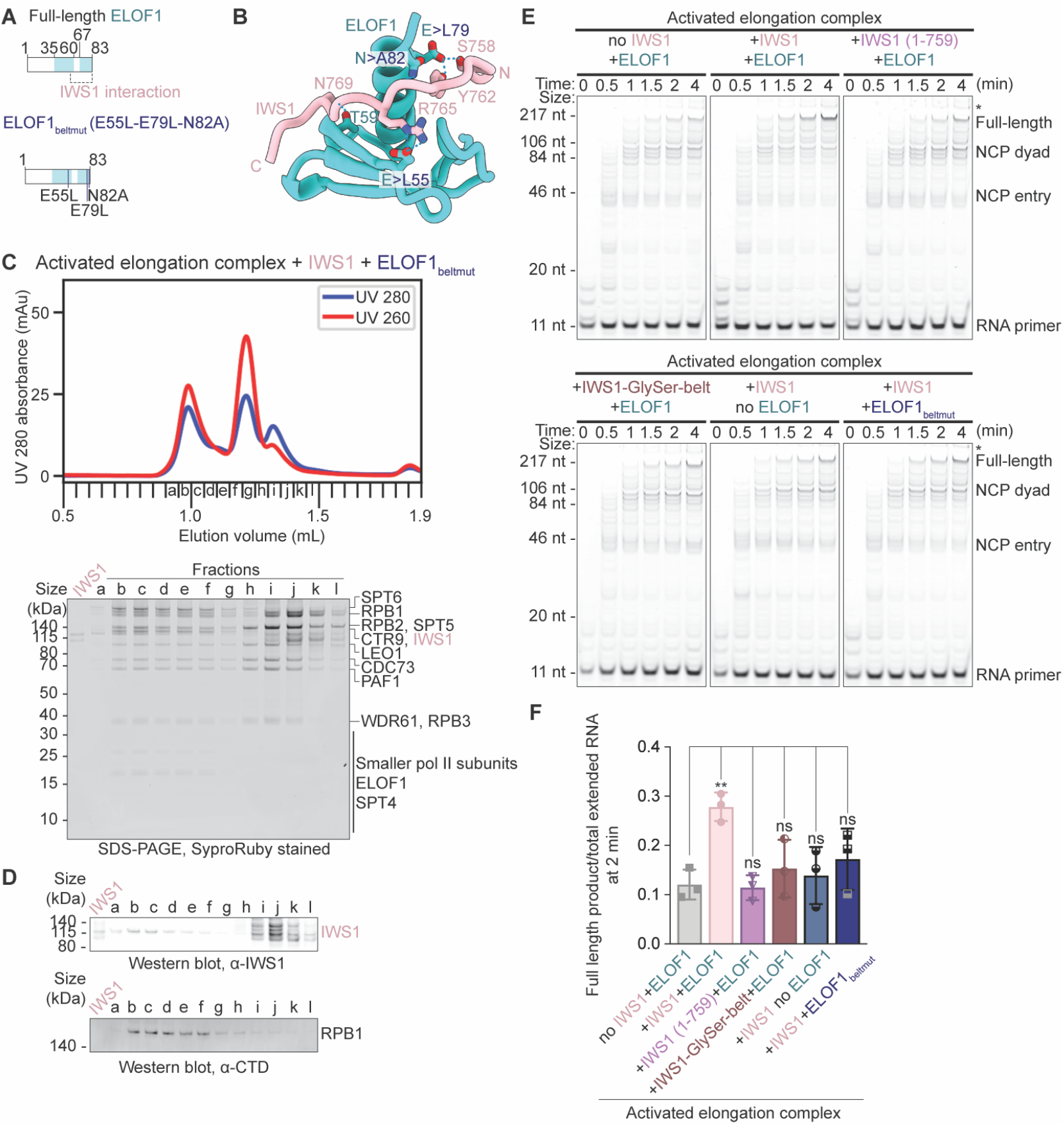
IWS1-ELOF1 interaction is required for transcription stimulation. **(A)** Domain architecture of ELOF1 and ELOF1_beltmut_. **(B)** Interaction between IWS1 ELOF1 belt and ELOF1. Residues mutated in ELOF1_beltmut_ indicated. Interacting residues shown as sticks and hydrogen bonds shown as dotted lines. **(C)** UV 280 nm and UV 260 nm traces for size exclusion chromatograms and corresponding SDS-PAGE of elongation complexes assembled with full-length IWS1 and ELOF1_beltmut_. Complex does not contain RTF1 or TFIIS. Fractions were loaded onto 4-12% Bis-Tris SDS-PAGE gels, run in 1x MES buffer, and stained with SyproRuby. **(D)** Western blots of size exclusion fractions in (C) for activated elongation complex assembled with full-length IWS1 and ELOF1_beltmut_. **(E)** RNA extension assays of IWS1 mutants with or without ELOF1 or ELOF1_beltmut_ on mono-nucleosome. Samples were separated by denaturing gel electrophoresis. RNA extension was monitored using FAM dye on RNA. Initial RNA and extended products are marked. Results are representative of at least three independent experiments. Not fully denatured RNA marked with asterisk (*) on gels. **(F)** Quantification of (E) from n = 3 independent experiments. Individual data points are shown. Activated elongation complex = Pol II, DSIF, SPT6, PAF1c including RTF1, TFIIS. Columns represent mean values across n = 3 independent experiments and error bars represent standard deviations. * p<0.05, ** p<0.01, ns = not significant relative to -IWS1 +ELOF1 using unpaired t-test.

### Multiple elongation factors bind an overlapping surface on the RPB1 jaw

The extreme C-terminus of IWS1 is important for IWS1 association with the elongation complex. Importantly, the IWS1 RPB1 jaw binding site overlaps with the RPB1 jaw binding sites of TC-NER factor UVSSA (37,38), Pol II chaperone RPAP2 (44), and TFIIH subunit XPB (45) (Figure S12B). This suggests that the RPB1 jaw interaction could represent an important regulatory surface that is mutually exclusive to the binding of a wider variety of protein factors in transcription initiation and elongation.

To identify additional RPB1 jaw binders, we performed an AlphaFold-Multimer pairwise interaction screen containing human RPB1 and key transcription initiation and elongation factors (2) (Figures 6A, S12A). We ranked the interactions by their cSPOC score (46) (Methods). Using this approach, we identified multiple high-confidence interactors of RPB1 (Figures 6A, S12A). We further categorized the high-confidence interactors based on whether they are predicted to interact with the RPB1 jaw. These specific interactors included the transcription elongation factor Elongin A (ELOA1) and the transcription-related helicase RECQL5 (47-49) (Figure S12C). Specifically, a C-terminal helix in ELOA1 (residues 756-772) is predicted to bind the RPB1 jaw but has not been characterized. To structurally validate this prediction, we analyzed existing cryo-EM data of an Elongin ABC-containing transcription elongation complex where the C-terminal helix of ELOA1 was not modeled (50). Upon low-pass filtering the map (Methods), we observed a helical density positioned next to the RPB1 jaw which overlapped with the predicted binding site of the ELOA1 C-terminal helix (Figure 6B).

**Figure 6.**
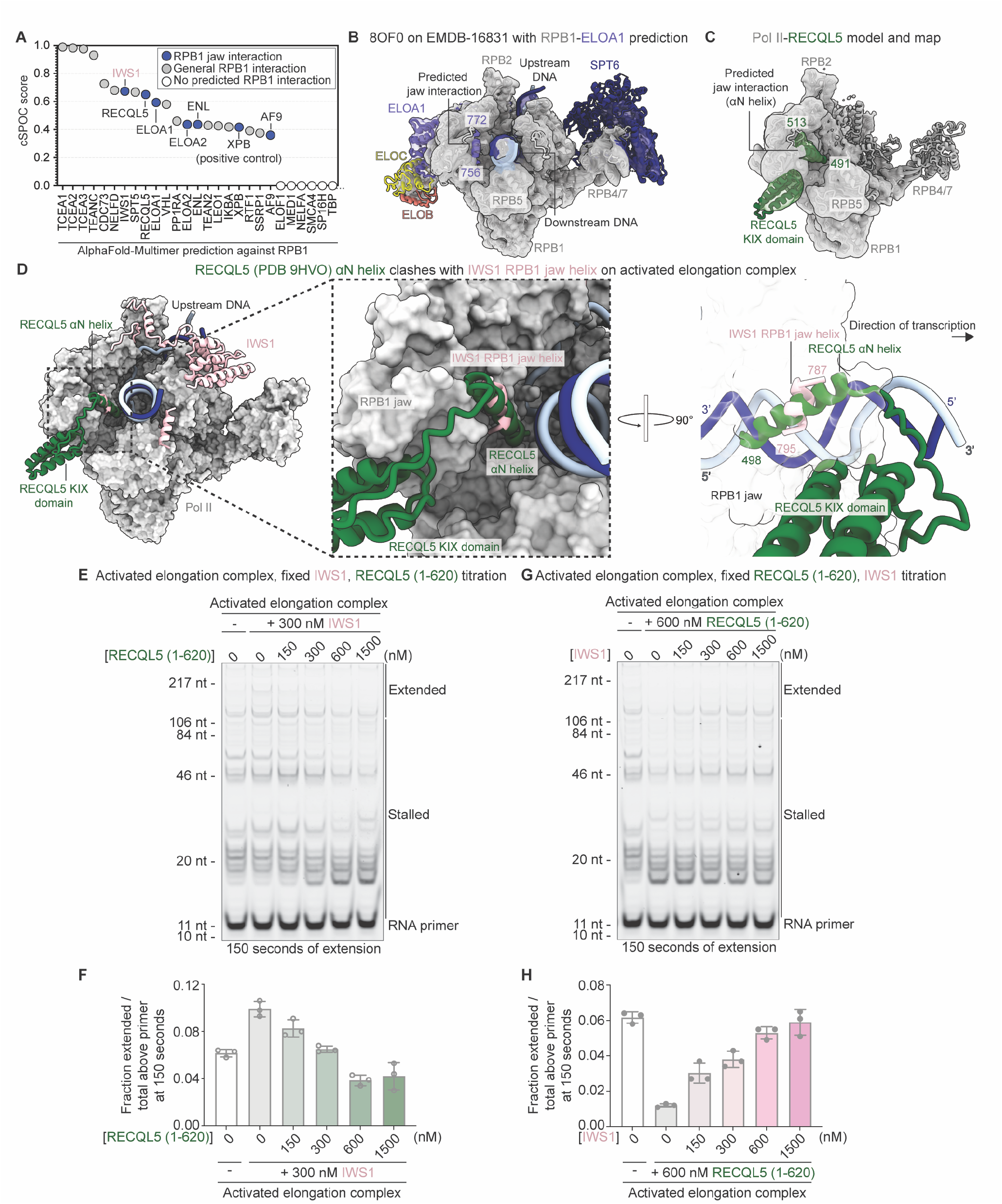
A protein-protein interaction screen identifies a surface of the RPB1 jaw as a binding platform for elongation factors. **(A)** Interaction screen using AlphaFold-Multimer. Predictions ranked by cSPOC score. Representative RPB1 jaw interactors are shown as dark blue circles, general RPB1 interactions shown as grey circles, and no predicted RPB1 interactions shown as white circles. **(B)** Fit of predicted helical element of ELOA1 into Pol II–ELOABC elongation complex model (PDB 8OF0) and Coulomb potential map (EMDB-16831) confirms interaction of ELOA1 C-terminal helix and RPB1 jaw. **(C)** Fit of RECQL5-RPB1 protein-protein interaction prediction into Pol II–RECQL5 complex Coulomb potential map (this study) confirms interaction of RECQL5 helix and RPB1 jaw. RECQL5 KIX domain is also resolved. **(D)** Overlay of RECQL5 αN helix and KIX domain from RECQL5-Pol II model (PDB 9HVO) on activated elongation complex model (this study) shows binding overlap of RECQL5 αN helix and IWS1 RPB1 jaw helix. Pol II model shown as surfaces and IWS1 model shown as cartoons. **(E)** RNA extension assay on linear substrate with increasing concentrations of RECQL5 (1-620) in the presence of 300 nM IWS1. The 0 nM IWS1 and 0 nM RECQL5 reaction is included as a reference. Reactions were quenched 150 sec after addition of ATP, CTP, GTP, and UTP (100 µM). Results are representative of three independent experiments. **(F)** Quantification of (E) from n = 3 independent experiments. Repeated measures one-way ANOVA was performed for the conditions containing 300 nM IWS1, where conditions containing 150 nM, 300 nM, 600 nM, and 1500 nM RECQL5 were compared to the mean of the 0 nM RECQL5 condition. Resulting p-value 0.016. **(G)** RNA extension assay on linear substrate with increasing concentrations of IWS1 in the presence of 600 nM RECQL5. The 0 nM IWS1 and 0 nM RECQL5 reaction is included as a reference. Reactions were quenched 150 sec after addition of ATP, CTP, GTP, and UTP (100 µM). Results are representative of three independent experiments. **(H)** Quantification of (G) from n = 3 independent experiments. Repeated measures one-way ANOVA was performed for the conditions containing 600 nM RECQL5, where conditions containing 150 nM, 300 nM, 600 nM, and 1500 nM IWS1 were compared to the mean of the 0 nM IWS1 condition. Resulting p-value 0.016. Individual data points are shown, columns represent mean values across n = 3 independent experiments, and error bars represent standard deviations for (F) and (H). Activated elongation complex = Pol II, DSIF, SPT6, PAF1c excluding RTF1, ELOF1.

Our AlphaFold-Multimer screen also predicted that the αN “brake” helix of RECQL5 (residues 491-513) (47-49) binds to the RPB1 jaw (Figure S12C). We prepared a cryo-EM sample of a Pol II complex with RECQL5 (Figure S13A-F) and visualized the interaction between the RECQL5 αN brake helix and the RPB1 jaw, along with the RECQL5 KIX domain and the RPB1 jaw (Figure 6C). The interaction between the RECQL5 αN brake helix and the RPB1 jaw has also been observed in two recent publications at high-resolution (47,48).

Although RECQL5 was shown to bind to an elongation complex (48), its interaction with the RPB1 jaw would clash with the IWS1 binding site (Figures 6D, S13G). To test whether RECQL5 and IWS1 could compete for transcription elongation complex binding through their jaw domain interactions, we prepared RECQL5 (residues 1-620) (Figure S13H) to include in RNA extension assays (Methods). We were able to recapitulate a transcriptional stalling effect (48,49) by increasing concentrations of RECQL5 in reactions with a fixed concentration of IWS1 (Figure 6E-F). Notably, the stalling effect was reversed when we titrated increasing concentrations of IWS1 in reactions with a fixed concentration of RECQL5 (Figure 6G-H), indicating that IWS1 and RECQL5 compete for binding to the transcription elongation complex.

### LEDGF binds the distal side of the transcribed nucleosome

We also analyzed the cryo-EM densities corresponding to the nucleosomal substrate and associated factors (Figure 7).

**Figure 7.**
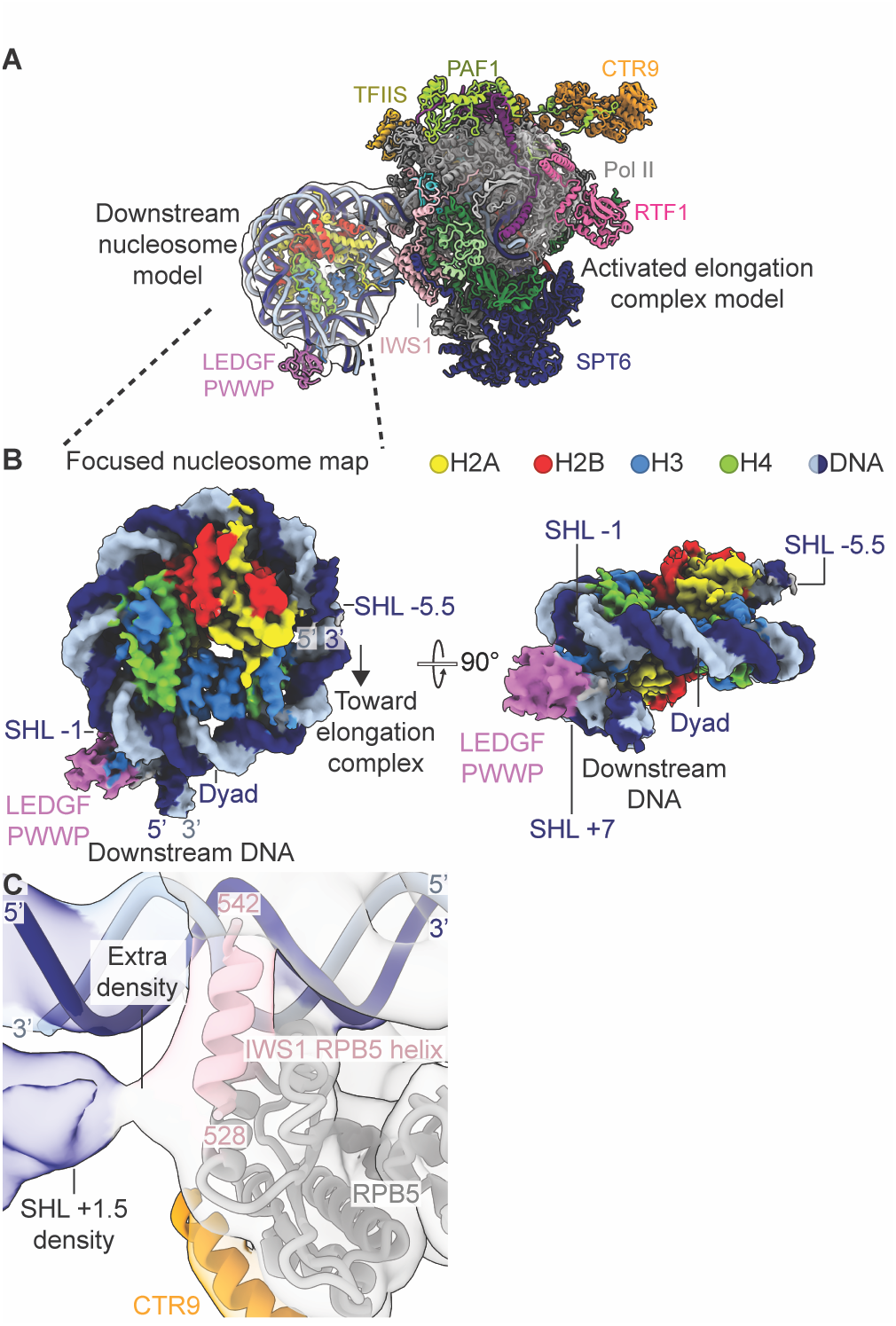
Interaction details of LEDGF PWWP and IWS1 with transcribed nucleosome. **(A)** Cryo-EM density of the nucleosome model positioned with respect to the activated elongation complex atomic model using map R. Nucleosome and activated elongation complex models shown as cartoons. **(B)** Cryo-EM density of the nucleosome substrate (map Q). The transcribed nucleosome is unwrapped to SHL -5.5. LEDGF (orchid) binds the H3 tail at the promoter-distal side of the transcribed nucleosome and contacts both DNA gyres. **(C)** The IWS1 N-terminus reaches toward SHL +1.5 of the transcribed nucleosome after its interaction with downstream DNA and RPB5. Activated elongation complex model overlaid on map I. SHL = super helical location.

Using particle subtraction to keep the signal for the 10-subunit core Pol II and nucleosome (Figure S5, Methods), we obtained focused maps of the nucleosome (7.6 Å) and 10-subunit Pol II (8.7 Å). These maps allowed us to position the nucleosome model relative to the activated elongation complex (Figure 7A). We obtained a 3.5 Å map of the nucleo-some using particle subtraction and focused refinements of the same particle set (Figure 7B). The nucleosomal DNA is unwrapped to super helical location (SHL) -5.5, exposing the promoter-proximal H2A/H2B dimer (Figures 7B, S8B-D).

We observed binding of the LEDGF PWWP domain only on the promoter-distal side of the nucleosome because LEDGF requires interactions with two DNA gyres to engage the methylated H3 tail (Figure 7B) (33,34). The observed LEDGF-nucleosome interaction is consistent with previous observations of LEDGF binding to mono-nucleosomes (33,34). The IWS1 N-terminal region harbors histone chaperone activity and the IWS1 interaction with RPB5 positions it near the partially unwrapped nucleosome. Accordingly, we observed density of the IWS1 N-terminus projecting towards SHL +1.5 of the nucleosome (Figure 7C). This likely positions IWS1 to engage histones upon further unwrapping of nucleosomal DNA. We did not observe a direct interaction between LEDGF and IWS1, likely because this involves TIM3 of IWS1 (residues 482-491) and the TND of LEDGF (residues 342-426) (2,51), both of which were not visualized in our reconstructions.

## Discussion

Here we report the structure of the mammalian activated elongation complex in the presence of transcription elongation factors ELOF1 and IWS1 and demonstrate that IWS1 stimulates transcription elongation. We identify that the IWS1 C-terminus makes extensive SLiM-like interactions with multiple core components of the elongation complex, including Pol II subunits RPB1, RPB2, RPB5, downstream DNA, ELOF1, DSIF, and SPT6 (Figure 3). Disruption of these interactions through C-terminal IWS1 truncations, mutations of the C-terminal SLiMs to GlySer repeats, or targeted mutations on ELOF1 reduces the stimulatory effect of IWS1 and disfavors the formation of activated elongation complexes (Figures 4 and 5). As we observe a stabilization of the DSIF NGN domain in the cryo-EM density when compared to a complex formed in the absence of IWS1 and ELOF1 (Figure S9C), this network of interactions may function to stabilize elongation factor binding and maintain Pol II in an elongation state. Accordingly, single-molecule studies of elongation factor binding dynamics show that yeast Spn1 (human IWS1), Spt4/5 (human DSIF), Spt6 (human SPT6), and Elf1 (human ELOF1) are stably associated with Pol II and have the longest residence times during transcription elongation (52,53). These similar binding dynamics may be enhanced by the stabilization of elongation factors on the polymerase by IWS1.

Recently, Zheenbekova and colleagues (41) observed that deleting residues 786-819 from IWS1 (corresponding to the RPB1 jaw helix and RPB1 cleft tail interacting regions) led to a loss of transcription stimulation on a mononucleosomal substrate. They proposed that residues 786-819 of IWS1 restricts movement of the downstream DNA and observed that ELOF1 may be important for positioning this region of IWS1. Complexes without ELOF1 did not show stable cryo-EM density of the RPB1 jaw helix and cleft tail. Their findings could be interpreted in such a way where the RPB1 jaw helix and cleft tail stimulate transcription by positioning the downstream DNA, and without these elements, transcription stimulation no longer occurs. Our study agrees that deletion of the RPB1 jaw helix and cleft tail abrogates the transcription stimulation activity of IWS1 (Figure 4M, IWS1 1-786). However, our size exclusion chromatography experiments (Figures 4B-D, S11A) demonstrate that the loss of IWS1 transcription stimulation activity from this far C-terminal deletion is due to a severe reduction of IWS1-elongation complex association.

Importantly, we identify that the IWS1 ELOF1 belt and the IWS1 RPB2 lobe loop function in transcription stimulation (Figures 4M, 5F), demonstrating that IWS1 requires the coordinated interactions involving all its C-terminal motifs to stimulate transcription, not just its extreme C-terminus. Our Spn1 AlphaFold-Multimer predictions (Figure S9) and IWS1 C-terminal sequence alignments (Figure S10) reinforce these findings, as the RPB1 jaw helix, RPB2 lobe loop, and ELOF1 belt are well-conserved, but yeast Spn1 does not appear to contain a conserved interaction with the RPB1 cleft. In fact, this region is lacking in yeast Spn1. Additionally, RTF1 and IWS1 appear to stimulate elongation in an additive manner that may suggest different mechanisms, consistent with genetic evidence in yeast (5,12) where a Spn1 HEAT/TND mutation (K192N) or the Spn1 HEAT/TND expressed on its own exacerbates phenotypes of RTF1 deletion. We noticed that progressive C-terminal truncations of IWS1 showed a graded effect on IWS1 transcription stimulation (Figure 4L-M). Although the C-terminal truncations lack the RPB1 jaw binding region and therefore could not stably associate with the elongation complex, it is possible that limited interactions between elongation factors and remaining IWS1 C-terminal SLiMs allowed for transient association and consequently, transcription stimulation. This would affirm our interpretation that all IWS1 C-terminal SLiMs are required for transcription stimulation, and the presence of more possible contacts improves elongation complex stability and therefore transcription rate.

Notably, IWS1 ΔN (551-819) showed a significant reduction of transcription stimulation activity relative to fulllength IWS1 on a dinucleosomal substrate that was not detected on linear or mono-nucleosomal substrates (Figure S2J-L). A similar effect was observed with an IWS1 N-terminal truncation (T4, 550-819) (41), that reduced transcription stimulation activity relative to full-length IWS1 on a mono-nucleosomal substrate in the presence of higher NTP concentrations (0.4 mM). Since IWS1 has a role in chromatin transcription and the N-terminus can bind histones (11-17), this suggests that the IWS1 N- and C-termini serve distinct but complementary roles in chromatin transcription. The histone-binding activity of the N-terminus may be crucial for Pol II traversal through complex chromatin substrates, whereas the stabilizing property of the C-terminus generally increases transcription efficiency.

Structural and *in silico* prediction data of RPB1 jaw interactors show that IWS1, UVSSA, RPAP2, XPB, RECQL5, ELOA1, and others bind an overlapping region on the RPB1 jaw. We show that a fully assembled activated elongation complex with IWS1 and ELOF1 protects the transcription elongation complex from inhibition by RECQL5 (Figure 6).

Together, these findings imply a regulatory mechanism where different elongation factors may compete for RPB1 jaw binding depending on the transcriptional state or cellular context. It is likely that RECQL5 association requires the partial disassembly of the activated elongation complex by the dissociation of at least IWS1, suggesting a more complex mechanism of factor exchange than previously proposed (48). Furthermore, the interaction stability of DSIF with the transcription machinery plays an important role in the initial recruitment of the ATPase CSB to the transcription machinery during TC-NER (54). Destabilization and displacement of the SPT5 NGN domain and SPT4 is a prerequisite for CSB ATPase engagement. Because IWS1 and ELOF1 have a stabilizing effect on DSIF, IWS1 must also be displaced or rearranged during TC-NER. Accordingly, IWS1 is not compatible with the binding of multiple TC-NER factors including UVSSA, CSA, CSB, and STK19 (Figure S14A). Comparison of our activated elongation complex containing IWS1 and ELOF1 suggests that the positioning of IWS1 and ELOF1 are, however, compatible with the paused elongation complex (Figure S14B) (55,56).

We could not resolve a map with a clear downstream nucleosome bound by the elongation complex, which is likely related to the flexibility of the downstream DNA, suggesting that a nucleosome substrate with an alternative pause site could improve resolution of the polymerase-nucleosome interface (36). Although we observed additional density N-terminal to the RPB5 helix on IWS1 (Figure 7C), an alternative substrate with a more stably positioned nucleosome could potentially resolve IWS1-histone contacts.

Our focused processing of the nucleosome density shows that the LEDGF PWWP binds to the distal H3K36me3 site when the polymerase is positioned at bp -9 (Figure 7B). We did not observe any PWWP densities at a second nucleosome binding site reported previously (34), and we also did not observe density for the LEDGF integrase binding domain (IBD) even though it could interact with IWS1, SPT6, SPT5, or PAF1, all of which were present in our complex (2,32,57,58). Additional structural studies where the polymerase is transcribed to different stall sites within the nucleosome, or with the polymerase bound to an upstream transcribed nucleosome, could address whether other co-transcriptional PWWP binding sites are possible, whether LEDGF interacts with an upstream H3K36me3-modified nucleosome (22), and whether the LEDGF IBD interactions are maintained during transcription.

Together, we provide the most complete model of the mammalian transcription elongation complex to date and show a modular, multivalent engagement of IWS1 that integrates structural flexibility with specific molecular recognition, revealing a mechanistic principle of elongation factor function.

### Limitations of the study

While we and others have shown that SPT6/Spt6 (22,59) and Spn1 N-termini (13) can bind to histones, the polymerase may need to unravel more nucleosomal DNA by transcribing further into the nucleosome for this binding to occur.

The positioning of the IWS1 RPB5 interacting helix at the downstream DNA may allow the disordered IWS1 N-terminus to engage the exposed histones during transcription. How this occurs in concert with SPT6 and/or the histone chaperone FACT remains to be elucidated (15,21,22,59). Although our nucleosome substrate for our cryo-EM structure contains a T-less cassette up to nucleosome position bp -4, we observed that our complex is backtracked to bp -9 (Figure S8B-C). The stable wrapping of the DNA in our nucleosome density suggests that the nucleosome acts as a barrier to the elongation complex (43). It is possible that the polymerase adopts a backtracked state due to this barrier.

The dataset for our RNA polymerase II-DSIF-SPT6-PAF1c-TFIIS-IWS1-ELOF1-LEDGF-nucleosome complex was collected on a sample crosslinked with glutaraldehyde, which could favor certain stabilized states of the elongation complex. It does not necessarily represent all possible conformational states and assemblies.

We also observed a major shift of the PAF1-LEO1 dimer on the Pol II surface. We were unable to directly attribute this architectural change to the presence of ELOF1 and IWS1, but we note that previous structures (23,35) also showed positional heterogeneity of the PAF1-LEO1 dimer. Allosteric changes introduced by IWS1 and ELOF1 could favor the observed positioning on top of RPB2.

## Acknowledgements

We thank all members of the Farnung lab for discussions. We thank J. W. Markert for support with data processing. We thank The Harvard Cryo-EM Center for Structural Biology at Harvard Medical School for support with data collection. We thank the Taplin Mass Spectrometry Facility at Harvard Medical School for support with mass spectrometry analysis. We thank S.M. Vos for critical reading and input. This work was supported by the Smith Family Awards Program for Excellence in Biomedical Research to L.F.; the Damon Runyon Rachleff Innovator Award Program to L.F.; the National Institutes of Health [DP2-ES036404 to L.F., HL098316 to J.C.W.; R01GM139960 to K.A.]; the Boehringer Ingelheim Fonds to S.R.; and the National Science Foundation [DGE 2140743 to F.S. and E.W.S.]. J.C.W. is a Howard Hughes Medical Institute Investigator. L.F. is a Freeman Hrabowski Scholar of the Howard Hughes Medical Institute. This paper was typeset with the bioRxiv word template by @Chrelli: www.github.com/chrelli/bioRxiv-word-template

## Author contributions

D.S. conducted all biochemical experiments unless stated otherwise. D.S. prepared the IWS1 complex for cryo-EM. D.S. collected and analyzed IWS1 cryo-EM data. F.S. prepared RECQL5, the RECQL5-RNA polymerase II complex, and conducted cryo-EM analysis of RECQL5. S.R. prepared the RECQL5 (residues 1-620) protein. D.S. and S.R. purified RNA polymerase II. D.S., J.C.W, K.A., and L.F. conceived the transcription elongation *in silico* prediction screen. E.S. conducted the *in silico* prediction screen and provided analysis. L.F. designed and supervised research. D.S. and L.F. wrote the manuscript with input from all authors.

## Data availability

Materials are available from Lucas Farnung upon request and a completed materials transfer agreement with Harvard Medical School and HHMI.

### Declaration of Generative AI and AI-assisted technologies in the writing process

During the preparation of this work the authors used ChatGPT in order to revise text for clarity and conciseness. After using this tool/service, the authors reviewed and edited the content as needed and take full responsibility for the content of the publication.

### Quantification and statistical analysis

No statistical methods were used to predetermine sample size. The experiments were not randomized, and the investigators were not blinded to allocation during experiments and outcome assessment.

## Materials and Methods

### Cell culture

Sf9 cells (Expression Systems, Cat# 94-001S), *T*.*ni* (Hi5) cells (Expression Systems, Cat# 94-002S), and Sf21 cells (Expression Systems, Cat# 94-003S) were cultured in ESF 921 insect cell culture medium (Expression Systems, Cat# 96-001-01) at 27°C for baculovirus production and protein expression, unless otherwise indicated. *E. coli* DH5α (New England Biolabs (NEB) Cat# C2987H), DH10EMBacY (Geneva Biotech), and BL21-CodonPlus (DE3)-RIL (Agilent Cat# 230245) were cultured in LB broth (EMD-Millipore Cat# 71753-5) at 37°C for plasmid production and protein expression, unless otherwise indicated.

### Cloning

The following sequence containing the TspRI cut site and the modified Widom 601 sequence with an A-less cassette to nucleosomal bp -4 was synthesized and inserted into a pIDTSmart-Kan vector (IDT): 5’-ACG AAG CGT AGC ATC ACT GTC TTG TGT TTG GTG TGT CTG GGT GGT GGC AAT TTT CGA TGT ATA TAT CTG ACA CGT GCC TGG AGA CTA GGG AGT AAT CCC CTT GGC GGT TAA AAC GCG GGG GAC AGC GCG TAC GTG CGT TTA AGC GGT GCT AGA GCT GTC TAC GAC CAA TTG AGC GGC CTC GGC ACC GGG ATT CTG ATA TCG CGC GTG ATC TTA CGG

CAT TAT ACG TA-3’ (-4 vector) using the same approaches as previously described (36). The bp +64 vector and bp +115 vector were generated as previously described (22,36).

The dinucleosome vector was generated by assembling the reverse Widom 601 sequence and the Widom 601 sequence with a 15 bp linker between the two sequences: 5’-GAT CAG ACT CGA CTA-3’ into a pIDTSmart-Kan vector (IDT).

The H3K36C histone H3 variant was generated by introducing the lysine-to-cysteine mutation to a vector encoding X. laevis H3 C110A using ‘Round-The-Horn cloning (60).

Plasmid constructs for *H. sapiens* DSIF, *H. sapiens* SPT6, *H. sapiens* PAF1 complex, *H. sapiens* RTF1, *H. sapiens* TFIIS, *H. sapiens* IWS1, *H. sapiens* LEDGF, and *H. sapiens* PTEF-b were previously generated (22,33,36). A gene-optimized *H. sapiens* ELOF1 gBlock was synthesized and cloned into a 1-B vector using ligation-independent cloning (LIC). Similarly, a gene-optimized *H. sapiens* ELOF1 gBlock containing E55L, E79L, and N82A mutations was synthesized and cloned into a 1-B vector using LIC.

IWS1 (1-786), IWS1 (1-759), IWS1 (1-721), IWS1ΔC (1-699), IWS1ΔN (551-819), and IWS1 HEAT/TND (551-699) were generated using ‘Round-The-Horn cloning. IWS1-GlySer-loop (747-756), IWS1-GlySer-belt (760-771), and IWS1-GlySer-RPB5helix (524-550) were generated by amplifying the IWS1 438-C vector backbone followed by Gibson assembly with gBlocks (IDT) encoding GlyGlyGlyGlySer repeats of equivalent length to the segment of IWS1 being replaced.

RECQL5 was amplified from cDNA and cloned into the 438-C vector using LIC. RECQL5 (1-620) was amplified from the full-length RECQL5 vector and cloned into the 1-C vector using LIC.

### Protein expression

*H. sapiens* DSIF, *H. sapiens* TFIIS, and X. laevis histones were expressed in *E. coli. H. sapiens* SPT6, *H. sapiens* PAF1 complex, *H. sapiens* RTF1, *H. sapiens* IWS1 and mutants, *H. sapiens* LEDGF, and *H. sapiens* PTEF-b were expressed in Hi5 insect cells.

*H. sapiens* ELOF1 and ELOF1_beltmut_ were expressed in BL21 (DE3) RIL cells at 37 °C until an OD of 0.6 at 600 nm was reached. Expression was induced with the addition of 1 mM IPTG and cells were grown at 18 °C overnight. Cells were centrifuged and pellets were resuspended in lysis buffer (20 mM Na·HEPES pH 7.4, 300 mM NaCl, 30 mM imidazole pH 8.0, 10% (v/v) glycerol, 5 mM b-mercaptoethanol, 0.284 µg mL–1 leupeptin, 1.37 µg mL–1 pepstatin A, 0.17 mg mL–1 PMSF, and 0.33 mg mL–1 benzamidine), then snap-frozen in liquid nitrogen and stored at -80 °C until purification.

### Protein purification

S. scrofa RNA polymerase II, *H. sapiens* DSIF, *H. sapiens* SPT6, *H. sapiens* RTF1, *H. sapiens* TFIIS, *H. sapiens* IWS1, *H. sapiens* LEDGF, and *H. sapiens* PTEF-b were purified as previously described (22,23,33,35,56). X. laevis histones were purified as previously described (61).

*H. sapiens* ELOF1 with an N-terminal 6xHis tag and TEV protease cleavage site was purified at 4°C. Cells expressing ELOF1 were resuspended in lysis buffer and subsequently lysed by sonication. The lysate was centrifuged and cleared by ultra-centrifugation. The supernatant containing ELOF1 was subsequently filtered using 5 mm and 0.45 mm syringe filters. The filtered supernatant was applied to a HisTrap HP 5 mL (Cytiva). The column was subsequently washed with 5 CV lysis buffer, 5 CV high salt buffer (20 mM Na·HEPES pH 7.4, 1000 mM NaCl, 30 mM imidazole pH 8.0, 10% (v/v) glycerol, 5 mM b-mercaptoethanol), and 5 CV lysis buffer. ELOF1 was eluted from the HisTrap column using a 10 CV 0-100% gradient of nickel elution buffer (20 mM Na·HEPES pH 7.4, 300 mM NaCl, 500 mM imidazole pH 8.0, 10% (v/v) glycerol, 5 mM b-mercaptoethanol). The elution was fractionated and analyzed using SDS-PAGE. Fractions containing ELOF1 were pooled and applied to 3.5k MWCO SnakeSkin dialysis tubing (Thermo Scientific) in lysis buffer at 4°C overnight. 1.5 mg of TEV protease was added to the sample prior to dialysis to remove the N-terminal His6 tag. The dialyzed sample was applied to a HisTrap HP 5 mL, pre-equilibrated in lysis buffer. The flow-through containing ELOF1 was collected and subsequently concentrated using an Amicon 3,000 MWCO centrifugal filter unit (Millipore). The concentrated sample was applied to a HiLoad 16/600 Superdex 75 pg (Cytiva), equilibrated in gel filtration buffer (20 mM Na·HEPES pH 7.4, 300 mM NaCl, 10% (v/v) glycerol, 1 mM TCEP). The elution was fractionated and analyzed by SDS-PAGE. Sample containing ELOF1 was concentrated using an Amicon 3,000 MWCO centrifugal filter unit (Millipore).

*H. sapiens* PAF1 complex containing untagged PAF1, LEO1, CDC73, WDR61, and CTR9 with a C-terminal MBP, 6xHis tag, and TEV protease cleavage site was purified at 4°C. Cells expressing the PAF1 complex were resuspended in lysis buffer and lysed by douncing using a B-type douncer. The lysate was centrifuged and cleared by ultra-centrifugation. The supernatant containing the PAF1 complex was subsequently filtered using 5 mm and 0.45 mm syringe filters. The filtered supernatant was applied to a HisTrap HP 5 mL (Cytiva). The column was subsequently washed with 9 CV lysis buffer. A self-packed XK column (Cytiva) with 15 mL of Amylose resin (New England Biolabs) was attached to the HisTrap column. PAF1 complex was eluted from the HisTrap column using nickel elution buffer. The HisTrap column was removed, and the amylose column was washed with 5 CV lysis buffer. PAF1 complex was eluted from the amylose column with 5 CV amylose elution buffer (20 mM Na·HEPES pH 7.4, 300 mM NaCl, 30 mM imidazole pH 8.0, 116.9 mM maltose, 10% (v/v) glycerol, 5 mM b-mercaptoethanol). The elution was fractionated and analyzed using SDS-PAGE. Fractions containing PAF1 complex were pooled and applied to 3.5k MWCO SnakeSkin dialysis tubing (Thermo Scientific) in dialysis buffer (20 mM Na·HEPES pH 7.4, 150 mM NaCl, 30 mM imidazole pH 8.0, 10% (v/v) glycerol, 5 mM b-mercaptoethanol) at 4°C overnight. 1.5 mg of TEV protease was added to the sample prior to dialysis to remove the C-terminal MBP-His6 tag. The dialyzed sample was applied to a HiTrap Q HP 5 mL (Cytiva), pre-equilibrated in dialysis buffer. The column was subsequently washed with 10 CV dialysis buffer. PAF1 complex was eluted from the HiTrap Q column with a 10 CV 0-100% gradient of high salt buffer (20 mM Na·HEPES pH 7.4, 1000 mM NaCl, 30 mM imidazole pH 8.0, 10% (v/v) glycerol, 5 mM b-mercaptoethanol). The elution was fractionated and analyzed by SDS-PAGE. Sample containing the PAF1 complex was collected and subsequently concentrated using an Amicon 30,000 MWCO centrifugal filter unit (Millipore). The concentrated sample was applied to a Superose 6 Increase 10/300 GL (Cytiva), equilibrated in gel filtration buffer. The elution was fractionated and analyzed by SDS-PAGE. Sample containing the PAF1 complex was concentrated using an Amicon 30,000 MWCO centrifugal filter unit (Millipore).

*H. sapiens* IWS1 and IWS1 constructs (1-786), (1-759), (1-721), ΔC (1-699), GlySer-lobe loop, GlySer-belt, and GlySer-RPB5 helix were purified as described for full-length IWS1 (22).

*H. sapiens* IWS1 HEAT/TND (551-699) with an N-terminal 6xHis-MBP tag and TEV protease cleavage site was purified at 4°C. Cells expressing IWS1 HEAT/TND were resuspended in lysis buffer and subsequently lysed by sonication. The lysate was centrifuged and cleared by ultra-centrifugation. The supernatant containing IWS1 HEAT/TND was subsequently filtered using 5 mm and 0.45 mm syringe filters. The filtered supernatant was applied to a HisTrap HP 5 mL (Cytiva). The column was subsequently washed with 9 CV lysis buffer. A self-packed XK column (Cytiva) with 15 mL of Amylose resin (New England Biolabs) was attached to the HisTrap column. IWS1 HEAT/TND was eluted from the HisTrap column using nickel elution buffer. The HisTrap column was removed, and the amylose column was washed with 5 CV lysis buffer. IWS1 HEAT/TND was eluted from the amylose column with 5 CV amylose elution buffer (20 mM Na·HEPES pH 7.4, 300 mM NaCl, 30 mM imidazole pH 8.0, 116.9 mM maltose, 10% (v/v) glycerol, 5 mM b-mercaptoethanol). The elution was fractionated and analyzed using SDS-PAGE. Fractions containing IWS1 HEAT/TND were pooled and applied to 3.5k MWCO SnakeSkin dialysis tubing (Thermo Scientific) in lysis buffer at 4°C overnight. 1.5 mg of TEV protease with an N-terminal MBP tag was added to the sample prior to dialysis to remove the N-terminal His6-MBP tag. The dialyzed sample was applied to a HisTrap HP 5 mL, pre-equilibrated in lysis buffer. The flow-through containing IWS1 HEAT/TND was collected and subsequently applied to a self-packed XK column (Cytiva) with 15 mL of Amylose resin (New England Biolabs) pre-equilibrated in lysis buffer. The flow-through was concentrated using an Amicon 3,000 MWCO centrifugal filter unit (Millipore). The concentrated sample was applied to a Superose 6 Increase 10/300 GL (Cytiva), equilibrated in gel filtration buffer (20 mM Na·HEPES pH 7.4, 300 mM NaCl, 10% (v/v) glycerol, 1 mM TCEP). The elution was fractionated and analyzed by SDS-PAGE. Sample containing IWS1 HEAT/TND was concentrated using an Amicon 3,000 MWCO centrifugal filter unit (Millipore).

*H. sapiens* IWS1 ΔN (551-819) with an N-terminal 6xHis-MBP tag and TEV protease cleavage site was purified at 4°C. Cells expressing IWS1 ΔN were resuspended in lysis buffer and subsequently lysed by sonication. The lysate was centrifuged and cleared by ultra-centrifugation. The supernatant containing IWS1 ΔN was subsequently filtered using 5 mm and 0.45 mm syringe filters. The filtered supernatant was applied to a HisTrap HP 5 mL (Cytiva). The column was subsequently washed with 9 CV lysis buffer. A self-packed XK column (Cytiva) with 15 mL of Amylose resin (New England Biolabs) was attached to the HisTrap column. IWS1 ΔN was eluted from the HisTrap column using nickel elution buffer. The HisTrap column was removed, and the amylose column was washed with 10 CV low salt buffer (20 mM Na·HEPES pH 7.4, 150 mM NaCl, 30 mM imidazole pH 8.0, 10% (v/v) glycerol, 5 mM b-mercaptoethanol). A HiTrap Q HP 5 mL (Cytiva) column was attached to the amylose column, and IWS1 ΔN was eluted from the amylose column with 15 CV low salt amylose elution buffer (20 mM Na·HEPES pH 7.4, 150 mM NaCl, 30 mM imidazole pH 8.0, 116.9 mM maltose, 10% (v/v) glycerol, 5 mM b-mercaptoethanol). The amylose column was removed, and the HiTrap Q HP column was eluted with a 10 CV gradient of high salt buffer (20 mM Na·HEPES pH 7.4, 1000 mM NaCl, 30 mM imidazole pH 8.0, 10% (v/v) glycerol, 5 mM b-mercaptoethanol). The elution was fractionated and analyzed using SDS-PAGE. Fractions containing IWS1 ΔN were pooled and applied to 3.5k MWCO SnakeSkin dialysis tubing (Thermo Scientific) in low salt buffer at 4°C overnight. 1.5 mg of TEV protease with an N-terminal MBP tag was added to the sample prior to dialysis to remove the N-terminal His6-MBP tag. The dialyzed sample was applied to a 15 mL amylose column with a HiTrap SP HP 5 mL (Cytiva) column attached in tandem. The columns were washed with 8 CV low salt buffer. The amylose column was detached, then the SP column was eluted with a 12 CV gradient of high salt buffer. The elution was fractionated and analyzed using SDS-PAGE. Fractions containing IWS1 ΔN were pooled and concentrated using an Amicon 10,000 MWCO centrifugal filter unit (Millipore). The concentrated sample was applied to a Superdex 200 Increase 10/300 (Cytiva), equilibrated in gel filtration buffer (20 mM Na·HEPES pH 7.4, 300 mM NaCl, 10% (v/v) glycerol, 1 mM TCEP). The elution was fractionated and analyzed by SDS-PAGE. Sample containing IWS1 ΔN was concentrated using an Amicon 10,000 MWCO centrifugal filter unit (Millipore).

All concentrated samples were subsequently aliquoted, flash-frozen, and stored at -80°C prior to future use.

### RECQL5 expression and purification

Hi5 cells (600 mL) grown in ESF-921 media (Expression Systems) were infected with 300 µL of V1 virus for protein expression. The cells were grown for 48–72 h at 27 °C. Cells were harvested by centrifugation (238g, 4 °C, 30 min) and resuspended in lysis buffer. The cell resuspension was snapfrozen and stored at -80 °C.

Protein purification was performed at 4 °C. Frozen cell pellets were thawed and lysed by sonication. Lysates were cleared by centrifugation (18,000g, 4 °C, 30 min) and ultracentrifugation (235,000g, 4 °C, 60 min). The supernatant containing RECQL5 was filtered using 0.8-µm syringe filters (Millipore) and applied onto a HisTrap HP 5 ml (Cytiva), pre-equilibrated in lysis buffer. After sample application, the column was washed with 10 CV lysis buffer, 5 CV high salt buffer (1 M NaCl, 20 mM Na·HEPES pH 7.4, 10% (v/v) glycerol, 1 mM DTT, 30 mM imidazole pH 8.0, 0.284 µg mL™1 leupeptin, 1.37 µg mL™1 pepstatin A, 0.17 mg mL™1 PMSF, 0.33 mg mL™1 benzamidine), and 5 CV lysis buffer. The protein was eluted with a gradient of 0–100% elution buffer (300 mM NaCl, 20 mM Na·HEPES pH 7.4, 10% (v/v) glycerol, 1 mM DTT, 500 mM imidazole pH 8.0, 0.284 µg mL™1 leupeptin, 1.37 µg mL™1 pepstatin A, 0.17 mg mL™1 PMSF, 0.33 mg mL™1 benzamidine). Peak fractions were pooled and dialyzed for 16 h against 600 mL dialysis buffer (300 mM NaCl, 20 mM Na·HEPES pH 7.4, 10% (v/v) glycerol, 1 mM DTT, 30 mM imidazole) in the presence of 2 mg His6-TEV protease. The dialyzed sample was applied to a HisTrap HP 5 ml (Cytiva). The flowthrough containing RECQL5 was concentrated using an Amicon 50,000 MWCO centrifugal filter unit (Millipore) and applied to a Superdex 200 16/600 pg (Cytiva) size exclusion column, pre-equilibrated in gel filtration buffer (300 mM NaCl, 20 mM Na·HEPES pH 7.4, 10% (v/v) glycerol, 1 mM DTT). Peak fractions were concentrated to ∼100 µM, aliquoted, flash frozen, and stored at -80 °C.

### RECQL5 (1-620) expression and purification

*H. sapiens* RECQL5 (1-620) was expressed in BL21 (DE3) RIL cells at 37 °C until an OD of 0.6 at 600 nm was reached. Expression was induced with the addition of 1 mM IPTG and cells were grown at 18 °C overnight. Cells were centrifuged and pellets were resuspended in lysis buffer, then snap-frozen in liquid nitrogen and stored at -80 °C until purification.

Protein purification was performed at 4 °C. Frozen cell pellets were thawed and lysed by sonication. Lysates were cleared by centrifugation (47,800g, 4 °C, 30 min). The supernatant containing RECQL5 (1-620) was filtered using 0.8-µm syringe filters (Millipore) and applied onto a HisTrap HP 5 mL (Cytiva), pre-equilibrated in lysis buffer. After sample application, the column was washed with 6 CV lysis buffer, 5 CV high salt buffer (1 M NaCl, 20 mM Na·HEPES pH 7.4, 10% (v/v) glycerol, 1 mM DTT, 30 mM imidazole pH 8.0, 0.284 µg mL™1 leupeptin, 1.37 µg mL™1 pepstatin A, 0.17 mg mL™1 PMSF, 0.33 mg mL™1 benzamidine), and 5 CV lysis buffer. The protein was eluted with a 13 CV gradient of 0–100% elution buffer (300 mM NaCl, 20 mM Na·HEPES pH 7.4, 10% (v/v) glycerol, 1 mM DTT, 500 mM imidazole pH 8.0, 0.284 µg mL™1 leupeptin, 1.37 µg mL™1 pepstatin A, 0.17 mg mL™1 PMSF, 0.33 mg mL™1 benzamidine). Peak fractions were pooled and dialyzed for 16 h against 500 mL dialysis buffer (150 mM NaCl, 20 mM Na·HEPES pH 7.4, 10% (v/v) glycerol, 1 mM DTT, 30 mM imidazole) in the presence of 2 mg His6-TEV protease. The dialyzed sample was applied to a HisTrap HP 5 mL (Cytiva) with HiTrap Heparin HP 5 mL (Cytiva) connected in tandem. The columns were washed with 6 CV of dialysis buffer. The HisTrap HP 5 mL was removed, and the HiTrap Heparin HP 5 mL was eluted with a 13 CV gradient of 0–100% high salt buffer. The elution was fractionated and analyzed by SDS-PAGE, and fractions containing RECQL5 (1-620) were concentrated using an Amicon 30,000 MWCO centrifugal filter unit (Millipore) and applied to a Superdex 200 Increase 10/300 (Cytiva) column, pre-equilibrated in gel filtration buffer (300 mM NaCl, 20 mM Na·HEPES pH 7.4, 10% (v/v) glycerol, 1 mM DTT). The elution was fractionated and analyzed by SDS-PAGE. Sample containing RECQL5 (1-620) was concentrated using an Amicon 30,000 MWCO centrifugal filter unit (Millipore), aliquoted, flash frozen, and stored at -80 °C.

### Preparation of IWS1-GlySer-RPB5 helix for mass spectrometry analysis

500 ng of purified IWS1-GlySer-RPB5 helix was separated by SDS-PAGE using NuPAGE 4-12% Bis-Tris (Invitrogen) run in 1X MES buffer and stained with One-Step Blue Protein Gel Stain (Biotium). The gel was destained with water, then the protein band was excised from the gel and submitted to the Taplin Mass Spectrometry Facility at Harvard Medical School for in-gel digestion and micro-capillary LC/MS/MS analysis.

### H3K36cme3 octamer formation

Alkylation was used to install a trimethyl lysine analog on purified H3 containing the K36C mutation as previously described (62). Octamers with H3K36Cme3 were formed as previously described (36).

### Nucleosome preparation

The bp -4, bp +64, bp +115, and dinucleosome DNA vectors were used as templates for a large-scale PCR reaction as previously described (36). The PCR products were pooled, then purified using a HiTrap Q HP 5 mL (Cytiva) and eluted with a 20-40% NaCl gradient of TE buffer (10 mM Tris pH 8.0, 2 M NaCl, 1 mM EDTA pH 8.0). Peak fractions were pooled, ethanol-precipitated, and digested with TspRI (NEB) as previously described (36). An equal volume of phenol:chloroform:isoamyl alcohol 25:24:1 (Sigma-Aldrich) was added to TspRI-digested DNA, then the DNA was applied to a MaXtract High Density (Qiagen) 1.5 mL tube and centrifuged to separate the two phases. The aqueous phase containing DNA was further extracted using an equal volume of chloroform, then applied to another MaXtract High Density tube and centrifuged again. TspRI-digested DNA was purified from the aqueous phase using ethanol precipitation.

Nucleosomes containing H3K36Cme3-modified octamers and the TspRI-digested -4 DNA were reconstituted as described (61). Similarly, nucleosomes containing unmodified octamers and the TspRI-digested +115 DNA were reconstitued. Histone octamer and DNA were mixed at a 1:1 molar ratio in RB-High Buffer (20 mM Na·HEPES, pH 7.4, 2 M KCl, 1mM EDTA, pH 8.0, and 1 mM DTT), transferred to Slide-A-Lyzer Mini Dialysis Units 20K MWCO (Thermo Scientific), and gradient dialyzed against RB-Low Buffer (20 mM Na·HEPES, pH 7.4, 30 mM KCl, 1 mM EDTA, pH 8.0, and 1 mM DTT) for 18 hours at 4 ºC. The sample was further dialyzed against RB-Low Buffer for 3 hours at 4 ºC. The sample was centrifuged for 5 min at 21,000 x g to collect precipitate. The nucleosomes were subsequently purified by native PAGE using a Prep Cell apparatus (Bio-Rad). Peak fractions were pooled and concentrated in a 30K MWCO Amicon Ul-tra-4 Centrifugal Filter Unit (Millipore). Nucleosome concentration was quantified by absorbance at 280 nm. The molar extinction coefficient of the nucleosome was obtained by summing the molar extinction coefficients of the octamer and the DNA components at 280 nm. Dinucleosomes containing unmodified octamers and the TspRI-digested dinucleosome DNA were reconstitued similarly, except octamer and DNA were mixed at a 2:1 molar ratio.

### *In vitro* RNA extension assay

All concentrations refer to final reaction conditions. 120 nM of RNA (IDT) (5’-/6-FAM/-UUAUCACUGUC-3’) was incubated on ice with 120 nM of DNA (+64 linear DNA, +115 nucleosome, or dinucleosome) with a 3’ TspRI-digested overhang complementary to the RNA sequence. The RNA-DNA hybrid was incubated with 150 nM of Pol II on ice. A factor mix of 300 nM IWS1, 300 nM SPT6, 300 nM DSIF, 250 nM P-TEFb, 300 nM ELOF1, 300 nM PAF1c, and 300 nM RTF1 (if included) was added to the Pol II-RNA-DNA mixture. Buffer was added to adjust the final reaction conditions to 20 mM Na·HEPES pH 7.4, 65 mM NaCl, 10% glycerol, and 1 mM TCEP. 3 mM MgCl2 and 0.5 mM ATP was added to the sample, then the sample was incubated at 30 °C for 30 minutes. Transcription was started after the addition of 90 nM TFIIS, 0.01 mM CTP, 0.01 mM GTP, and 0.01 mM UTP. 3 µL reactions were quenched with 3 µL 2x STOP buffer (6.4 M urea, 50 mM EDTA, 2x TBE) at various timepoints (+64 linear DNA with RTF1: 0 sec, 10 sec, 30 sec, 45 sec, 60 sec, 90 sec. +115 nucleosome with RTF1: 0 min, 0.5 min, 1 min, 1.5 min, 2 min, 4 min. Dinucleosome with RTF1: 0 min, 1 min, 2 min, 3 min, 4 min, 6 min. +64 linear DNA without RTF1: 0 min, 1 min, 2 min, 5 min, 10 min, 20 min.). Samples were incubated with 30 mg Proteinase K (NEB) at 37 °C for 25 minutes, then treated with 36 µL formamide (molecular biology grade, Millipore) at 95 °C for 5 minutes. Samples for the +64 linear DNA or +115 nucleosome substrates were loaded onto 20% denaturing TBE-urea gels (8 M urea, 1x TBE, 10% formamide, 20% Bis-Tris acrylamide 19:1) and run with 1x TBE at 300 V for 90 minutes. Samples for the dinucleosome substrate were loaded onto 5% denaturing TBE-urea gels (8 M urea, 1x TBE, 10% formamide, 5% Bis-Tris acrylamide 19:1) and run with 1x TBE at 300 V for 20 minutes. The gels were imaged on an Amersham Typhoon (GE) using the 6-FAM fluorescence at 650 PMT.

Gel images were quantified using GelAnalyzer (GelAnalyzer 23.1.1 (available at www.gelanalyzer.com) by Istvan Lazar Jr., PhD and Istvan Lazar Sr., PhD, CSc). Original scanned TIFFs were opened in the GelAnalyzer application and bands were recognized as dark spots on a light background. Lanes were detected automatically with a threshold of 3, tilted, and equal width. Manual adjustment of lane boundaries was performed if lanes were curved and bands were not contained within the automatically defined lanes. Baseline was set with the rolling ball method and 10% peak width tolerance, which was applied to all lanes on the same gel image. Total extended RNA raw volume was measured for each lane by setting pixel boundaries in the region between the top of the primer band (bottom) and the edge of the well (top). Full-length product or strong band raw volume was measured for each lane by setting pixel boundaries in the region between the bottom of the full-length band (bottom) and the edge of the well (top). The full-length/strong band raw volume value was divided by the total extended RNA raw volume value for each lane, then imported into GraphPad Prism for mean, standard deviation, and statistical calculations as well as plotting.

Barrier indices for nucleosome entry sites were calculated as previously described (43), except using GelAnalyzer. Raw volume values were taken for the first nucleosome entry barrier band and second nucleosome entry barrier band setting pixel boundaries around each band. Raw volume values for the upper bands were taken by setting pixel boundaries in the region between the top of the barrier band (bottom) and the edge of the well (top). The raw volume values were used for the barrier index calculation, defined as barrier index = Intensity_barrier band_ / (Intensity_barrier band_ + Intensity_upper band_) (43). Calculated barrier indices were imported into GraphPad Prism for mean, standard deviation, and statistical calculations as well as plotting.

### RECQL5-IWS1 RNA extension assay

All concentrations refer to final reaction conditions. 120 nM of RNA (IDT) (5’-/6-FAM/-UUAUCACUGUC-3’) was incubated on ice with 120 nM of +64 linear DNA with a 3’ TspRI-digested overhang complementary to the RNA sequence. The RNA-DNA hybrid was incubated with 150 nM of Pol II on ice. A factor mix of 150 nM SPT6, 150 nM DSIF, 100 nM P-TEFb, 150 nM ELOF1, and 150 nM PAF1c was added to the Pol II-RNA-DNA mixture. Buffer was added to adjust the final reaction conditions to 20 mM Na·HEPES pH 7.4, 100 mM NaCl, 10% glycerol, and 1 mM TCEP. 3 mM MgCl_2_ and 1 mM ATP was added to the sample, then the sample was incubated at 30 °C for 30 minutes. The Pol II-factor mix was divided and IWS1 (0 nM, 150 nM, 300 nM, 600 nM, or 1500 nM) was added at 30 °C for 10 minutes. The mix was further divided and RECQL5 (1-620) (0 nM, 150 nM, 300 nM, 600 nM, 1500 nM) was added and incubated at 30 °C for 10 minutes. Transcription was started after the addition of 0.1 mM ATP, 0.1 mM CTP, 0.1 mM GTP, and 0.1 mM UTP. 3 µL reactions were quenched with 3 µL 2x STOP buffer (6.4 M urea, 50 mM EDTA, 2x TBE) after 150 seconds. Samples were incubated with 30 mg Proteinase K (NEB) at 37 °C for 25 minutes, then treated with 36 µL formamide (molecular biology grade, Millipore) at 95 °C for 5 minutes. Samples were loaded onto 20% denaturing TBE-urea gels (8 M urea, 1x TBE, 10% formamide, 20% Bis-Tris acrylamide 19:1) and run with 1x TBE at 300 V for 90 minutes. The gels were imaged on an Amersham Typhoon (GE) using the 6-FAM fluorescence at 650 PMT.

Gel images were quantified using GelAnalyzer (GelAnalyzer 23.1.1 (available at www.gelanalyzer.com) by Istvan Lazar Jr., PhD and Istvan Lazar Sr., PhD, CSc). Original scanned TIFFs were opened in the GelAnalyzer application and bands were recognized as dark spots on a light background. Lanes were detected automatically with a threshold of 3, tilted, and equal width. Manual adjustment of lane boundaries was performed if lanes were curved and bands were not contained within the automatically defined lanes. Baseline was set with the rolling ball method and 10% peak width tolerance, which was applied to all lanes on the same gel image.

Extended RNA raw volume was measured for each lane by setting pixel boundaries in the region between the bottom of the strong band above 106 nt and the edge of the well. Stalled RNA raw volume was also measured for each lane by setting pixel boundaries in the region between the top of the RNA primer band and the bottom of the strong band above 106 nt. The raw volume of the extended RNA and stalled RNA were added to obtain the “total above primer” raw volume. The extended RNA raw volume was divided by the total above primer raw volume value for each lane, then imported into GraphPad Prism for mean, standard deviation, and statistical calculations as well as plotting. To assess the statistical significance of each titration, a repeated measures one-way ANOVA was performed in GraphPad Prism, where the mean of each non-zero condition (150 nM, 300 nM, 600 nM, 1500 nM) was compared to the mean of the 0 nM condition. The 0 nM RECQL5, 0 nM IWS1 condition was excluded from the one-way ANOVA statistical analysis.

### Complex formation for size exclusion chromatography

All concentrations refer to final reaction conditions. 120 nM of RNA (IDT) (5’-/6-FAM/-UUAUCACUGUC-3’) was incubated on ice with 120 nM of the +64 linear DNA substrate with a 3’ TspRI-digested overhang complementary to the RNA sequence. The RNA-DNA hybrid was incubated with 150 nM of Pol II on ice. A factor mix of 600 nM IWS1, 300 nM SPT6, 300 nM DSIF, 250 nM P-TEFb, 300 nM ELOF1, and 300 nM PAF1c was added to the Pol II-RNA-DNA mixture. Buffer was added to adjust the final reaction conditions to 20 mM Na·HEPES pH 7.4, 65 mM NaCl, 10% glycerol, 3 mM MgCl2, and 1 mM TCEP. 1 mM ATP was added to the sample, then the sample was incubated at 30 °C for 30 minutes. The sample was centrifuged for 5 minutes at 13,000 x g and applied to a Superose 6 Increase 3.2/300 (Cytiva) column equilibrated in 20 mM Na·HEPES pH 7.4, 65 mM NaCl, 5% glycerol, 3 mM MgCl2, and 1 mM TCEP. Sample was detected at both 260 and 280 nm on an Äkta pure 25 with MicroKit. Corresponding fractions were analyzed for protein content using NuPAGE 4-12% Bis-Tris (Invitrogen) run in 1X MES buffer, stained with SyproRuby (Invitrogen), and detected using a SyproRuby fluorescence scan on an Amersham Typhoon (GE).

For Western blot analysis, samples were placed at 95 °C for 5 minutes and separated by NuPAGE 4-12% Bis-Tris (Invitrogen) gel ran in 1X MES buffer at constant voltage. 0.2 PVDF membrane (Millipore) was activated in 100% methanol and equilibrated with Transfer buffer (12 mM Tris pH 8.3, 96 mM glycine, 0.01% SDS, 10% methanol). Protein was transferred using the Mini Blot Module (Invitrogen catalog #B1000) according to manufacturer’s recommendations in Transfer buffer at 20 volts at room temperature for 65 minutes. The membrane was blocked in freshly made 5% (w/v) milk in PBS-T for 1 hour at room temperature. The membrane was subsequently transferred to IWS1 antibody (Proteintech 16943-1-AP) diluted 1:1000 or RNA polymerase II CTD 8WG16 antibody (Millipore 05-952-I) diluted to 0.2 µg/mL in 5% (w/v) milk in PBS-T and incubated overnight at 4 °C. After washing in PBS-T, the membrane was incubated with secondary antibody diluted in 5% (w/v) milk in PBS-T at room temperature for 2 hours. An Alexa Fluor Plus 647-labeled goat anti-rabbit IgG (Invitrogen A32733TR) was diluted 1:5000 and used for the membrane incubated with the IWS1 primary antibody. An Alexa Fluor 488 AffiniPure goat anti-mouse IgG (Jackson ImmunoResearch 115-545-003) was diluted to 2 µg/mL and used for the membrane incubated with the CTD primary antibody. The membrane was subsequently washed in PBS-T and analyzed using an Alexa-647 or Alexa-488 fluorescence scan on an Amersham Typhoon (GE) with automatic pre-scanned PMT. To maintain consistency between SyproRuby gels and Western blots, the same volume of each heated fraction (10 µL) was loaded into each gel well.

### Fluorescence anisotropy assay

A FAM-labeled 45mer double-stranded DNA was synthesized (IDT): 5’-/6-FAM-GCC GCG TAT AGG GTC CAT CAG AAT TCG GAT GAA CTC GGT

GTG AAG-3’. All concentrations refer to final reaction conditions. 20 microliter reactions were set up on ice containing 10 nM FAM-labeled 45mer double-stranded DNA, wild-type IWS1 or IWS1ΔC at 0, 10, 100, 1000, 2000, 3000, 4000, 5000, 6000, 7000, or 8000 nM, 20 mM Na·HEPES pH 7.4, 50 mM NaCl, 4% glycerol, 1 mM TCEP, 3 mM MgCl2, and 10 µg/mL BSA. 18 microliters of each reaction were transferred to a medium-binding 384-well microplate (Greiner 784076), then incubated on ice for 20 minutes. Plates were subsequently imaged on a Tecan Spark Multimode Microplate Reader using 470 nm excitation, 518 emission, gain high, and 100 flashes/read. A blank buffer well with no DNA or IWS1 was used for normalizing Z-height. A DNA only well was used as a blank.

### RNA polymerase II-DSIF-SPT6-PAF1c-TFIIS-IWS1-ELOF1-LEDGF-nucleosome complex cryo-EM sample preparation

All concentrations refer to final reaction conditions. RNA (5’-/6-FAM/-UUAUCACUGUC-3’) (1400 nM) and the H3K36Cme3 bp -4 nucleosome (1000 nM) were incubated for 5 minutes on ice. Pol II (700 nM) was further added for 5 minutes on ice. DSIF (1400 nM), SPT6 (1400 nM), PAF1c (1512 nM), RTF1 (2100 nM), P-TEFb (700 nM), IWS1 (3500 nM), ELOF1 (3500 nM), LEDGF (1750 nM), and MgCl2 were mixed and dialyzed into 400 mL of transcription buffer (50 mM NaCl, 20 mM Na·HEPES pH 7.4, 3 mM MgCl2, 4% (v/v) glycerol and 1 mM TCEP pH 8) at 4 °C for 2 hours. Next, 3’-dATP (1 mM) was added for 45 minutes at 30 °C to initiate phosphorylation by P-TEFb. Transcription was initiated by addition of CTP, GTP, UTP (1 mM each), and TFIIS (420 nM) at 30 °C for 30 minutes. The sample was centrifuged for 5 minutes at 21,300 x g and applied to a Superose 6 Increase 3.2/300 (Cytiva) column equilibrated in transcription buffer. Sample was detected at both 260 and 280 nm on an Äkta pure 25 with MicroKit. Corresponding fractions were analyzed for RNA extension products and protein content.

For RNA extension analysis, 5 µL of each fraction was mixed with 2x STOP buffer (6.4 M urea, 50 mM EDTA, 2X TBE). 40 µg proteinase K (NEB) was added to each sample and incubated at 37 °C for 30 minutes. 49.6 µL of formamide (molecular biology grade, Millipore) was added to each sample, then denatured at 95 °C for 15 minutes. 5 µL samples were separated by a 12% denaturing urea gel (8 M urea, 1X TBE, 12% acrylamide:bis-acrylamide, 10% formamide) and RNA product was detected using a 6-FAM fluorescence scan on a Amersham Typhoon (GE) with 500 PMT. Proteins were detected using a NuPAGE 4-12% Bis-Tris (Invitrogen) ran in 1X MES buffer and stained with One-Step Blue Protein Gel Stain (Biotium).

Fractions containing complex were crosslinked with 0.1 % (v/v) glutaraldehyde for 10 minutes on ice followed by quenching with 2.4 mM aspartate and 2 mM lysine for 10 minutes on ice. Samples were dialyzed for three hours in cryo-EM buffer (20 mM Na·HEPES pH 7.4, 1 mM TCEP pH 8, 50 mM NaCl, and 3 mM MgCl2). Samples were centrifuged for 5 minutes at 21,300 x g following dialysis.

Complex concentration was determined by measuring absorbance at 280 nm. The molar extinction coefficient of the complex was obtained by summing the molar extinction coefficients of all protein and nucleic acid components at 280 nm.

UltrAuFoil R 2/2 on 200 mesh gold grids were glow-discharged for 120 s at 30 mA using a Pelco Easiglow plasma discharge system. 4 µL of dialyzed sample was applied to grids for 8 s, blotted for 3 s with a blot force of 8, and vitrified by plunging into liquid ethane using a Vitrobot Mark IV (FEI) at 4 °C and 100 % humidity.

### RNA polymerase II-DSIF-SPT6-PAF1c-TFIIS-IWS1-ELOF1-LEDGF-nucleosome complex cryo-EM data collection and image processing

Cryo-EM data were collected on a ThermoFisher Scientific Titan Krios operated at 300 keV equipped with Falcon4i direct electron detector and a Selectris energy filter. Data collection was automated using EPU software. A single dataset was collected at a pixel size of 1.19 Å with a defocus range of -1.8 to -0.6 µm. The dataset yielded 30,316 micrographs. The dataset was collected with 48 movie frames at an exposure time of 4.21 s with an electron flux of 14.04 e-Å-2 s-1 for a total exposure of 37 e-Å-2.

Initial image processing was conducted in cryoSPARC (63).Movies were aligned using cryoSPARC Live Patch Motion Correction followed by CTF estimation. The cryoSPARC Blob picker was used to pick 4,597,368 particles. Particles were extracted at a box size of 460 pixels and downsampled to 400 pixels. Initial junk particles were removed, and then the remaining 1,691,625 particles that revealed clear density for Pol II and elongation factors were subjected to 3D classification using a mask encompassing the IWS1 HEAT/TND, DSIF NGN, and ELOF1. From this 3D classification, a class of 777,108 particles containing clear density for the IWS1 HEAT/TND was selected for downstream processing steps.

The 777,108 particles were re-extracted with a box size of 460 pixels and subjected to homogeneous refinement. Local CTF correction, global CTF correction, removal of duplicate particles, and reference motion correction were then followed by another round of homogeneous refinement. The corresponding 2.4 Å map (map A) revealed high resolution features for Pol II, the IWS1 HEAT/TND, DSIF NGN, and ELOF1, but low-resolution features for all other components. To visualize the other components, the 762,523 particles from map A were subjected to various rounds of classification and refinement. All mask generation, local refinements, and 3D classifications were performed in cryoSPARC unless stated otherwise.

Iterative 3D classifications using a mask encompassing the IWS1 HEAT/TND, ELOF1, and DSIF NGN domains (“IWS1+ELOF1 mask”) was performed to enrich for the IWS1 HEAT/TND, ELOF1, and DSIF NGN. After performing 3D classification, 378,540 particles were subjected to non-uniform refinement followed by local refinement using the focus mask to produce a IWS1+ELOF1 map of 2.6 Å (map B). A similar processing workflow was used for the IWS1 C-term map (“IWS1 C-term mask,” 2.4 Å, 156,410 particles, map C), TFIIS map (“TFIIS mask,” 4.3 Å, 36,198 particles, map D), DSIF map (“DSIF mask,” 2.7 Å, 79,487 particles, map E), RTF1 map (“RTF1 mask,” 2.8 Å, 27,130 particles, map F), 10-subunit Pol II map (mask encompassing Pol II excluding RPB4/7 “10-subunit Pol II mask,” 2.5 Å, 115,139 particles, map G), non-template DNA map (“Non-template DNA mask,” 2.6 Å, 113,665 particles, map H), RPB5+DNA map (“RPB5+DNA mask,” 2.7 Å, 59,649 particles, map I), CDC73 Ras-like domain map (“CDC73 mask,” 3.4 Å, 43,339 particles, map J), PAF1+LEO1 map (“PAF1+LEO1 mask,” 3.2 Å, 28,545 particles, map M), and SPT6 map (“SPT6 mask,” 2.7 Å, 90,114 particles, map O).

A mask encompassing the CTR9 TPRs and WDR61 (“CTR9+WDR61 mask”) was used to perform iterative 3D classifications of the 762,523 particles from map A. To enrich for the CTR9 N-terminus, another mask encompassing CTR9 residues 1-447 (“CTR9 TPR mask”) was used to perform further 3D classifications. The resulting 83,886 particles were subjected to non-uniform refinement followed by local refinement using the CTR9+WDR61 mask to produce a CTR9+WDR61 map (3.8 Å, 63,837 particles, map K), as well as local refinement using the CTR9 TPR mask to produce a CTR9 TPR map (2.6 Å, 63,837 particles, map L).

The 28,545 particles in the PAF1+LEO1 map were subjected to additional 3D classification using a mask encompassing the LEO1 C-terminus (“LEO1 C-term mask”) to enrich for the LEO1 C-terminal helix. The resulting 6,771 particles were subjected to non-uniform refinement followed by local refinement using the focus mask to produce a LEO1 C-term map (4 Å, 6,771 particles, map N).

The 90,114 particles in the SPT6 map were subjected to local refinement using the IWS1+ELOF1 mask to improve resolution of the SPT6-IWS1 interaction. This produced a SPT6 N-term map (2.9 Å, 90,114 particles, map P).

While map A revealed clear density for all elongation factors, we were unable to see the downstream nucleosome. Instead, all we could visualize in our maps was significant amounts of noise at low contour levels where we expected the downstream nucleosome, likely due to flexibility between the Pol II and the downstream nucleosome interface. Consequently, we created a mask encompassing the visible low-resolution features (“Nucleosome mask”) and performed particle subtraction in RELION (v4.0) (64) using map A particles. Subtracted particles were imported back into cryoSPARC for iterative rounds of 2D classification and heterogeneous refinement to obtain a distinct nucleosome class of 85,098 particles. 3D classification using a mask encompassing the entire density was performed to obtain a 33,160 particle class containing density for the LEDGF PWWP. To further enrich the PWWP density, 3D classification using a mask encompassing the PWWP binding site for the distal H3 (“PWWP mask”) was performed. The resulting class of 19,562 particles was subjected to non-uniform refinement followed by local refinement of the entire density to produce a nucleosome map of 3.5 Å (map Q).

We were unable to simultaneously visualize both Pol II and the nucleosome in the original data set (particles and processing corresponding to map A). Therefore, we reverted the 19,562 particle class from the nucleosome map Q to their original particles in RELION (v5.0) (64).We performed particle subtraction of the reverted particles in RELION (v5.0) using a map encompassing the downstream nucleosome, DNA, and Pol II without RPB4/7 (“10-subunit Pol II with nucleosome mask”). Subtracted particles were imported back into cryoSPARC for ab-initio reconstruction, non-uniform refinement, and local refinements with focus masks of the nucleosome or 10-subunit pol II to produce a nucleosome placement map (7.6 Å, 19,562 particles, map R) and a local 10-subunit Pol II map (8.8 Å, 19,562 particles, map S). Map S was aligned to map A using Align 3D Maps in cryoSPARC, then map R was aligned to the aligned map S.

### RNA polymerase II-DSIF-SPT6-PAF1c-TFIIS-IWS1-ELOF1-LEDGF-nucleosome complex model building and refinement

All fitting was performed in UCSF ChimeraX v1.7-1.9 (65) and refinements were performed in ISOLDE (66). An initial activated elongation complex model was generated where Pol II, DSIF, SPT6, PAF1c, and RTF1 from PDB 6TED were modeled into map A. The HEAT/TND of IWS1 and TFIIS from PDB 9EGZ were also modeled into map A. PAF1 and LEO1 from PDB 6TED were replaced with an AlphaFold-Multimer generated model of PAF1 and LEO1 with RPB2. Similarly, SPT6 was replaced with an AlphaFold-Multimer model of SPT6-RPB4-RPB7. AlphaFold-Multimer generated models of ELOF1-RPB2, IWS1-RPB5, IWS1-SPT4-SPT5, IWS1-SPT6, IWS1-RPB2, IWS1-ELOF1, and IWS1-RPB1 were also used to build initial models of ELOF1 bound to the elongation complex, the IWS1 RPB5 helix (residues 528-542), the IWS1-NGN interaction (IWS1 residues 693-721), the SPT6 N-terminal helix (residues 207-230) interaction with the IWS1 HEAT/TND, the IWS1 RPB2 lobe loop (IWS1 residues 749-760), the IWS1-ELOF1 belt (IWS1 residues 760-770), and the IWS1-RPB1 jaw and cleft interactions (IWS1 residues 787-819), respectively.

Maps A and G were used to refine all subunits of Pol II. Missing residues in the RPB2 protrusion domain from PDB 6TED were also built. Map B was used to refine IWS1 residues 554-712 and 749-770. Gaussian-filtered map B to 1.5 standard deviations provided sufficient density to place IWS1 residues 713-721 based on the AlphaFold model. Map C was used to refine IWS1 residues 805-819. IWS1 residues 787-795 from the IWS1-RPB1 AlphaFold prediction were docked into the Map C density. Map D was used to refine TFIIS. The Zn finger domain of TFIIS was rigid-body built into Gaussian-filtered map D. Map E was used to refine SPT4 and SPT5. Map F was used to refine the pincer helices and fastener of RTF1 and rigid-body fit the RTF1 linker and Plus3 domain. Map H was used to unambiguously assign the register as bp -9 by defining purine and pyrimidine bases in the DNA · RNA hybrid. Map H was also used to build the backtracked RNA strand using PyMOL v2.5.2 (67), and model the backbone of the singlestranded nontemplate DNA in the transcription bubble. Map I was used to rigid-body fit the IWS1 RPB5 helix (residues 528-542). Map J was used to rigid-body fit an AlphaFold model of the CDC73 Ras-like domain (residues 360-531). Map J was also Gaussian-filtered to 2 standard deviations to rigid-body fit the tSH2 domain. Map K was used to refine WDR61, CTR9 residues 440-810, and CDC73 residues 218-260. Map K was also used to rigid-body fit CTR9 residues 3-439 and PAF1 residues 25-113. Map L was used to rigid-body fit the CTR9 trestle helix. Map M was used to rigid-body fit PAF1 and LEO1. Map N was used to rigid-body fit the LEO1 C-terminal helix. Map O was used to refine RPB4 and RPB7 and fit SPT6. Map P was used to refine SPT6 residues 207-218, then map P Gaussian-filtered to 2 standard deviations was used to rigid-body fit SPT6 residues 219-230 using the IWS1-SPT6 AlphaFold model.

The nucleosome model was built separately from the activated elongation complex model. PDB 6S01 was fit into map Q, then the downstream DNA was extended in PyMOL v2.5.2 (67). Regions of the DNA from PDB 6S01 which did not have sufficient density in map Q were deleted.

To position the nucleosome relative to the activated elongation complex, the nucleosome model was fit into the aligned map R in UCSF ChimeraX v1.9 (65).

Due to low resolution, side-chains were removed for the following regions: TFIIS (residues 250-301), CDC73 (360-531), SPT6 (219-230, 332-339, 426-430, 483-488, 696-704, 759-763, 775-808, 815-882, 1075-1135, 1172-1175, 1227-1322,1323-1514), PAF1 (45-113, 151-371), LEO1 (356-512, 529-555), CTR9 (3-439, 811-892), IWS1 (528-542, 712-721, 787-795), SPT5 (270-397, 524-645, 753-781), RTF1 (353-502, 597-624), and LEDGF (1-91). Nucleotide bases were removed for the single-stranded non-template DNA.

The model was subsequently real space refined in PHENIX (68) with secondary restraints.

### Composite map of RNA polymerase II-DSIF-SPT6-PAF1c-TFIIS-IWS1-ELOF1 complex

To generate a composite map, maps B-P were aligned to map A using Align 3D Maps in cryoSPARC. Focus masks for each aligned map were prepared by generating a volume of the focus area using the atomic model of the activated elongation complex and molmap command with 15 Å resolution in UCSF ChimeraX v1.9 (65) followed by conversion to a mask in cryoSPARC. The chains used for each molmap are detailed as follows: map B, IWS1 residues 554-770, ELOF1, SPT4, SPT5 residues 160-269; map C, IWS1 residues 787-819; map D, TFIIS, RPB9; map E, SPT5 residues 270-753; map F, RTF1, SPT5 residues 754-781, PAF1 residues 120-149; map G, RPB3, RPB8, RPB10, RPB11, RPB12; map H, nontemplate DNA positions -27-10, template DNA positions -10-29, RNA; map I, IWS1 residues 528-542, RPB5, nontemplate DNA positions 3-18, template DNA positions -18--3; map J, CDC73 residues 360-531, RPB1, RPB6; map K, CTR9, WDR61, PAF1 residues 25-113, CDC73 residues 218-260; map L, CTR9 residues 830-892; map M, RPB2, PAF1 residues 151-371, LEO1 residues 356-514; map N, LEO1 residues 515-555; map O, RPB4, RPB7, SPT6 residues 266-1322; map P, SPT6 residues 207-230. The Frankenmap package from Warp (69) was then used to produce the composite map T using the aligned maps B-P and their corresponding focus masks.

### RNA polymerase II-RECQL5 complex formation and cryo-EM analysis and image processing

ECs were formed on a bubble scaffold with the following nucleic acid sequence: template DNA (IDT) (5′-GCT CCC AGC TCC CTG CTG GCT CCG AGT GGG TTC TGC CGC TCT CAA TGG-3′), non-template DNA (IDT) (5′-CCA TTG AGA GCG GCC CTT GTG TTC AGG AGC CAG CAG GGA GCT GGG AGC-3′) and RNA (IDT) (5’-/6-FAM/ UUA AGG AAU UAA GUC GUG CGU CUA AUA ACC GGA GAG GGA ACC CAC U-3′) (35). The scaffold contains 9 bps of RNA–DNA hybrid, 37 nt of exiting RNA, 11 nt of bubble, 14 bps of upstream DNA and 23 bps of downstream DNA. RNA and template DNA were mixed in equimolar ratios and annealed by incubating the nucleic acids at 95 °C for 5 min and then decreasing the temperature in steps of 1 °C min–1 to a final temperature of 30 °C in a thermocycler. All concentrations refer to the final concentrations used in the assay.

S. scrofa Pol II (300 nM) and the RNA–template hybrid (500 nM) were incubated for 5 min on ice. The non-template DNA (667 nM) was added and the sample was incubated for another 5 min. RECQL5 (500 nM) and ADP-BeF3 (1 mM) were added in a final buffer containing 150 mM NaCl, 20 mM Na·HEPES pH 7.4, 3 mM MgCl2, 4% (v/v) glycerol and 1 mM TCEP pH 8. The sample was incubated for 30 min on ice.

The sample was mildly cross-linked using glutaraldehyde. The sample was dialyzed into dialysis buffer (150 mM NaCl, 20 mM Na·HEPES pH 7.4 at 25 °C, 3 mM MgCl2, 1 mM TCEP pH 8) for 3 h.

Quantifoil R2/1 on 200 Mesh copper grids were glow discharged for 30 s at 15 mA using a Pelco Easiglow plasma discharge system. 4 µL of dialyzed sample was applied to grids for 8 s, blotted for 5 s with a blot force of 8, and vitrified by plunging into liquid ethane using a Vitrobot Mark IV (FEI) at 5 °C and 100 % humidity.

Cryo-EM data was collected on a ThermoFisher Scientific Talos Arctica operating at 200 kV equipped with a Gatan K3 direct electron detector. Data collection was automated using SerialEM software. Data was collected at a pixel size of 1.1 Å with a defocus range of 1 to 2.2 µm. The data set yielded 227 micrographs with 50 movie frames at an exposure time of 3.99 s with an electron flux of 13.045 e− per A2 per s for a total exposure dose of 52.11 e– per Å2.

Initial data processing was performed in cryoSPARC (63). Movie alignment was performed using patch motion correction followed by patch CTF estimation. Blob picker was then used and 52,539 particles were extracted at a box size of 350 pixels. Two rounds of heterogenous refinement were performed to enrich for Pol II. Remaining particles were subjected to non-uniform refinement. To enrich for the RECQL5 KIX domain, we performed masked classification without image alignments in RELION (v4.0) (64). After classification, particles were re-imported to cryoSPARC and subjected to non-uniform refinement (map U).

### Fit of predicted ELOA helix in coulomb potential map

PDB 8OF0 and EMD-16831 were opened in UCSF ChimeraX v1.8 (65) and EMD-16831 was Gaussian-filtered to 1 standard deviation. RPB1 from a RPB1-ELOA1 AlphaFold prediction was aligned to the RPB1 model in PDB 8OF0, placing the predicted RPB1 jaw-interacting ELOA1 helix (residues 756-772) from the AlphaFold model into the unassigned density in EMD-16831.

### Sequence alignments

Sequence alignments were performed using PROMALS3D (70) with structure inputs. IWS1 homolog sequences were fetched from UniProt (71). IWS1 homolog structures were fetched from the AlphaFold Protein Structure Database (72,73) using their UniProt ID or predicted using AlphaFold2 (74) when not available in the database.

### Selection of Candidates for AlphaFold-Multimer (AF-M) Screen

Human proteins with the GO term “transcription elongation from RNAP2 promoters” (GO:0006368) were selected to screen against RPB1 (2). To identify other interactors of RPB1 beyond those involved in elongation, “top” interactors with more than 5 evidence types listed for the *H. sapiens* RPB1 identifier on BioGRID (75) were selected. A total of 80 proteins were selected for the screen. All proteins were screened against the *H. sapiens* RPB1 (Uniprot ID P24928) residues 1-1487.

### AF-M Structure Generation

Unless stated otherwise, all interactions were predicted using AlphaFold-Multimer (76) (AF-M) via an in-house modified version of ColabFold 1.52 (46,77) . AF-M was run with v2.3 weights, 1 ensemble, 3 recycles, templates enabled, dropout disabled, and maximum Multiple Sequence Alignments (MSA) depth settings (max_seq = 508, max_extra_seq = 2048). MSAs (paired and unpaired) were fetched from a remote server via the MMSeqs2 (78) API that is integrated into the ColabFold (77) pipeline. To reduce computational burden during the interaction screens we only ran 3 models (1,2, and 4) and did not run any pairs with total amino acid lengths exceeding 3,600 residues (GPU memory limit). The generation time of each structure prediction as well as the run settings have been recorded as REMARK lines in the PDB output files.

### AF-M Prediction Analysis

Python scripts were utilized to analyze structural predictions generated by AF-M as previously described (46). Briefly, confident interchain residue contacts were extracted from AF-M structures by identifying proximal residues (heavy atom distance <5 Å) where both residues have pLDDT values >50 and PAE score <15 Å. All downstream analysis of interface statistics (average pLDDT, average PAE) were calculated on data from these selected inter-residue pairs (contacts). Average interface pLDDT values above 70 are generally considered confident. The average models score was calculated by averaging the number of independent AF-M models that predicted a specific inter-residue contact across all unique pairs in all models. This number was additionally normalized by dividing by the number of models run to produce a final average model score that ranges from 0 (worst) to 1 (best). An average models value above 0.5 is generally considered confident. pDockQ (74) estimates of interface accuracy scores were calculated independently of the contact analysis described above using code that was adapted from the original implementation (79). pDockQ values above 0.23 are considered confident.

### SPOC Interaction Analysis

The random forest classifier algorithm (Structure Prediction and Omics Classifier) SPOC (46) was used to score the binary interaction predicted by via AF-M. Briefly, this classifier was trained to distinguish biologically relevant interacting protein pairs from non-biologically likely interaction pairs in AF-M interaction screens. SPOC assigns each interaction a score that ranges from 0 (worst) to 1 (best). Higher scores indicate that AF-M interface data and several types of externally sourced omics datasets support the potential existence of the binary interaction produced by AF-M. The highest confident interactions are generally associated with SPOC scores above 0.5.

## Figure generation

All figures were generated using Adobe Illustrator, GraphPad Prism, and UCSF ChimeraX v1.7-1.9 (65). Sequence alignments were formatted in Unipro UGENE (80), then exported to .svg format and imported into Adobe Illustrator.

Local resolution estimations for each map were generated using the Local Resolution Estimation job in cryoSPARC. Local resolution maps were generated in ChimeraX using Surface Color of the refinement volume by the volume data of the local resolution output. FSC resolution values were obtained using the following calculation: Resolution (Å) = (box size * pixel size) / wave number. Wave number values were generated in cryoSPARC after FSC-mask auto-tightening. FSC curves were plotted in GraphPad Prism using the FSC tightmask values and calculated resolutions.

