## Supplemental Figures for "Structure and function of IWS1 in transcription elongation"

### Figure S1

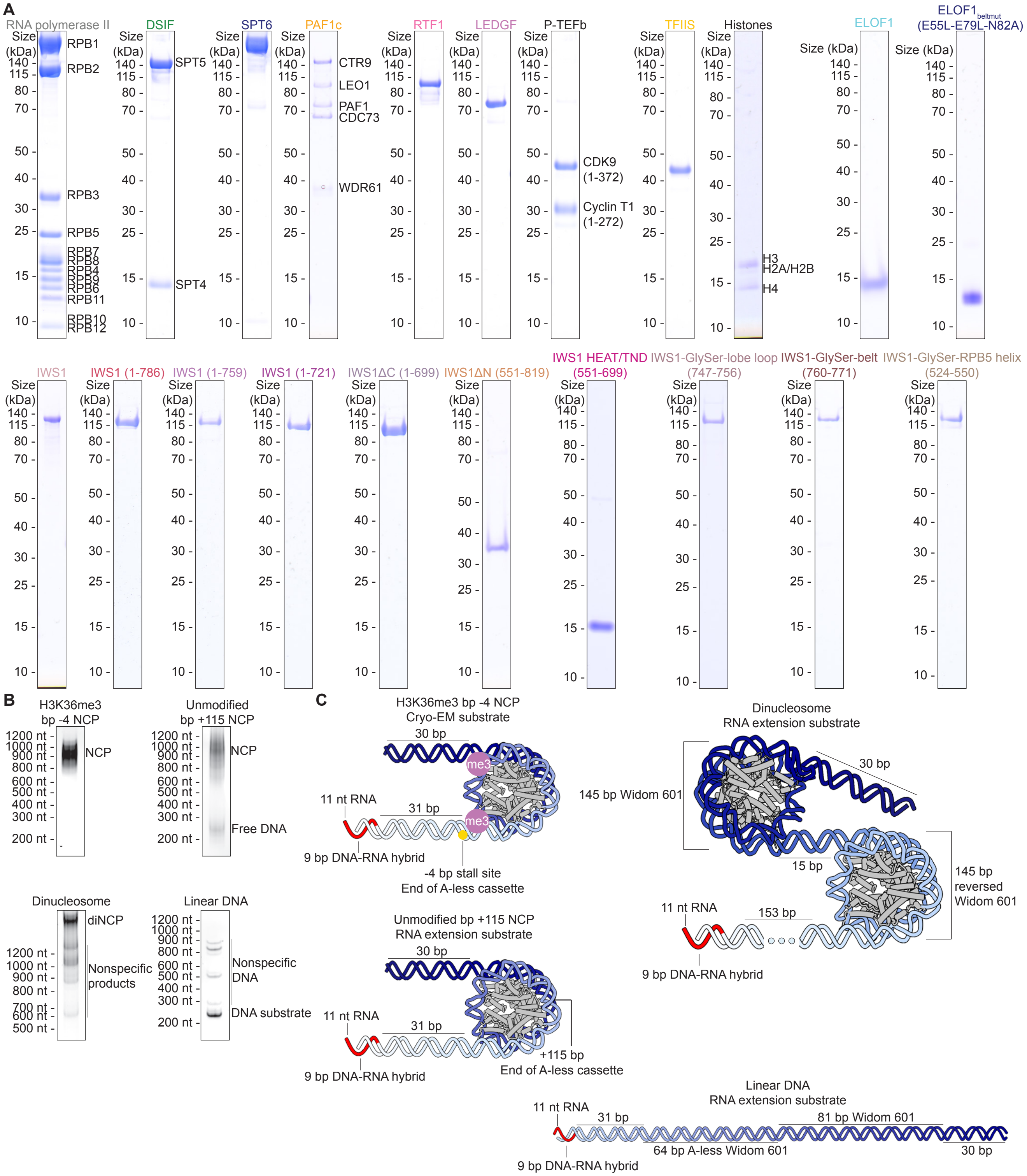
A
RNA polymerase II
RPB1
RPB2
RPB3
RPB5
RPB7
RPB8
RPB4
RPB9
RPB6
RPB11
RPB10
RPB12
DSIF
SPT5
SPT4
SPT6
PAF1c
CTR9
LEO1
PAF1
CDC73
WDR61
RTF1
LEDGF
P-TEFb
CDK9 (1-372)
Cyclin T1 (1-272)
TFIIS
Histones
H3
H2A/H2B
H4
ELOF1
ELOF1beltmut (E55L-E79L-N82A)
IWS1
IWS1 (1-786)
IWS1 (1-759)
IWS1 (1-721)
IWS1ΔC (1-699)
IWS1ΔN (551-819)
IWS1 HEAT/TND (551-699)
IWS1-GlySer-lobe loop (747-756)
IWS1-GlySer-belt (760-771)
IWS1-GlySer-RPB5 helix (524-550)
Size (kDa)
140
115
80
70
50
40
30
25
15
10
B
H3K36me3 bp -4 NCP
Unmodified bp +115 NCP
NCP
Free DNA
Dinucleosome
diNCP
Nonspecific products
Linear DNA
Nonspecific DNA
DNA substrate
1200 nt
1000 nt
900 nt
800 nt
700 nt
600 nt
500 nt
400 nt
300 nt
200 nt
C
H3K36me3 bp -4 NCP Cryo-EM substrate
30 bp
me3
11 nt RNA
31 bp
9 bp DNA-RNA hybrid
-4 bp stall site End of A-less cassette
Unmodified bp +115 NCP RNA extension substrate
+115 bp End of A-less cassette
Dinucleosome RNA extension substrate
145 bp Widom 601
15 bp
145 bp reversed Widom 601
153 bp
Linear DNA RNA extension substrate
81 bp Widom 601
64 bp A-less Widom 601

**Figure S1. Constructs and substrates. (A)** SDS-PAGE of purified protein components. Samples were loaded onto 4-12% Bis-Tris SDS-PAGE gels, run in 1x MES buffer, and stained with OneStep Blue. **(B)** Native gels of substrates used. Samples were run on TBE native gels and stained with Sybr Gold. **(C)** Schematic of substrates used. DNA is colored from light to dark blue in the direction of transcription. RNA colored in red. Histones colored in gray.

Figure S2

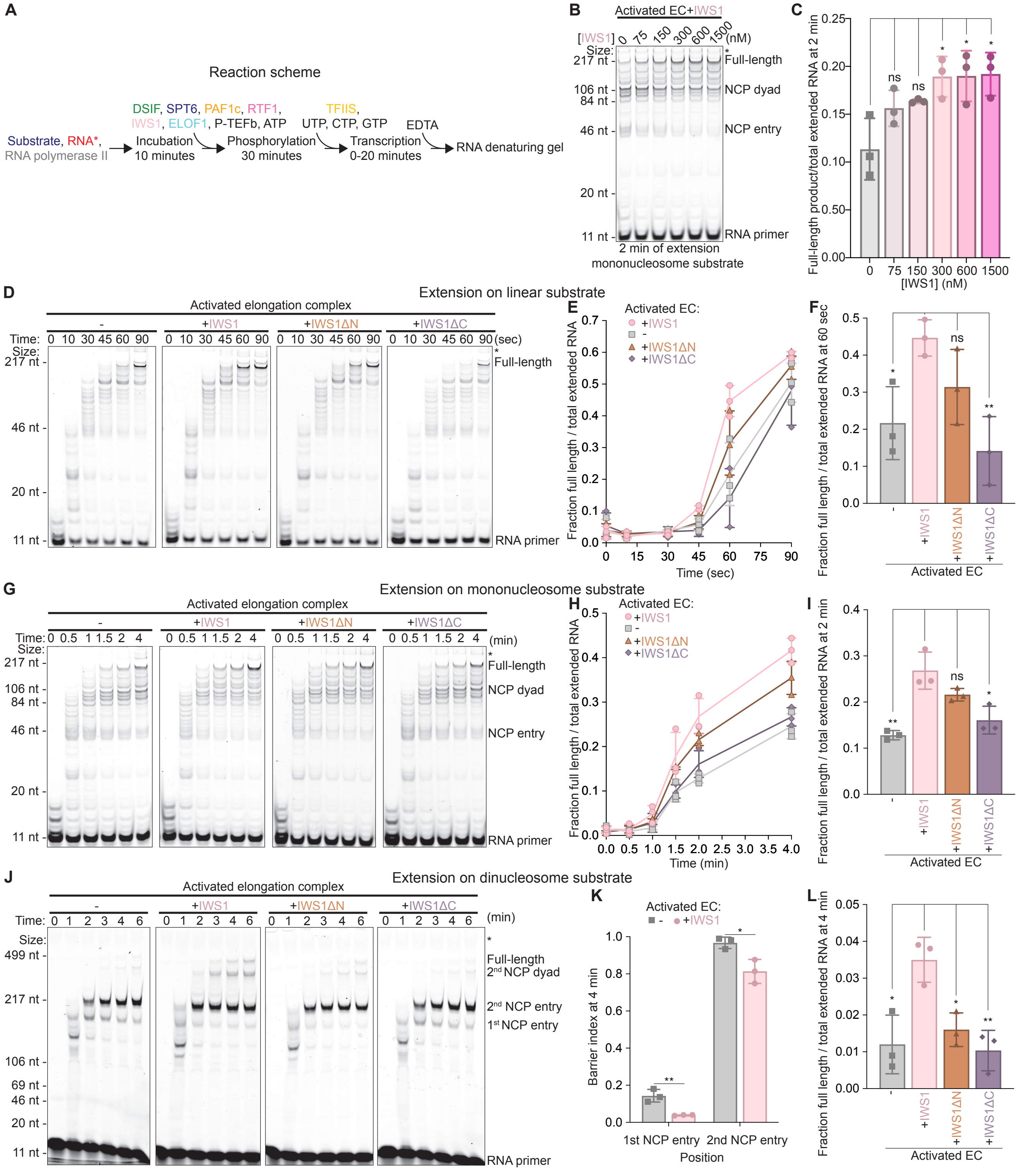
A
Reaction scheme
Substrate, RNA*,
RNA polymerase II
DSIF, SPT6, PAF1c, RTF1,
IWS1, ELOF1, P-TEFb, ATP
Incubation
10 minutes
Phosphorylation
30 minutes
TFIIS,
UTP, CTP, GTP
Transcription
0-20 minutes
EDTA
RNA denaturing gel
B
Activated EC+IWS1
[IWS1] 0 75 150 300 600 1500 (nM)
2 min of extension
mononucleosome substrate
Full-length
NCP dyad
NCP entry
RNA primer
C
Full-length product/total extended RNA at 2 min
[IWS1] (nM)
D
Extension on linear substrate
Activated elongation complex
+IWS1
+IWS1ΔN
+IWS1ΔC
Time: 0 10 30 45 60 90 (sec)
E
Activated EC:
Fraction full length / total extended RNA
Time (sec)
F
Fraction full length / total extended RNA at 60 sec
Activated EC
G
Extension on mononucleosome substrate
Time: 0 0.5 1 1.5 2 4 (min)
H
Time (min)
I
Fraction full length / total extended RNA at 2 min
J
Extension on dinucleosome substrate
Time: 0 1 2 3 4 6 (min)
2nd NCP dyad
2nd NCP entry
1st NCP entry
K
Barrier index at 4 min
1st NCP entry
2nd NCP entry
Position
L
Fraction full length / total extended RNA at 4 min

**Figure S2. RNA extension assays on different substrates. (A)** Reaction scheme of RNA extension assay. **(B)** RNA extension assay with IWS1 titration on a mononucleosomal substrate. The samples on the gel were quenched 2 min after addition of CTP, GTP and UTP (10 μM). Samples were separated by denaturing gel electrophoresis. RNA extension was monitored using FAM dye on RNA. Initial RNA and extended products are marked. Results are representative of at least three independent experiments. **(C)** Quantification of (B) from n = 3 independent experiments. * p<0.05, ns = not significant relative to -IWS1 using unpaired t-test. **(D)** RNA extension assays on linear DNA with no IWS1, full-length IWS1, IWS1ΔN, and IWS1ΔC. Results are representative of at least three independent experiments. **(E)** Quantification of (D) from n = 3 independent experiments. Individual data points and standard deviation are shown. **(F)** Quantification of 60 sec timepoint from (D) from n = 3 independent experiments. Individual data points and standard deviation are shown. **(G)** RNA extension assays on mononucleosome with no IWS1, full-length IWS1, IWS1ΔN, and IWS1ΔC. Results are representative of at least three independent experiments. **(H)** Quantification of (G) from n = 3 independent experiments. Individual data points and standard deviation are shown. **(I)** Quantification of 2 min timepoint from (G) from n = 3 independent experiments. Individual data points and standard deviation are shown. **(J)** RNA extension assays on dinucleosome substrate with no IWS1, full-length IWS1, IWS1ΔN, and IWS1ΔC. Results are representative of at least three independent experiments. **(K)** Quantification of nucleosome barrier index at two NCP entry sites from n = 3 independent experiments. **(L)** Quantification of 4 min timepoint from (J) from n = 3 independent experiments. Individual data points and standard deviation are shown. Activated EC = Pol II, DSIF, SPT6, PAF1c including RTF1, TFIIS, ELOF1. Columns represent mean values across n = 3 independent experiments and error bars represent standard deviations for C, E, F, H, I, K, and L. Not fully denatured RNA marked with asterisk (*) on gels. * p<0.05, ** p<0.01, *** p<0.001, ns=not significant relative to +IWS1 using unpaired t-test for F, I, K, and L.

Figure S3

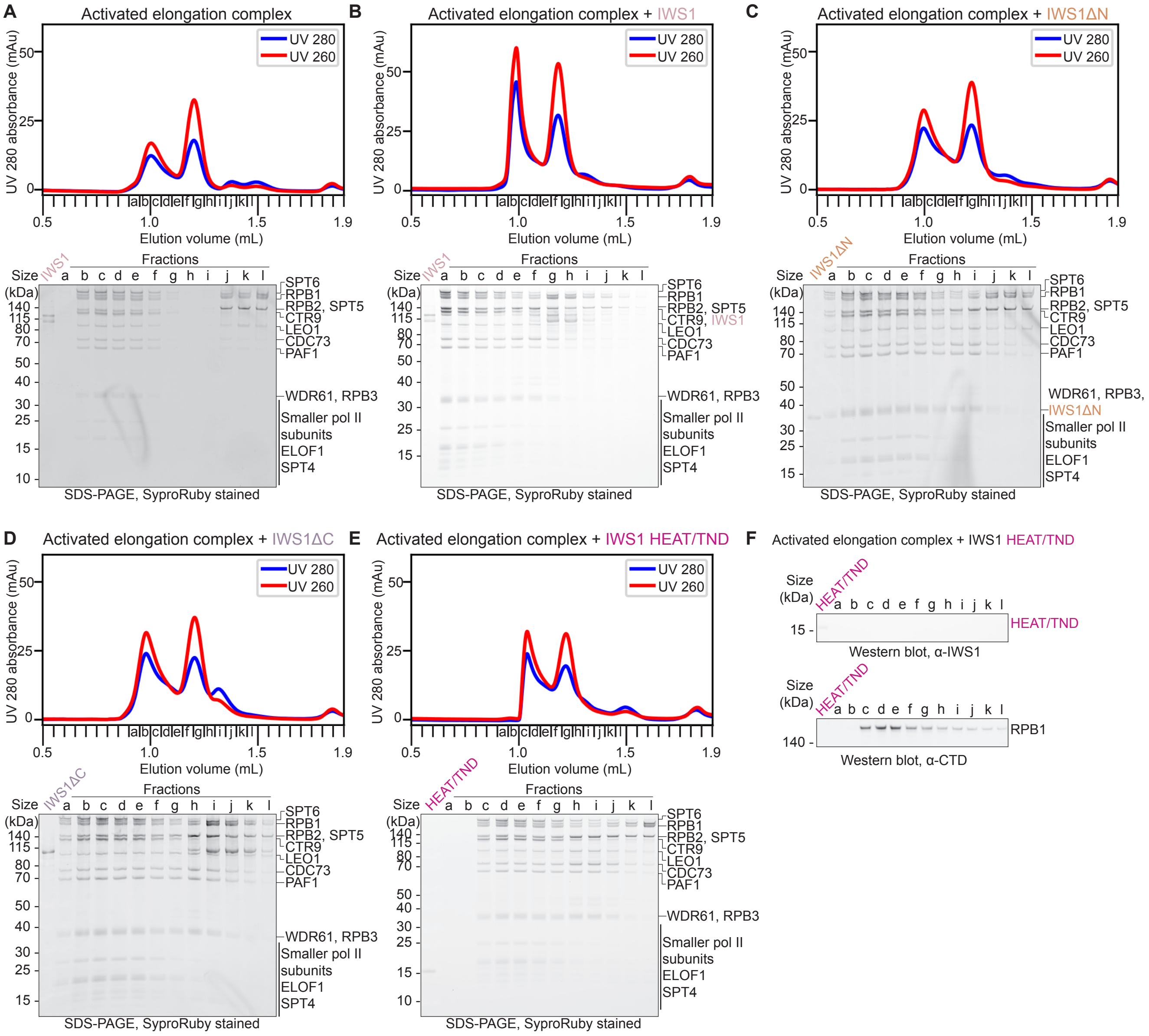
A
Activated elongation complex
UV 280
UV 260
UV 280 absorbance (mAu)
Elution volume (mL)
Fractions
Size (kDa)
IWS1
SPT6
RPB1
RPB2, SPT5
CTR9
LEO1
CDC73
PAF1
WDR61, RPB3
Smaller pol II subunits
ELOF1
SPT4
SDS-PAGE, SyproRuby stained
B
Activated elongation complex + IWS1
CTR9, IWS1
C
Activated elongation complex + IWS1ΔN
WDR61, RPB3,
IWS1ΔN
D
Activated elongation complex + IWS1ΔC
IWS1ΔC
E
Activated elongation complex + IWS1 HEAT/TND
HEAT/TND
F
Activated elongation complex + IWS1 HEAT/TND
Western blot, α-IWS1
Western blot, α-CTD

**Figure S3. Full size exclusion chromatograms and SDS-PAGE related to Figure 1.** UV 280 nm and UV 260 nm traces for size exclusion chromatograms and corresponding SDS-PAGE of elongation complexes assembled **(A)** without IWS1, **(B)** with full-length IWS1, **(C)** with IWS1ΔN, **(D)** with IWS1ΔC, and **(E)** with IWS1 HEAT/TND (551-699). Activated elongation complex = Pol II, DSIF, SPT6, PAF1c without RTF1, ELOF1. Fractions were loaded onto 4-12% Bis-Tris SDS-PAGE gels, run in 1x MES buffer, and stained with SyproRuby. **(F)** Western blots of size exclusion fractions in (E) for activated elongation complex assembled with IWS1 HEAT/TND.

### Figure S4

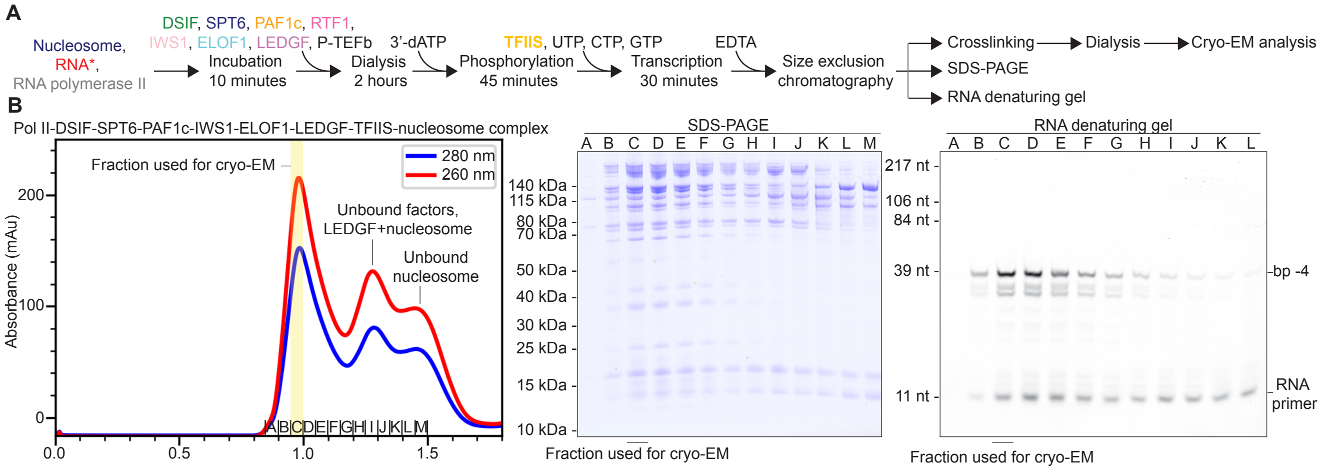

A
Nucleosome,
RNA*,
RNA polymerase II
DSIF, SPT6, PAF1c, RTF1,
IWS1, ELOF1, LEDGF, P-TEFb
Incubation
10 minutes
3'-dATP
Dialysis
2 hours
TFIIS, UTP, CTP, GTP
Phosphorylation
45 minutes
Transcription
30 minutes
EDTA
Size exclusion
chromatography
Crosslinking
Dialysis
Cryo-EM analysis
SDS-PAGE
RNA denaturing gel
B
Pol II-DSIF-SPT6-PAF1c-IWS1-ELOF1-LEDGF-TFIIS-nucleosome complex
Fraction used for cryo-EM
280 nm
260 nm
Unbound factors,
LEDGF+nucleosome
Unbound
nucleosome
Absorbance (mAu)
SDS-PAGE
140 kDa
115 kDa
80 kDa
70 kDa
50 kDa
40 kDa
30 kDa
25 kDa
15 kDa
10 kDa
Fraction used for cryo-EM
RNA denaturing gel
217 nt
106 nt
84 nt
39 nt
11 nt
bp -4
RNA
primer
Fraction used for cryo-EM

**Figure S4. Complex formation for cryo-EM. (A)** Reaction scheme for complex formation for cryo-EM sample. **(B)** Size-exclusion chromatogram, SDS-PAGE, and TBE-urea denaturing PAGE of Pol II-DSIF-SPT6-PAF1c-IWS1-ELOF1-LEDGF-TFIIS-nucleosome complex. Fractions represented by the letters A-M. Fraction used for cryo-EM indicated by yellow bar on chromatogram and black bar on gels.

Figure S5

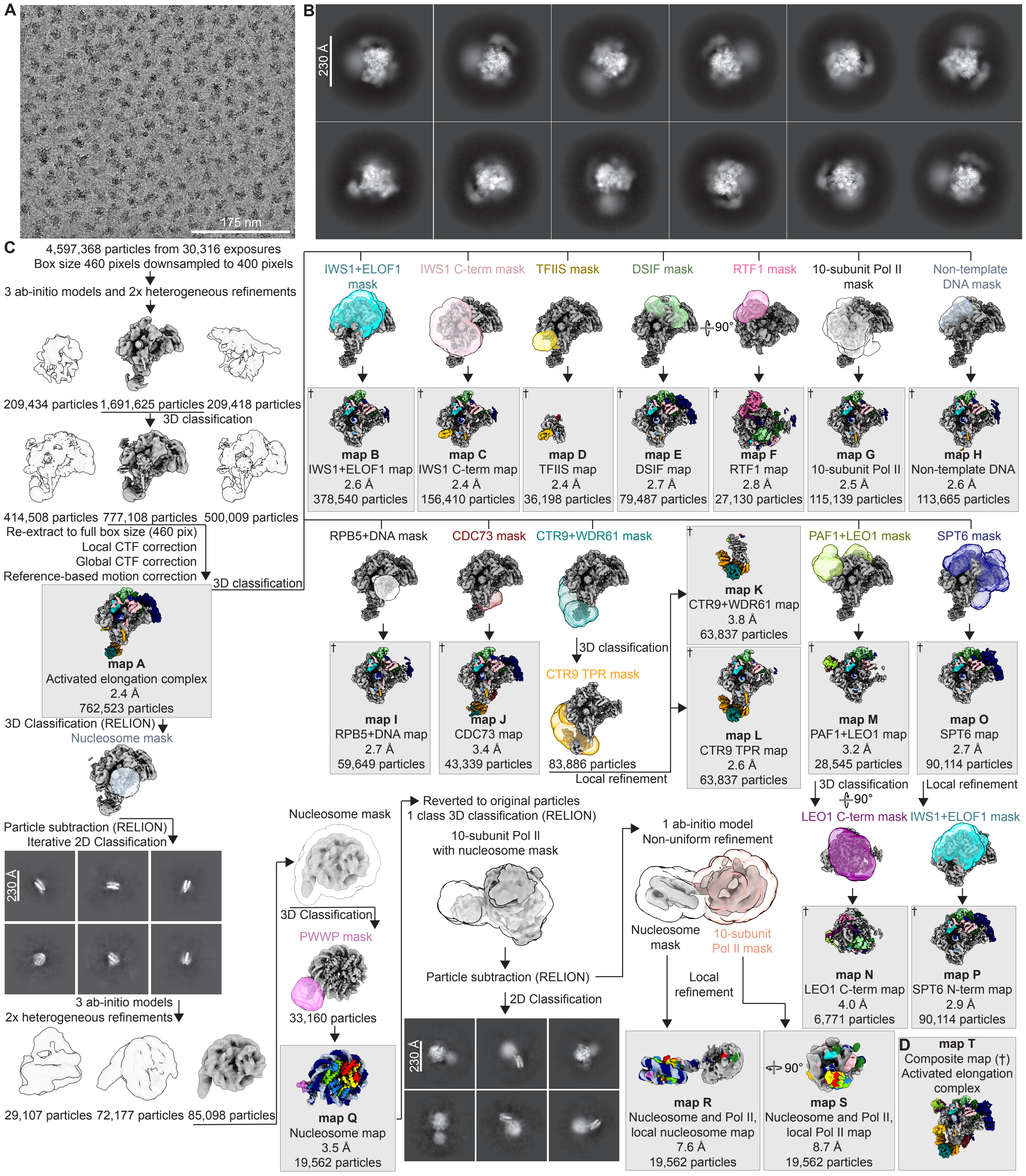
A
175 nm
B
230 Å
C
4,597,368 particles from 30,316 exposures
Box size 460 pixels downsampled to 400 pixels
3 ab-initio models and 2x heterogeneous refinements
209,434 particles
1,691,625 particles
209,418 particles
3D classification
414,508 particles
777,108 particles
500,009 particles
Re-extract to full box size (460 pix)
Local CTF correction
Global CTF correction
Reference-based motion correction
3D classification
map A
Activated elongation complex
2.4 Å
762,523 particles
3D Classification (RELION)
Nucleosome mask
Particle subtraction (RELION)
Iterative 2D Classification
230 Å
3 ab-initio models
2x heterogeneous refinements
29,107 particles
72,177 particles
85,098 particles
IWS1+ELOF1 mask
IWS1 C-term mask
TFIIS mask
DSIF mask
RTF1 mask
90°
10-subunit Pol II mask
Non-template DNA mask
map B
IWS1+ELOF1 map
2.6 Å
378,540 particles
map C
IWS1 C-term map
2.4 Å
156,410 particles
map D
TFIIS map
2.4 Å
36,198 particles
map E
DSIF map
2.7 Å
79,487 particles
map F
RTF1 map
2.8 Å
27,130 particles
map G
10-subunit Pol II
2.5 Å
115,139 particles
map H
Non-template DNA
2.6 Å
113,665 particles
RPB5+DNA mask
CDC73 mask
CTR9+WDR61 mask
map K
CTR9+WDR61 map
3.8 Å
63,837 particles
PAF1+LEO1 mask
SPT6 mask
3D classification
CTR9 TPR mask
map I
RPB5+DNA map
2.7 Å
59,649 particles
map J
CDC73 map
3.4 Å
43,339 particles
83,886 particles
Local refinement
map L
CTR9 TPR map
2.6 Å
63,837 particles
map M
PAF1+LEO1 map
3.2 Å
28,545 particles
map O
SPT6 map
2.7 Å
90,114 particles
3D classification
90°
Local refinement
LEO1 C-term mask
IWS1+ELOF1 mask
Nucleosome mask
Reverted to original particles
1 class 3D classification (RELION)
10-subunit Pol II
with nucleosome mask
1 ab-initio model
Non-uniform refinement
Nucleosome mask
10-subunit Pol II mask
3D Classification
PWWP mask
33,160 particles
Particle subtraction (RELION)
Local refinement
2D Classification
230 Å
map N
LEO1 C-term map
4.0 Å
6,771 particles
map P
SPT6 N-term map
2.9 Å
90,114 particles
map Q
Nucleosome map
3.5 Å
19,562 particles
map R
Nucleosome and Pol II,
local nucleosome map
7.6 Å
19,562 particles
90°
map S
Nucleosome and Pol II,
local Pol II map
8.7 Å
19,562 particles
D
map T
Composite map (†)
Activated elongation
complex

**Figure S5. Cryo-EM data processing of Pol II-DSIF-SPT6-PAF1c-IWS1-ELOF1-LEDGF-TFIIS-nucleosome complex. (A)** Representative micrograph from data collection with scale bar (175 nm). **(B)** Representative 2D classes with scale bar (230 Å). **(C)** Sorting and classification of cryo-EM data. Resolution and particle count indicated for each final map. Resolutions reported for tight mask values after FSC-mask auto-tightening. All processing performed in cryoSPARC unless indicated. Densities colored according to Figure 2. Maps contributing to composite map indicated with dagger symbol (†). **(D)** Composite map of Pol II-DSIF-SPT6-PAF1c-IWS1-ELOF1-TFIIS activated elongation complex. Map was created using Frankenmap of maps B, C, D, E, F, G, H, I, J, K, L, M, N, O, and P using local focus maps constructed from the atomic model. Generation of local focus maps detailed in Material and Methods.

### Figure S6

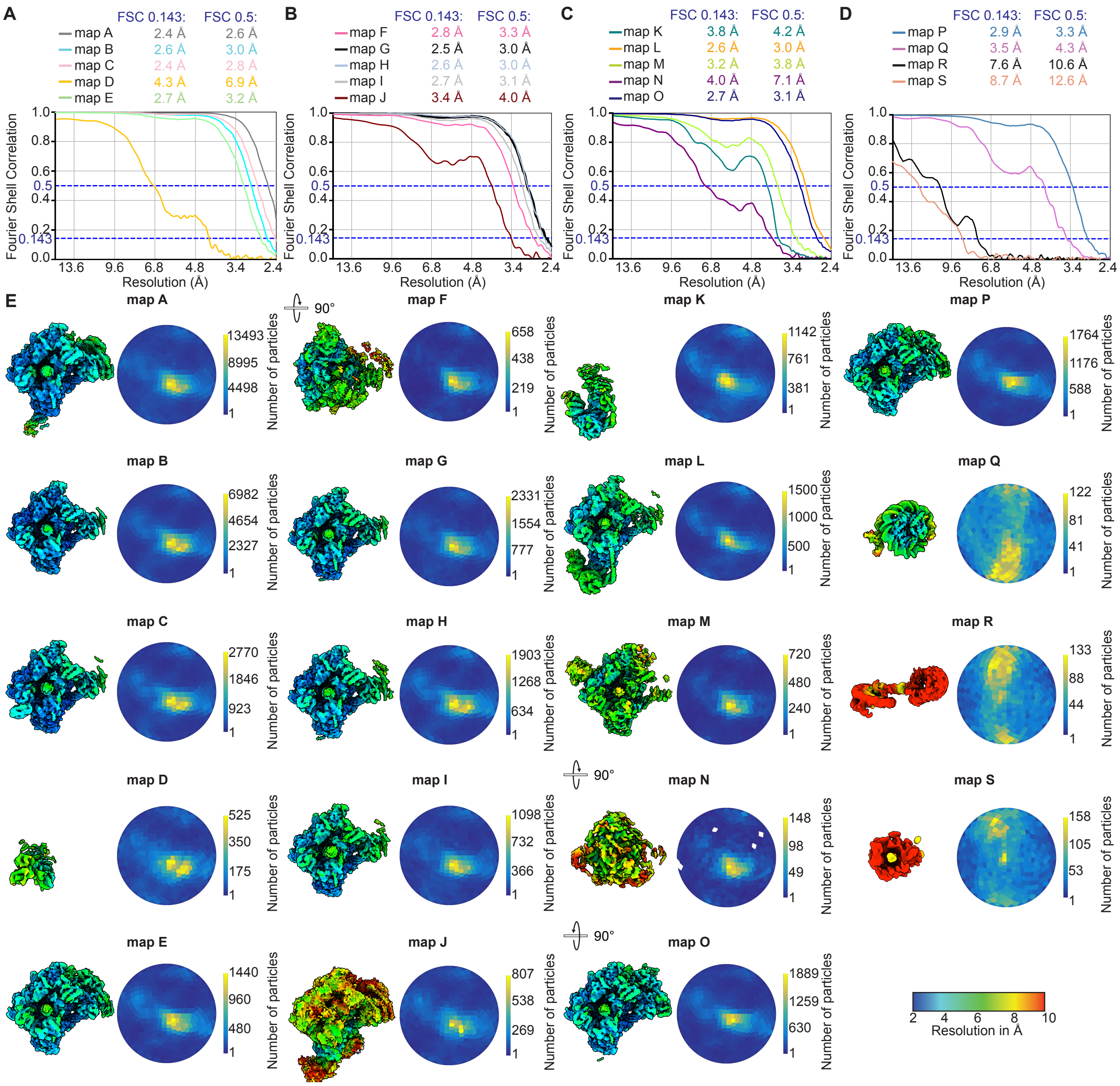
A
FSC 0.143: FSC 0.5:
map A 2.4 Å 2.6 Å
map B 2.6 Å 3.0 Å
map C 2.4 Å 2.8 Å
map D 4.3 Å 6.9 Å
map E 2.7 Å 3.2 Å
B
map F 2.8 Å 3.3 Å
map G 2.5 Å 3.0 Å
map H 2.6 Å 3.0 Å
map I 2.7 Å 3.1 Å
map J 3.4 Å 4.0 Å
C
map K 3.8 Å 4.2 Å
map L 2.6 Å 3.0 Å
map M 3.2 Å 3.8 Å
map N 4.0 Å 7.1 Å
map O 2.7 Å 3.1 Å
D
map P 2.9 Å 3.3 Å
map Q 3.5 Å 4.3 Å
map R 7.6 Å 10.6 Å
map S 8.7 Å 12.6 Å
Fourier Shell Correlation
Resolution (Å)
E
Number of particles
90°
Resolution in Å

**Figure S6. Data quality and metrics of Pol II-DSIF-SPT6-PAF1c-IWS1-ELOF1-LEDGF-TFIIS-nucleosome complex.** FSC curves for maps **(A)** A-E, **(B)** F-J, **(C)** K-O, and **(D)** P-S. Resolution at FSC 0.5 and 0.143 reported for tight mask values after FSC-mask auto-tightening and color-coded according to map. **(D)** Local resolution estimation and particle angular distribution plots for each map.

Figure S7

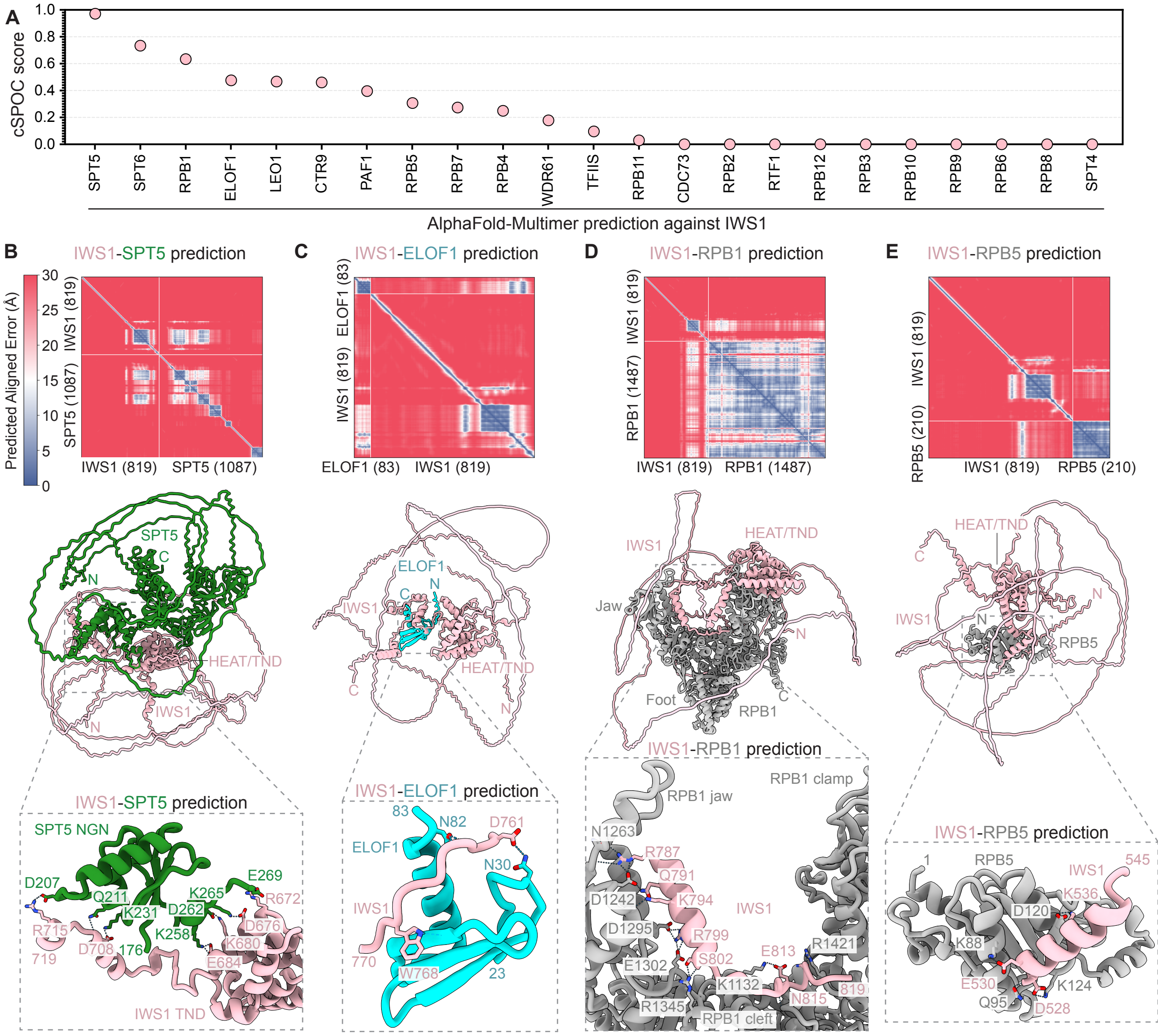

A
cSPOC score
1.0
0.8
0.6
0.4
0.2
0.0
SPT5
SPT6
RPB1
ELOF1
LEO1
CTR9
PAF1
RPB5
RPB7
RPB4
WDR61
TFIIS
RPB11
CDC73
RPB2
RTF1
RPB12
RPB3
RPB10
RPB9
RPB6
RPB8
SPT4
AlphaFold-Multimer prediction against IWS1
B
IWS1-SPT5 prediction
Predicted Aligned Error (Å)
IWS1 (819)
SPT5 (1087)
C
IWS1-ELOF1 prediction
ELOF1 (83)
IWS1 (819)
D
IWS1-RPB1 prediction
RPB1 (1487)
E
IWS1-RPB5 prediction
RPB5 (210)
HEAT/TND
IWS1
N
C
Jaw
Foot
RPB1
RPB5
SPT5 NGN
D207
Q211
K231
D262
K265
E269
R672
D676
K680
E684
R715
719
D708
176
K258
IWS1 TND
83
N82
D761
N30
770
W768
23
RPB1 jaw
RPB1 clamp
N1263
R787
Q791
D1242
K794
D1295
R799
S802
E1302
E813
R1421
K1132
R1345
RPB1 cleft
N815
819
1
545
K536
D120
K88
E530
Q95
K124
D528

**Figure S7. AlphaFold-Multimer screen of IWS1 with core human elongation factors.** **(A)** Interaction screen using AlphaFold-Multimer. Predictions ranked by cSPOC score. **(B)** PAE plot of IWS1-SPT5 prediction and representative predicted model with IWS1-SPT5 NGN contacts shown. Interacting atoms shown as sticks and hydrogen bonds (H-bonds) shown as dotted lines. **(C)** PAE plot of IWS1-ELOF1 prediction and representative predicted model shown. Interacting atoms shown as sticks and H-bonds shown as dotted lines. **(D)** PAE plot of IWS1-RPB1 prediction and representative predicted model with IWS1-RPB1 jaw and cleft contacts shown. Interacting atoms shown as sticks and H-bonds shown as dotted lines. **(E)** PAE plot of IWS1-RPB5 prediction and representative predicted model shown. Interacting atoms shown as sticks and H-bonds shown as dotted lines.

### Figure S8

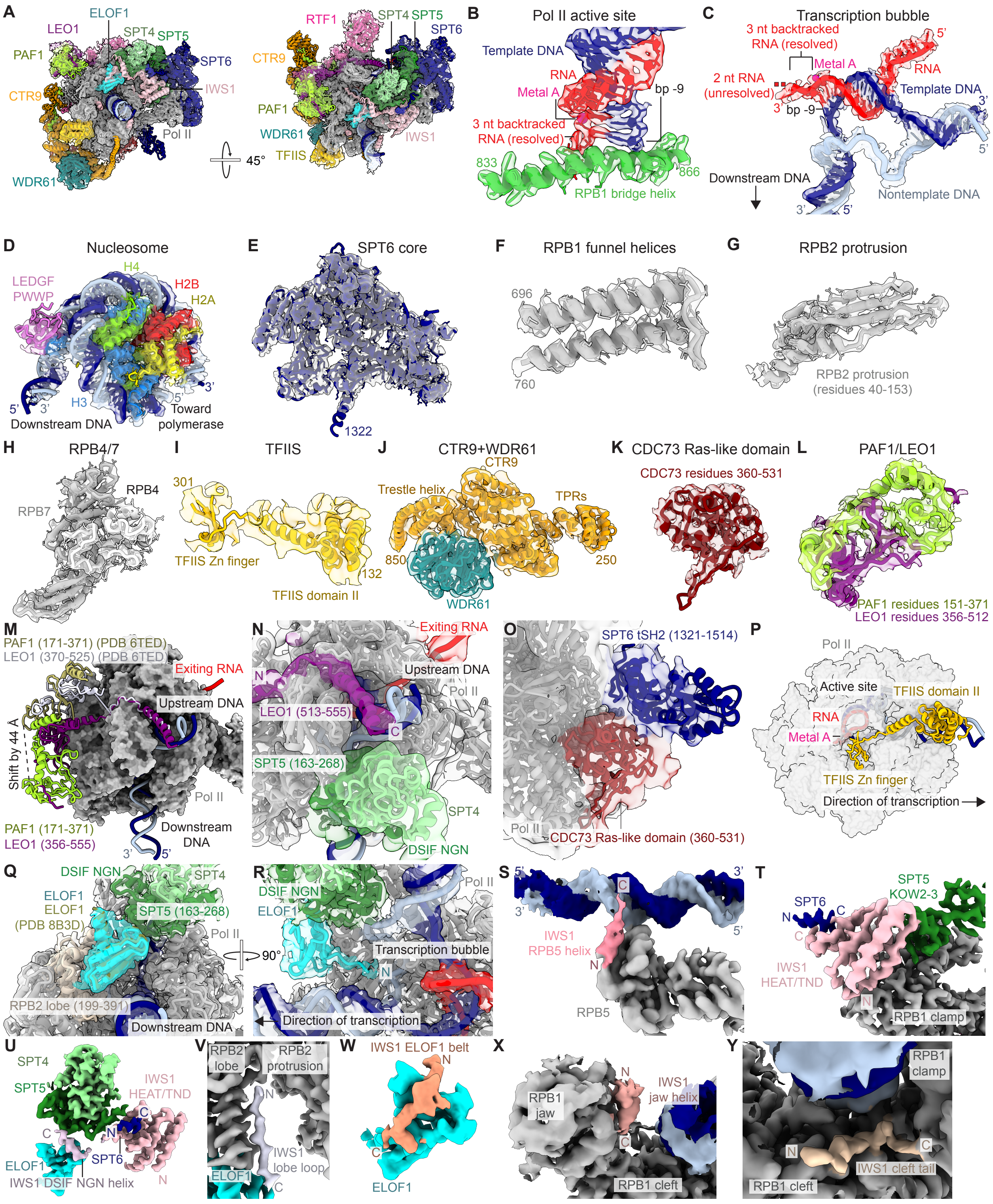

A
ELOF1
LEO1
SPT4
SPT5
PAF1
SPT6
CTR9
IWS1
Pol II
WDR61
45°
RTF1
SPT4
SPT5
SPT6
CTR9
PAF1
WDR61
TFIIS
IWS1
B
Pol II active site
Template DNA
RNA
Metal A
bp -9
3 nt backtracked
RNA (resolved)
833
866
RPB1 bridge helix
C
Transcription bubble
3 nt backtracked
RNA (resolved)
5'
Metal A
RNA
2 nt RNA
(unresolved)
Template DNA
3'
bp -9
3'
5'
Downstream DNA
Nontemplate DNA
3'
5'
D
Nucleosome
H4
LEDGF
PWWP
H2B
H2A
3'
5'
3'
H3
5'
Downstream DNA
Toward
polymerase
E
SPT6 core
1322
F
RPB1 funnel helices
696
760
G
RPB2 protrusion
RPB2 protrusion
(residues 40-153)
H
RPB4/7
RPB4
RPB7
I
TFIIS
301
TFIIS Zn finger
132
TFIIS domain II
J
CTR9+WDR61
CTR9
Trestle helix
TPRs
850
250
WDR61
K
CDC73 Ras-like domain
CDC73 residues 360-531
L
PAF1/LEO1
PAF1 residues 151-371
LEO1 residues 356-512
M
PAF1 (171-371) (PDB 6TED)
LEO1 (370-525) (PDB 6TED)
Exiting RNA
Upstream DNA
Shift by 44 Å
Pol II
PAF1 (171-371)
LEO1 (356-555)
Downstream
DNA
3'
5'
N
Exiting RNA
N
Upstream DNA
Pol II
LEO1 (513-555)
C
SPT5 (163-268)
SPT4
DSIF NGN
O
SPT6 tSH2 (1321-1514)
Pol II
CDC73 Ras-like domain (360-531)
P
Pol II
Active site
TFIIS domain II
RNA
Metal A
TFIIS Zn finger
Direction of transcription
Q
DSIF NGN
SPT4
ELOF1
ELOF1
(PDB 8B3D)
SPT5 (163-268)
Pol II
90°
RPB2 lobe (199-391)
Downstream DNA
R
DSIF NGN
Pol II
ELOF1
Transcription bubble
N
Direction of transcription
S
5'
3'
C
3'
5'
IWS1
RPB5 helix
N
RPB5
T
SPT5
KOW2-3
SPT6
N
C
C
IWS1
HEAT/TND
N
RPB1 clamp
U
SPT4
SPT5
IWS1
HEAT/TND
C
C
N
SPT6
ELOF1
IWS1 DSIF NGN helix
N
V
RPB2
lobe
RPB2
protrusion
N
IWS1
lobe loop
ELOF1
C
W
IWS1 ELOF1 belt
N
C
ELOF1
X
RPB1
jaw
N
IWS1
jaw helix
C
RPB1 cleft
Y
RPB1
clamp
N
C
IWS1 cleft tail
RPB1 cleft

**Figure S8. Structural validation and cryo-EM densities of Pol II-DSIF-SPT6-PAF1c-IWS1-ELOF1-LEDGF-TFIIS-nucleosome complex. (A)** Two views of transparent composite map T (Frankenmap) of Pol II-DSIF-SPT6-PAF1c-IWS1-ELOF1-TFIIS activated elongation complex overlaid with atomic model. **(B)** RPB1 bridge helix, RNA, and template DNA of Pol II active site atoms overlaid on map H. Pol II is stalled at bp -9 with 3 nucleotides of backtracked RNA resolved. **(C)** RNA, template, and nontemplate DNA cartoons and atoms overlaid on map H. Single-stranded nontemplate backbone shown as cartoon. 3 nucleotides of backtracked RNA are resolved with 2 nucleotides of backtracked RNA unresolved. **(D)** LEDGF PWWP (cartoons) bound to promoter distal H3 and transcribed nucleosome (atoms) overlaid on map Q. **(E)** SPT6 core (residues 267-1322) cartoons overlaid on map O. **(F)** RPB1 funnel helices (residues 696-760) cartoons and atoms overlaid on map A. **(G)** $\beta$-sheets of RPB2 protrusion domain (residues 40-153) cartoons and atoms overlaid on map E. **(H)** RPB4 and RPB7 cartoons and atoms overlaid on map O. **(I)** TFIIS domain II, helix, and Zn finger shown as cartoons overlaid on map D. **(J)** CTR9 residues 250-850 encompassing TPRs and part of the trestle helix and WDR61 cartoons overlaid on map K. **(K)** CDC73 Ras-like domain cartoons overlaid on map J. **(L)** PAF1/LEO1 dimer cartoons overlaid on map M. **(M)** PAF1 and LEO1 from activated elongation complex model (PDB 6TED) overlaid on PAF1 and LEO1 model (this study) shown as cartoons. Pol II shown in surface representation. Distance of 44 Å measured between $\beta$ carbon of Asp 285 on PAF1 models. **(N)** LEO1 C-terminal helix positioned next to upstream DNA and DSIF NGN shown as cartoons. Model overlaid on map N which was gaussian-filtered to 2 standard deviations. **(O)** Positions of SPT6 tSH2 domain and CDC73 Ras-like domain with respect to Pol II shown as cartoons. Model overlaid on map J which was gaussian-filtered to 2 standard deviations. **(P)** TFIIS domain II, helix, and Zn finger position with respect to Pol II active site. Pol II shown as transparent surface. **(Q)** ELOF1 from transcription coupled nucleotide excision repair complex (PDB 8B3D) overlaid on ELOF1 model (this study) and map B. ELOF1 position relative to DSIF NGN and RPB2 lobe shown. **(R)** ELOF1 N-terminus reaches toward transcription bubble. Model overlaid on map B. **(S)** IWS1 interaction with RPB5 colored on map I. **(T)** IWS1 HEAT/TND and interactions with SPT5 and SPT6 colored on map B. **(U)** IWS1 interaction with DSIF NGN and ELOF1 colored on map B. **(V)** IWS1 interaction with RPB2 lobe colored on map B. **(W)** IWS1 interaction with ELOF1 colored on map B. **(X)** IWS1 interaction with RPB1 jaw colored on map C. **(Y)** IWS1 interaction with RPB1 cleft colored on map C.

Figure S9

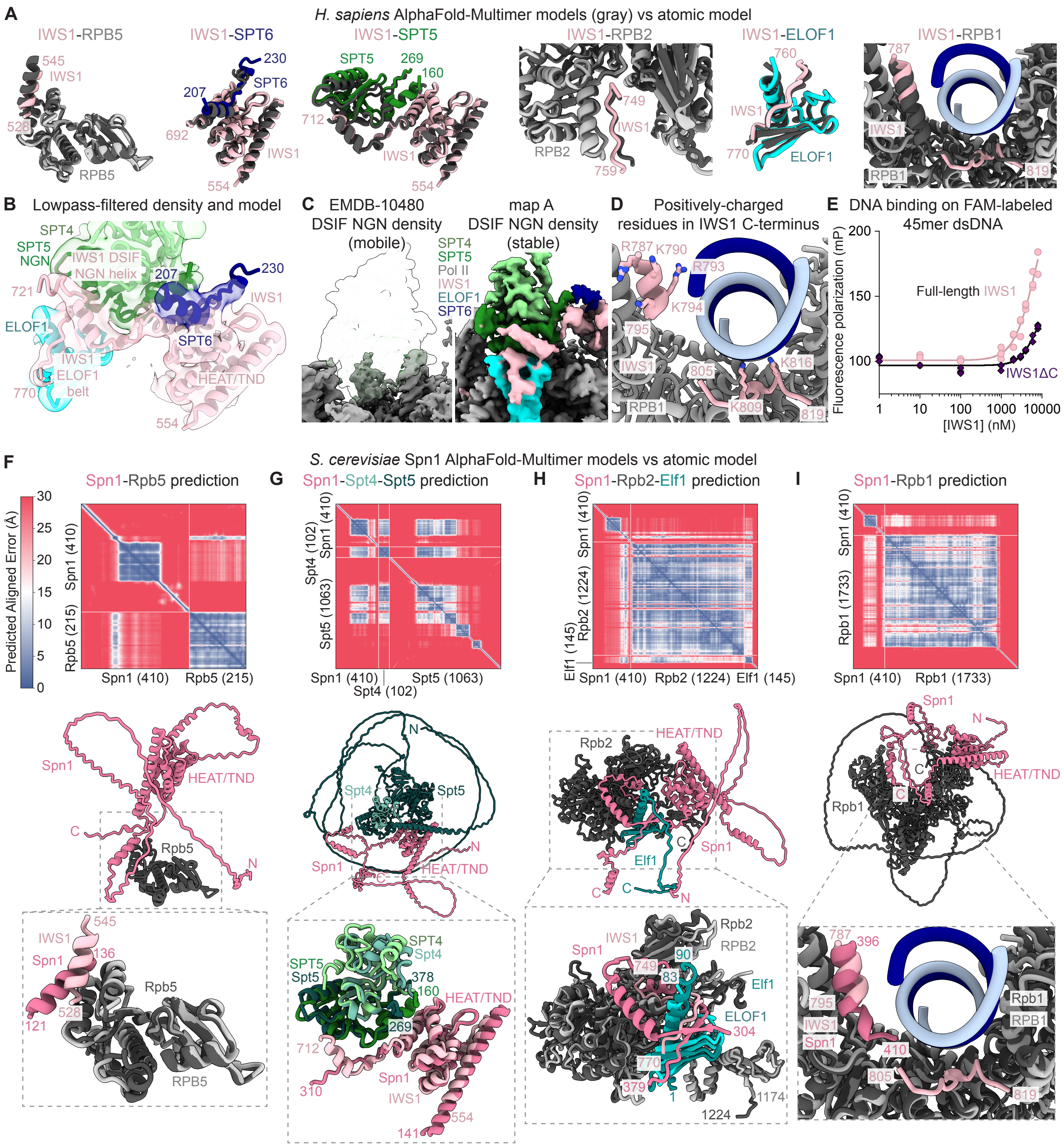
A
H. sapiens AlphaFold-Multimer models (gray) vs atomic model
IWS1-RPB5
IWS1-SPT6
IWS1-SPT5
IWS1-RPB2
IWS1-ELOF1
IWS1-RPB1
B
Lowpass-filtered density and model
C
EMDB-10480 DSIF NGN density (mobile)
map A DSIF NGN density (stable)
D
Positively-charged residues in IWS1 C-terminus
E
DNA binding on FAM-labeled 45mer dsDNA
Fluorescence polarization (mP)
Full-length IWS1
IWS1ΔC
[IWS1] (nM)
F
S. cerevisiae Spn1 AlphaFold-Multimer models vs atomic model
Spn1-Rpb5 prediction
Predicted Aligned Error (Å)
G
Spn1-Spt4-Spt5 prediction
H
Spn1-Rpb2-Elf1 prediction
I
Spn1-Rpb1 prediction

**Figure S9. Comparisons of AlphaFold models to atomic model and additional structural details. (A)** AlphaFold-Multimer predicted models (gray) for IWS1-RPB5, IWS1-SPT6, IWS1-SPT5, IWS1-RPB2, IWS1-ELOF1, and IWS1-RPB1 interactions overlaid on atomic model. **(B)** Atomic model overlaid on map B gaussian-filtered to 1.5 standard deviation. Density from gaussian-filtered map allows placement of SPT6 residues 219-230 and IWS1 residues 713-721. **(C)** DSIF NGN density comparison between Pol II-DSIF-PAF-SPT6 activated elongation complex EMDB-10480 and map A. Density threshold set based on similarity of visible features of RNA polymerase II. **(D)** Positively charged residues (R787, K790, R793, K794, K809, K816) in IWS1 RPB1 jaw helix and RPB1 cleft tail shown as sticks. **(E)** Fluorescence DNA polarization assays of wild-type IWS1 and IWS1ΔC. Fluorescence polarization (mP) was measured for a 10 nM FAM-labeled 45mer dsDNA substrate. Nonlinear regression curve fit to data points. IC50 for IWS1 and IWS1ΔC calculated as 6580 and 7398, respectively. **(F)** PAE plot of *S. cerevisiae* Spn1-Rpb5 AlphaFold-Multimer prediction shown and representative predicted model overlaid with atomic model of human IWS1 and RPB5 from this study. **(G)** PAE plot of *S. cerevisiae* Spn1-Spt4-Spt5 AlphaFold-Multimer prediction shown and representative predicted model overlaid with atomic model of human IWS1, SPT4, and SPT5 from this study. **(H)** PAE plot of *S. cerevisiae* Spn1-Rpb2-Elf1 AlphaFold-Multimer prediction shown and representative predicted model overlaid with atomic model of human IWS1, RPB2, and ELOF1 from this study. **(I)** PAE plot of *S. cerevisiae* Spn1-Rpb1 AlphaFold-Multimer prediction shown and representative predicted model overlaid with atomic model of human IWS1 and RPB1 from this study.

Figure S10

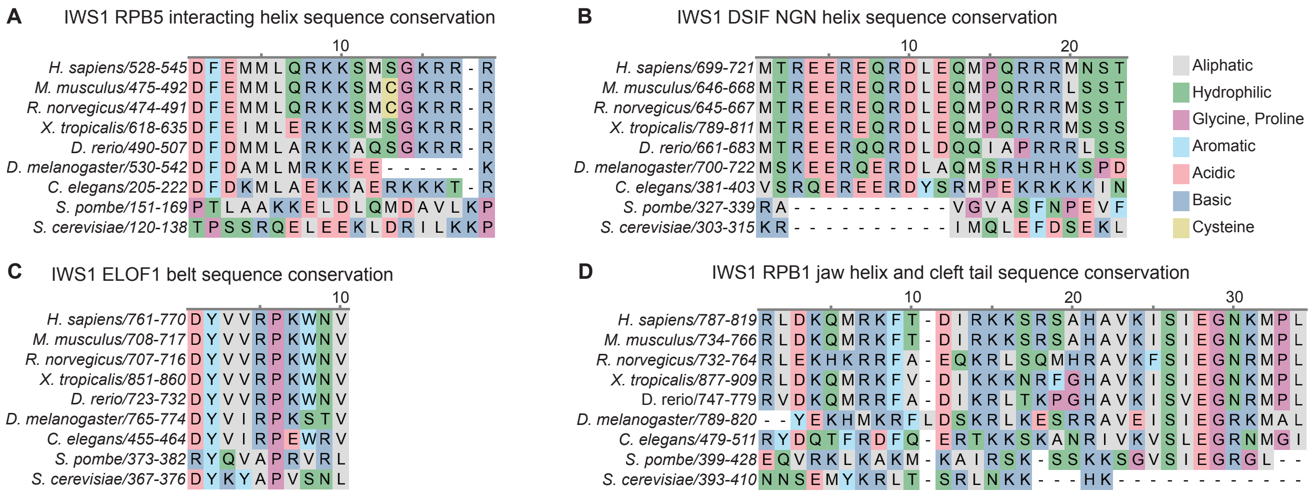
A
IWS1 RPB5 interacting helix sequence conservation
10
H. sapiens/528-545 DFEMMLQRKKSMSGKRR-R
M. musculus/475-492 DFEMMLQRKKSMCGKRR-R
R. norvegicus/474-491 DFEMMLQRKKSMCGKRR-R
X. tropicalis/618-635 DFEIMLERKKSMSGKRR-R
D. rerio/490-507 DFDMMLARKKAQSGKRR-R
D. melanogaster/530-542 DFDAMLMRKKEE------K
C. elegans/205-222 DFDKMLAEKKAERKKKT-R
S. pombe/151-169 PTLAAKKELDLQMDAVLKP
S. cerevisiae/120-138 TPSSRQELEEKLDRILKKP
B
IWS1 DSIF NGN helix sequence conservation
10
20
H. sapiens/699-721 MTREEREQRDLEQMPQRRRMNST
M. musculus/646-668 MTREEREQRDLEQMPQRRRLSST
R. norvegicus/645-667 MTREEREQRDLEQMPQRRRMSST
X. tropicalis/789-811 MTREEREQRDLEQMPQRRRMSSS
D. rerio/661-683 MTREERQQRDLDQQIAPRRRLSS
D. melanogaster/700-722 MSKEERQERDLAQMSRHRHKSPD
C. elegans/381-403 VSRQEREERDYSRMPEKRKKKIN
S. pombe/327-339 RA----------VGVASFNPEVF
S. cerevisiae/303-315 KR----------IMQLEFDSEKL
Aliphatic
Hydrophilic
Glycine, Proline
Aromatic
Acidic
Basic
Cysteine
C
IWS1 ELOF1 belt sequence conservation
10
H. sapiens/761-770 DYVVRPKWNV
M. musculus/708-717 DYVVRPKWNV
R. norvegicus/707-716 DYVVRPKWNV
X. tropicalis/851-860 DYVVRPKWNV
D. rerio/723-732 DYVVRPKWNV
D. melanogaster/765-774 DYVIRPKSTV
C. elegans/455-464 DYVIRPEWRV
S. pombe/373-382 RYQVAPRVRL
S. cerevisiae/367-376 DYKYAPVSNL
D
IWS1 RPB1 jaw helix and cleft tail sequence conservation
10
20
30
H. sapiens/787-819 RLDKQMRKFT-DIRKKSRSAHAVKISIEGNKMPL
M. musculus/734-766 RLDKQMRKFT-DIRKKSRSAHAVKISIEGNKMPL
R. norvegicus/732-764 RLEKHKRRFA-EQKRLSQMHRAVKFSIEGNRMPL
X. tropicalis/877-909 RLDKQMRKFV-DIKKKNRFGHAVKISIEGNKMPL
D. rerio/747-779 RVDKQMRRFA-DIKRLTKPGHAVKISVEGNRMPL
D. melanogaster/789-820 --YEKHMKRFLDSKRLKESRRAVEISIEGRKMAL
C. elegans/479-511 RYDQTFRDFQ-ERTKKSKANRIVKVSLEGRNMGI
S. pombe/399-428 EQVRKLKAKM-KAIRSK-SSKKSGVSIEGRGL--
S. cerevisiae/393-410 NNSEMYKRLT-SRLNKK---HK------------

**Figure S10. IWS1 sequence alignments.** PROMALS3D multiple sequence alignments demonstrate high conservation of **(A)** IWS1 RPB5 helix (residues 528-545), **(B)** IWS1 DSIF NGN helix (699-721), **(C)** IWS1 ELOF1 belt (residues 761-770), and **(D)** IWS1 RPB1 jaw helix and cleft tail (residues 787-819).

Figure S11

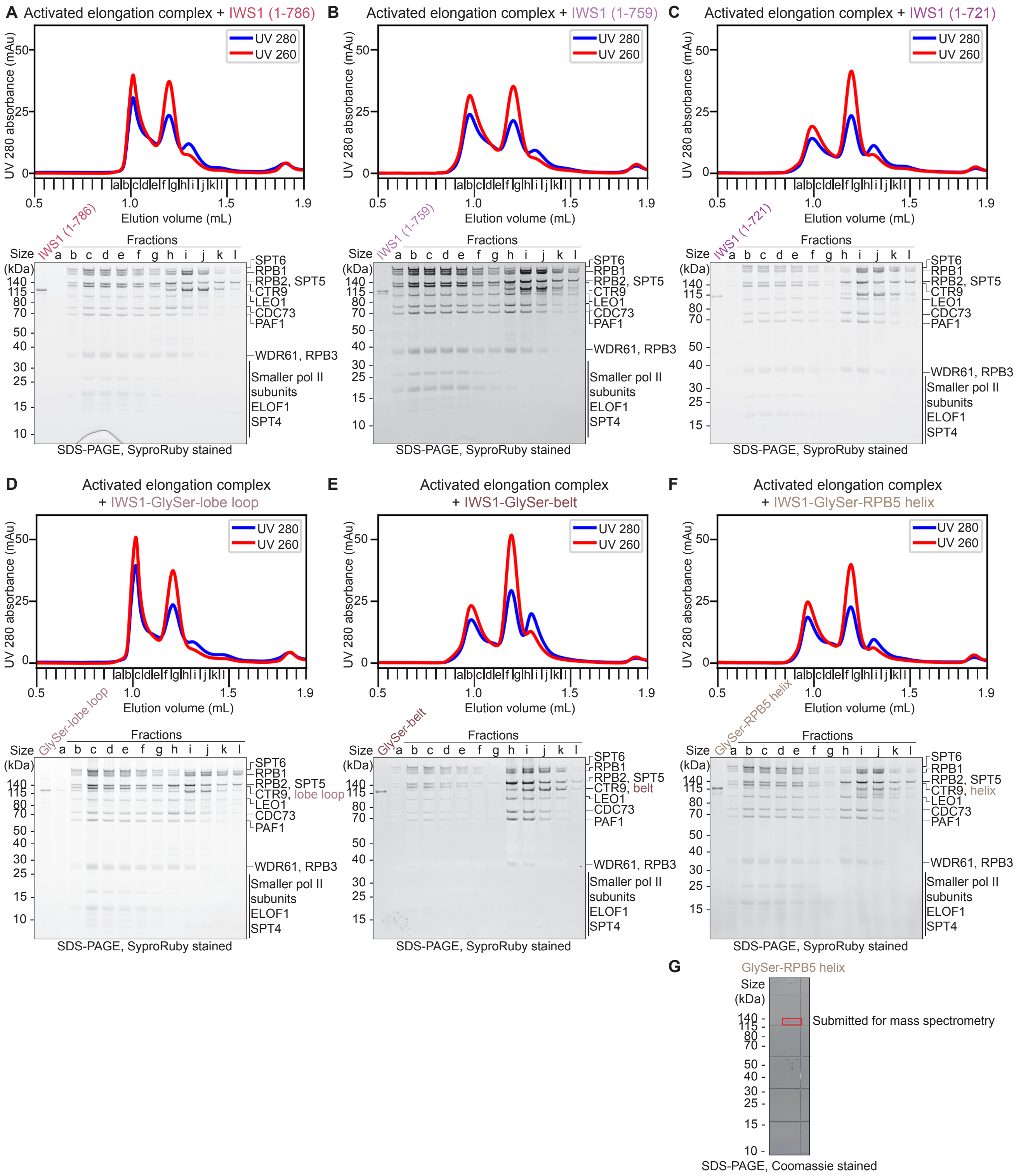

A Activated elongation complex + IWS1 (1-786)
B Activated elongation complex + IWS1 (1-759)
C Activated elongation complex + IWS1 (1-721)
D Activated elongation complex + IWS1-GlySer-lobe loop
E Activated elongation complex + IWS1-GlySer-belt
F Activated elongation complex + IWS1-GlySer-RPB5 helix
G GlySer-RPB5 helix
UV 280
UV 260
UV 280 absorbance (mAu)
Elution volume (mL)
Fractions
Size (kDa)
SPT6
RPB1
RPB2, SPT5
CTR9
LEO1
CDC73
PAF1
WDR61, RPB3
Smaller pol II subunits
ELOF1
SPT4
CTR9, lobe loop
CTR9, belt
CTR9, helix
SDS-PAGE, SyproRuby stained
Submitted for mass spectrometry
SDS-PAGE, Coomassie stained

**Figure S11. Full size exclusion chromatograms and SDS-PAGE related to Figure 4.** UV 280 nm and UV 260 nm traces for size exclusion chromatograms and corresponding SDS-PAGE of elongation complexes assembled with **(A)** IWS1 (1-786), **(B)** IWS1 (1-759), **(C)** IWS1 (1-721), **(D)** IWS1-GlySer-lobe loop, **(E)** IWS1-GlySer-belt, and **(F)** IWS1-GlySer-RPB5 helix. Activated elongation complex = Pol II, DSIF, SPT6, PAF1c without RTF1, ELOF1. Fractions were loaded onto 4-12% Bis-Tris SDS-PAGE gels, run in 1x MES buffer, and stained with SyproRuby. **(G)** Image of purified IWS1-GlySer-RPB5 helix run on 4-12% Bis-Tris SDS-PAGE gel and stained with OneStep Blue. Red box indicates region of gel excised for mass spectrometry.

Figure S12

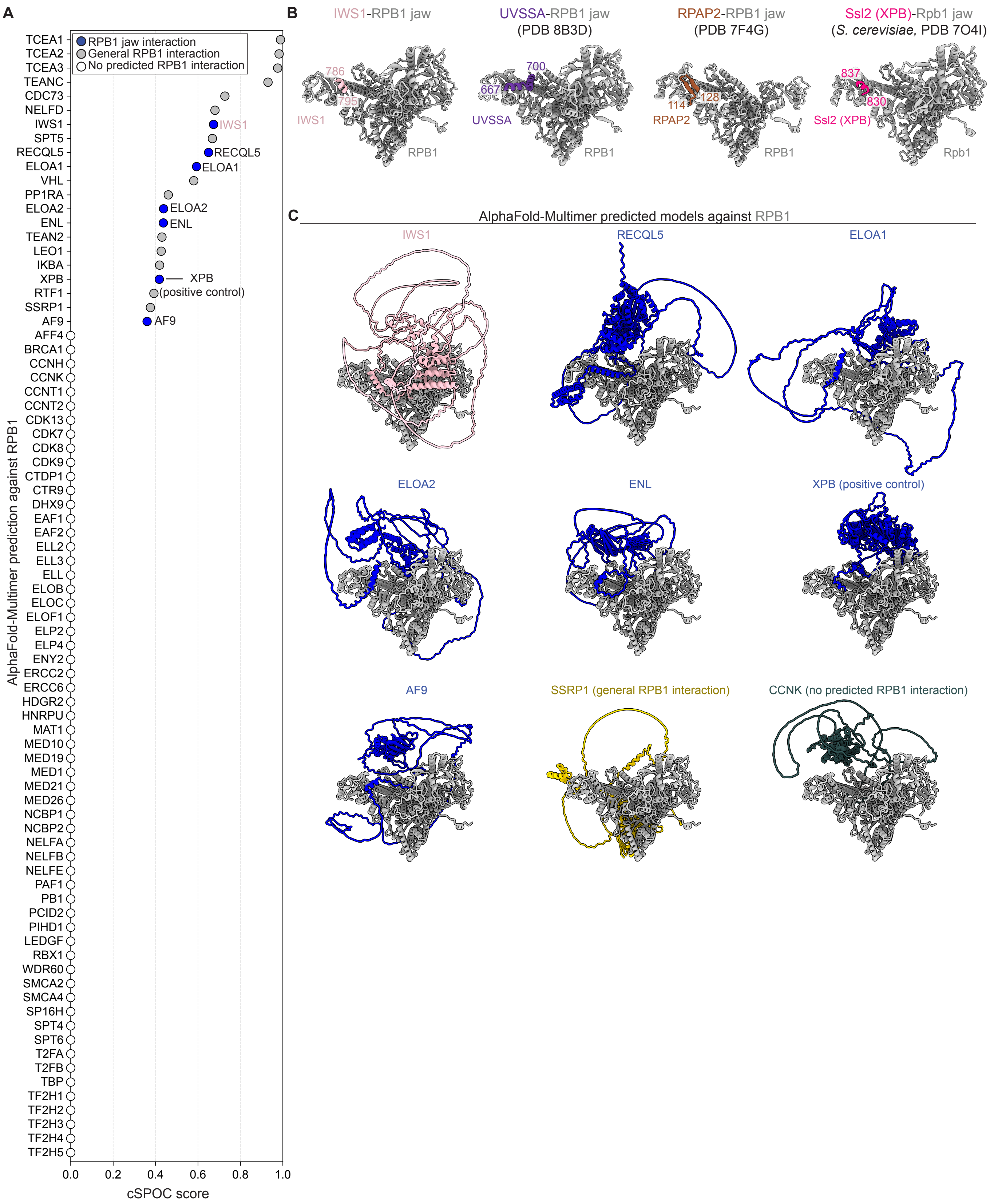

A
RPB1 jaw interaction
General RPB1 interaction
No predicted RPB1 interaction
TCEA1
TCEA2
TCEA3
TEANC
CDC73
NELFD
IWS1
SPT5
RECQL5
ELOA1
VHL
PP1RA
ELOA2
ENL
TEAN2
LEO1
IKBA
XPB
RTF1
SSRP1
AF9
AFF4
BRCA1
CCNH
CCNK
CCNT1
CCNT2
CDK13
CDK7
CDK8
CDK9
CTDP1
CTR9
DHX9
EAF1
EAF2
ELL2
ELL3
ELL
ELOB
ELOC
ELOF1
ELP2
ELP4
ENY2
ERCC2
ERCC6
HDGR2
HNRPU
MAT1
MED10
MED19
MED1
MED21
MED26
NCBP1
NCBP2
NELFA
NELFB
NELFE
PAF1
PB1
PCID2
PIHD1
LEDGF
RBX1
WDR60
SMCA2
SMCA4
SP16H
SPT4
SPT6
T2FA
T2FB
TBP
TF2H1
TF2H2
TF2H3
TF2H4
TF2H5
XPB (positive control)
AlphaFold-Multimer prediction against RPB1
0.0
0.2
0.4
0.6
0.8
1.0
cSPOC score
B
IWS1-RPB1 jaw
786
795
IWS1
RPB1
UVSSA-RPB1 jaw
(PDB 8B3D)
700
667
UVSSA
RPB1
RPAP2-RPB1 jaw
(PDB 7F4G)
114
128
RPAP2
RPB1
Ssl2 (XPB)-Rpb1 jaw
(S. cerevisiae, PDB 7O4I)
837
830
Ssl2 (XPB)
Rpb1
C
AlphaFold-Multimer predicted models against RPB1
IWS1
RECQL5
ELOA1
ELOA2
ENL
XPB (positive control)
AF9
SSRP1 (general RPB1 interaction)
CCNK (no predicted RPB1 interaction)

**Figure S12. RPB1 interaction screen. (A)** All human proteins screened against human RPB1 using AlphaFold-Multimer. Predictions ranked by cSPOC score. Representative RPB1 jaw interactors are shown as dark blue circles, general RPB1 interactions shown as grey circles, no predicted RPB1 interaction shown as white circles. **(B)** Models of known RPB1 jaw interactors, including IWS1 (this study), UVSSA (PDB 8B3D), RPAP2 (PDB 7F4G), and yeast Ssl2 (human XPB) (PDB 7O4I). **(C)** AlphaFold-Multimer predicted models of representative candidates with RPB1. RPB1 jaw interactors colored in blue with IWS1 colored in pink. Representative general RPB1 interaction SSRP1 colored in gold. Representative non-interacting example (CCNK) colored in dark slate gray.

Figure S13

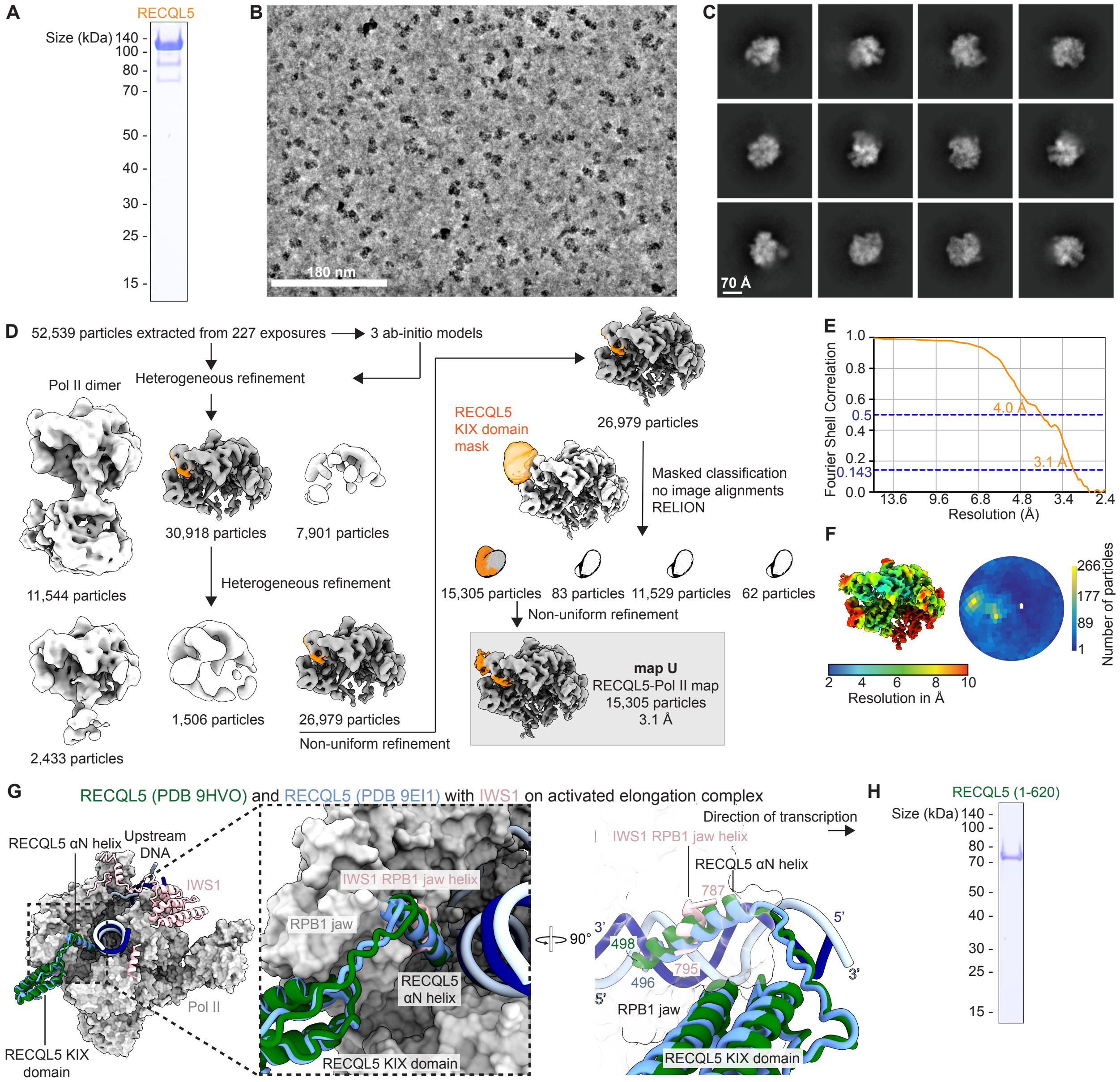
A
RECQL5
Size (kDa) 140 -
100 -
80 -
70 -
50 -
40 -
30 -
25 -
15 -
B
180 nm
C
70 Å
D
52,539 particles extracted from 227 exposures
3 ab-initio models
Heterogeneous refinement
Pol II dimer
11,544 particles
30,918 particles
7,901 particles
Heterogeneous refinement
2,433 particles
1,506 particles
26,979 particles
Non-uniform refinement
26,979 particles
RECQL5
KIX domain
mask
Masked classification
no image alignments
RELION
15,305 particles
83 particles
11,529 particles
62 particles
Non-uniform refinement
map U
RECQL5-Pol II map
15,305 particles
3.1 Å
E
Fourier Shell Correlation
1.0
0.8
0.6
0.5
0.4
0.2
0.143
0.0
4.0 Å
3.1 Å
13.6
9.6
6.8
4.8
3.4
2.4
Resolution (Å)
F
2
4
6
8
10
Resolution in Å
266
177
89
1
Number of particles
G
RECQL5 (PDB 9HVO) and RECQL5 (PDB 9EI1) with IWS1 on activated elongation complex
RECQL5 αN helix
Upstream DNA
IWS1
Pol II
RECQL5 KIX domain
IWS1 RPB1 jaw helix
RPB1 jaw
RECQL5 αN helix
RECQL5 KIX domain
90°
Direction of transcription
IWS1 RPB1 jaw helix
RECQL5 αN helix
787
3'
5'
498
795
496
3'
5'
RPB1 jaw
RECQL5 KIX domain
H
RECQL5 (1-620)
Size (kDa) 140 -
100 -
80 -
70 -
50 -
40 -
30 -
25 -
15 -

**Figure S13. RECQL5-Pol II complex cryo-EM data processing and construct used in RECQL5+IWS1 RNA extension assay. (A)** SDS-PAGE of purified RECQL5. Gel stained with OneStep Blue. **(B)** Representative denoised micrograph from RECQL5-Pol II sample cryo-EM data collection. Scale bar of 180 nm. **(C)** Representative 2D classes from data processing with scale bar of 70 Å. **(D)** Sorting and classification of cryo-EM data. All processing performed in cryoSPARC unless indicated. Densities colored according to model in Figure 5C. **(E)** FSC curve for RECQL5-Pol II map. Resolution at FSC 0.5 and 0.143 indicated. **(F)** Local resolution estimation and angular distribution plot for RECQL5-Pol II map. **(G)** Overlay of RECQL5 $\alpha$N helix and KIX domain from RECQL5-Pol II models (PDB 9HVO, PDB 9EI1) on activated elongation complex model (this study) shows binding overlap of RECQL5 $\alpha$N helix and IWS1 RPB1 jaw helix. Pol II model shown as surfaces and IWS1 model shown as cartoons. **(H)** SDS-PAGE of purified RECQL5 (1-620). Gel stained with OneStep Blue.

Figure S14

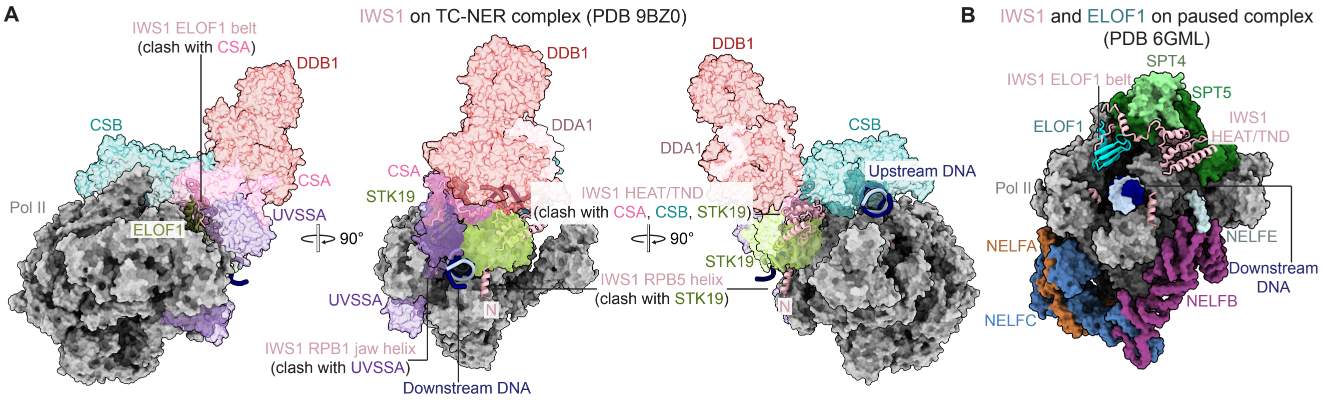
A
IWS1 on TC-NER complex (PDB 9BZ0)
IWS1 ELOF1 belt
(clash with CSA)
DDB1
CSB
CSA
Pol II
UVSSA
ELOF1
90°
DDA1
STK19
IWS1 HEAT/TND
(clash with CSA, CSB, STK19)
UVSSA
IWS1 RPB5 helix
(clash with STK19)
IWS1 RPB1 jaw helix
(clash with UVSSA)
Downstream DNA
N
Upstream DNA
B
IWS1 and ELOF1 on paused complex
(PDB 6GML)
SPT4
SPT5
IWS1 HEAT/TND
NELFE
NELFA
NELFB
NELFC
Downstream DNA

**Figure S14. IWS1 and ELOF1 compatibility with known transcription complexes.** **(A)** Three views of IWS1 model shown as cartoons overlaid on TC-NER+STK19 complex (PDB 9BZ0). 9BZ0 ELOF1 shown in cartoons and the remaining TC-NER+STK19 factors shown as transparent surfaces. Clashes are indicated: IWS1 RPB5 helix with STK19, IWS1 HEAT/TND with CSA, CSB, and STK19, IWS1 ELOF1 belt with CSA, and IWS1 RPB1 jaw helix with UVSSA. **(B)** IWS1 and ELOF1 models shown as cartoons and overlaid on mammalian paused elongation complex (PDB 6GML). Positions of IWS1 and ELOF1 do not clash with NELF.
