## Supplementary material for "Structure and function of IWS1 in transcription elongation": Table 1 and 2

**Table 1. Cryo-EM data collection, refinement and validation statistics**

| | #1 composite map T RNA polymerase II-DSIF-SPT6-PAF1c-TFIIS-IWS1-ELOF1 | #2 activated elongation complex (consensus) map A | #3 IWS1+ELOF1 map B | #4 IWS1 C-term map C | #5 TFIIS map D |
|---|---|---|---|---|---|
| **Data collection and processing** | | | | | |
| Magnification | 105,000 | 105,000 | 105,000 | 105,000 | 105,000 |
| Voltage (kV) | 300 | 300 | 300 | 300 | 300 |
| Electron exposure (e–/Å$^2$) | 37 | 37 | 37 | 37 | 37 |
| Defocus range (μm) | -1.8 to -0.6 | -1.8 to -0.6 | -1.8 to -0.6 | -1.8 to -0.6 | -1.8 to -0.6 |
| Pixel size (Å) | 1.19 | 1.19 | 1.19 | 1.19 | 1.19 |
| Symmetry imposed | C1 | C1 | C1 | C1 | C1 |
| Initial particle images (no.) | 4,597,368 | 4,597,368 | 762,523 | 762,523 | 762,523 |
| Final particle images (no.) | 762,523 | 762,523 | 378,540 | 156,410 | 36,198 |
| Map resolution (Å) | 2.4 | 2.4 | 2.6 | 2.4 | 4.3 |
| FSC threshold | 0.143 | 0.143 | 0.143 | 0.143 | 0.143 |
| Map resolution range (Å) | 2.4-10 | 2.4-10 | 2.6-10 | 2.4-10 | 4.3-10 |
| **Refinement** | | | | | |
| Initial model used (PDB code) | 6TED,9EGZ | | | | |
| Model resolution (Å) | 2.4 | | | | |
| FSC threshold | 0.143 | | | | |
| Model resolution range (Å) | 2.4-3.1 | | | | |
| Map sharpening *B* factor (Å$^2$) | 63.2 | | | | |
| Model composition | | | | | |
| Non-hydrogen atoms | 60482 | | | | |
| Protein residues | 8304 | | | | |
| Ligands | 11 | | | | |
| *B* factors (Å$^2$) | | | | | |
| Protein | 97.29 | | | | |
| Ligand | 149.34 | | | | |
| R.m.s. deviations | | | | | |
| Bond lengths (Å) | 0.007 | | | | |
| Bond angles (°) | 0.813 | | | | |
| Validation | | | | | |
| MolProbity score | 2.27 | | | | |
| Clashscore | 12.82 | | | | |
| Poor rotamers (%) | 2.31% | | | | |
| Ramachandran plot | | | | | |
| Favored (%) | 94.47% | | | | |
| Allowed (%) | 5.38% | | | | |
| Disallowed (%) | 0.15% | | | | |

| | #6 DSIF map E | #7 RTF1 map F | #8 10-subunit Pol II map G | #9 Non-template DNA map H | #10 RPB5+DNA map I |
|---|---|---|---|---|---|
| **Data collection and processing** | | | | | |
| Magnification | 105,000 | 105,000 | 105,000 | 105,000 | 105,000 |
| Voltage (kV) | 300 | 300 | 300 | 300 | 300 |
| Electron exposure (e–/Å$^2$) | 37 | 37 | 37 | 37 | 37 |
| Defocus range (μm) | -1.8 to -0.6 | -1.8 to -0.6 | -1.8 to -0.6 | -1.8 to -0.6 | -1.8 to -0.6 |
| Pixel size (Å) | 1.19 | 1.19 | 1.19 | 1.19 | 1.19 |
| Symmetry imposed | C1 | C1 | C1 | C1 | C1 |
| Initial particle images (no.) | 762,563 | 762,523 | 762,523 | 762,523 | 762,523 |
| Final particle images (no.) | 79,487 | 27,130 | 115,139 | 113,665 | 59,649 |
| Map resolution (Å) | 2.7 | 2.8 | 2.5 | 2.6 | 2.7 |
| FSC threshold | 0.143 | 0.143 | 0.143 | 0.143 | 0.143 |
| Map resolution range (Å) | 2.7-10 | 2.8-10 | 2.5-10 | 2.6-10 | 2.7-10 |
| **Refinement** | | | | | |
| Initial model used (PDB code) | | | | | |
| Model resolution (Å) | | | | | |
| FSC threshold | | | | | |
| Model resolution range (Å) | | | | | |
| Map sharpening *B* factor (Å$^2$) | | | | | |
| Model composition | | | | | |
| Non-hydrogen atoms | | | | | |
| Protein residues | | | | | |
| Ligands | | | | | |
| *B* factors (Å$^2$) | | | | | |
| Protein | | | | | |
| Ligand | | | | | |
| R.m.s. deviations | | | | | |
| Bond lengths (Å) | | | | | |
| Bond angles (°) | | | | | |
| Validation | | | | | |
| MolProbity score | | | | | |
| Clashscore | | | | | |
| Poor rotamers (%) | | | | | |
| Ramachandran plot | | | | | |
| Favored (%) | | | | | |
| Allowed (%) | | | | | |
| Disallowed (%) | | | | | |

| | #11 CDC73 map J | #12 CTR9+WDR61 map K | #13 CTR9 TPR map L | #14 PAF1+LEO1 map M | #15 LEO1 C-term map N |
|---|---|---|---|---|---|
| **Data collection and processing** | | | | | |
| Magnification | 105,000 | 105,000 | 105,000 | 105,000 | 105,000 |
| Voltage (kV) | 300 | 300 | 300 | 300 | 300 |
| Electron exposure (e–/Å$^2$) | 37 | 37 | 37 | 37 | 37 |
| Defocus range (μm) | -1.8 to -0.6 | -1.8 to -0.6 | -1.8 to -0.6 | -1.8 to -0.6 | -1.8 to -0.6 |
| Pixel size (Å) | 1.19 | 1.19 | 1.19 | 1.19 | 1.19 |
| Symmetry imposed | C1 | C1 | C1 | C1 | C1 |
| Initial particle images (no.) | 762,563 | 762,523 | 762,523 | 762,523 | 762,523 |
| Final particle images (no.) | 43,339 | 63,837 | 63,837 | 28,545 | 6,771 |
| Map resolution (Å) | 3.4 | 3.8 | 2.6 | 3.2 | 4.0 |
| FSC threshold | 0.143 | 0.143 | 0.143 | 0.143 | 0.143 |
| Map resolution range (Å) | 3.4-10 | 3.8-10 | 2.6-10 | 3.2-10 | 4.0-10 |
| **Refinement** | | | | | |
| Initial model used (PDB code) | | | | | |
| Model resolution (Å) | | | | | |
| FSC threshold | | | | | |
| Model resolution range (Å) | | | | | |
| Map sharpening *B* factor (Å$^2$) | | | | | |
| Model composition | | | | | |
| Non-hydrogen atoms | | | | | |
| Protein residues | | | | | |
| Ligands | | | | | |
| *B* factors (Å$^2$) | | | | | |
| Protein | | | | | |
| Ligand | | | | | |
| R.m.s. deviations | | | | | |
| Bond lengths (Å) | | | | | |
| Bond angles (°) | | | | | |
| Validation | | | | | |
| MolProbity score | | | | | |
| Clashscore | | | | | |
| Poor rotamers (%) | | | | | |
| Ramachandran plot | | | | | |
| Favored (%) | | | | | |
| Allowed (%) | | | | | |
| Disallowed (%) | | | | | |

| | #16 SPT6 map O | #17 SPT6 N-term map P | #18 LEDGF+ nucleosome map Q | #19 Nucleosome placement map R | #20 Local 10-subunit Pol II map S |
|---|---|---|---|---|---|
| **Data collection and processing** | | | | | |
| Magnification | 105,000 | 105,000 | 105,000 | 105,000 | 105,000 |
| Voltage (kV) | 300 | 300 | 300 | 300 | 300 |
| Electron exposure (e–/Å$^2$) | 37 | 37 | 37 | 37 | 37 |
| Defocus range (μm) | -1.8 to -0.6 | -1.8 to -0.6 | -1.8 to -0.6 | -1.8 to -0.6 | -1.8 to -0.6 |
| Pixel size (Å) | 1.19 | 1.19 | 1.19 | 1.19 | 1.19 |
| Symmetry imposed | C1 | C1 | C1 | C1 | C1 |
| Initial particle images (no.) | 762,523 | 762,523 | 762,523 | 19,562 | 19,562 |
| Final particle images (no.) | 90,114 | 90,114 | 19,562 | 19,562 | 19,562 |
| Map resolution (Å) | 2.7 | 2.9 | 3.5 | 7.6 | 8.7 |
| FSC threshold | 0.143 | 0.143 | 0.143 | 0.143 | 0.143 |
| Map resolution range (Å) | 2.7-10 | 2.9-10 | 3.5-10 | 7.6-20 | 8.7-20 |
| **Refinement** | | | | | |
| Initial model used (PDB code) | | | 6S01 | | |
| Model resolution (Å) | | | 3.5 | | |
| FSC threshold | | | 0.143 | | |
| Model resolution range (Å) | | | 3.5-4.2 | | |
| Map sharpening *B* factor (Å$^2$) | | | 48.0 | | |
| Model composition | | | | | |
| Non-hydrogen atoms | | | 12505 | | |
| Protein residues | | | 853 | | |
| Ligands | | | 0 | | |
| *B* factors (Å$^2$) | | | | | |
| Protein | | | 144.86 | | |
| Ligand | | | - | | |
| R.m.s. deviations | | | | | |
| Bond lengths (Å) | | | 0.007 | | |
| Bond angles (°) | | | 0.980 | | |
| Validation | | | | | |
| MolProbity score | | | 2.14 | | |
| Clashscore | | | 15.50 | | |
| Poor rotamers (%) | | | 1.83 | | |
| Ramachandran plot | | | | | |
| Favored (%) | | | 96.27 | | |
| Allowed (%) | | | 3.73 | | |
| Disallowed (%) | | | 0.00 | | |

| | #21 RECQL5-RNA polymerase II map U |
|---|---|
| **Data collection and processing** | |
| Magnification | 36,000 |
| Voltage (kV) | 200 |
| Electron exposure (e–/Å$^2$) | 52.11 |
| Defocus range (μm) | 1 to 2.2 |
| Pixel size (Å) | 1.1 |
| Symmetry imposed | C1 |
| Initial particle images (no.) | 52,539 |
| Final particle images (no.) | 15,305 |
| Map resolution (Å) | 3.1 |
| FSC threshold | 0.143 |
| Map resolution range (Å) | 3.1-10 |
| **Refinement** | |
| Initial model used (PDB code) | |
| Model resolution (Å) | |
| FSC threshold | |
| Model resolution range (Å) | |
| Map sharpening *B* factor (Å$^2$) | |
| Model composition | |
| Non-hydrogen atoms | |
| Protein residues | |
| Ligands | |
| *B* factors (Å$^2$) | |
| Protein | |
| Ligand | |
| R.m.s. deviations | |
| Bond lengths (Å) | |
| Bond angles (°) | |
| Validation | |
| MolProbity score | |
| Clashscore | |
| Poor rotamers (%) | |
| Ramachandran plot | |
| Favored (%) | |
| Allowed (%) | |
| Disallowed (%) | |

**Table 2. Input structural models and model confidence**

| Complex/domain | Chain id(s) | Input model | Level of confidence |
|---|---|---|---|
| RNA Polymerase II | A-L | 6TED | Atomic |
| SPT6 | M | 6TED, Alphafold | Atomic (residues 207-218, 266-331, 340-425, 431-482, 512-695, 705-758, 809-814, 883-1074, 1136-1171), Rigid body fitting (residues 219-230, 332-339, 426-430, 483-488, 696-704, 759-763, 775-808, 815-882, 1075-1135, 1172-1175, 1227-1322,1323-1514) |
| DNA | N,T | 6TED, 6S01, *de novo* | Atomic except for single-stranded non-template DNA (residues -18--8) on 9MLC |
| ELOF1 | O | Alphafold | Atomic |
| RNA | P | 6TED, *de novo* | Atomic |
| CTR9 | Q | 6TED | Atomic (residues 440-810), Rigid body fitting (residues (3-439, 811-892) |
| RTF1 | R | 6TED | Atomic (residues 503-596), Rigid body fitting (residues 353-502, 597-624) |
| TFIIS | S | 9EGZ | Atomic (residues 132-249), Rigid body fitting (residues 250-301) |
| LEO1 | U | Alphafold | Atomic (residues 513-528), Rigid body fitting (residues 356-512, 529-555) |
| PAF1 | V | Alphafold | Atomic (residues 120-149), Rigid body fitting (residues 45-113, 151-371) |
| WDR61 | W | Alphafold | Atomic |
| CDC73 | X | 6TED, Alphafold | Atomic (residues 218-260), Rigid body fitting (residues 360-531) |
| SPT4 | Y | 6TED | Atomic |
| SPT5 | Z | 6TED | Atomic (residues 160-269, 417-523, 702-752), Rigid body fitting (residues 270-397, 524-645, 753-781) |
| H3 | a,e | 6S01 | Atomic |
| H4 | b,f | 6S01 | Atomic |
| H2A | d,h | 6S01 | Atomic |
| H2B | c,g | 6S01 | Atomic |
| IWS1 | i | 9EGZ, Alphafold | Atomic (residues 554-711, 749-770, 787-795, 805-819), Rigid body fitting (residues 528-542, 712-721, 787-795) |
| LEDGF PWWP | l | 6S01 | Rigid body fitting |
